# Splicing of HPV16 E6 promotes aggressive invasion in oropharyngeal cancer via redistribution of E-cadherin

**DOI:** 10.1101/2025.10.16.682706

**Authors:** Yvonne X. Lim, Min Liu, Bailey F. Garb, Allison Furgal, Shiting Li, Shaomiao Xia, Andrew Whiteman, Junsouk Choi, Qingzhi Liu, Huira Kopera, Roland Hilgarth, Marcell Costa de Medeiros, Laura Gonzalez-Maldonado, Thomas Lanigan, Jonathan McHugh, Gregory Wolf, Michelle L. Mierzwa, Veerabhadran Baladandayuthapani, Laura S. Rozek, Maureen A. Sartor, Nisha J. D’Silva

**Affiliations:** Department of Periodontics and Oral Medicine, University of Michigan School of Dentistry, 1011 N. University Ave, Ann Arbor, Michigan, USA; Rogel Cancer Center, University of Michigan, 1500 E Medical Center Dr, Ann Arbor; Department of Biostatistics, University of Michigan School of Public Health, Ann Arbor; Department of Computational Medicine and Bioinformatics, University of Michigan Medical School, Ann Arbor; Department of Human Genetics, University of Michigan Medical School, Ann Arbor; Vector Core, Biomedical Research Core Facilities, University of Michigan Medical School, Ann Arbor; Department of Internal Medicine, University of Michigan Medical School, Ann Arbor; Department of Pathology, University of Michigan Medical School, Ann Arbor; Department of Otolaryngology, University of Michigan Medical School, Ann Arbor; Department of Radiation Oncology, University of Michigan Medical School, Ann Arbor; Department of Oncology, School of Medicine, Georgetown University, Washington D.C

## Abstract

Human papillomavirus-positive oropharyngeal squamous cell carcinoma (HPV+ OPSCC) has become the most common HPV-associated cancer in developed countries, yet a significant subset of patients develops recurrence despite favorable treatment responses. Biomarkers that identify aggressive tumors at diagnosis are therefore needed. In HPV+ cancers, the viral E6 transcript can be expressed as two main variants, the full-length isoform (E6FL) and the spliced E6*I isoform. Here, we investigated the functional and clinical impact of E6FL and E6*I in HPV+ OPSCC. Across multiple preclinical models, E6*I promoted migration and aggressive invasion relative to E6FL and was associated with redistribution of E-cadherin away from the cell membrane. In multi-institutional patient cohorts, a previously developed E6FL:E6_ALL_ influence score, which captures E6 splicing-associated host transcriptional biology, was associated with clinical outcomes. Lower scores, reflecting greater E6*I impact, were associated with worse overall survival and recurrence-free survival. Multiplexed immunofluorescence of 75 primary pretreatment biopsies further showed that a lower influence score was linked to reduced membrane and increased cytoplasmic localization of E-cadherin, while a tissue-based E-cadherin localization score was associated with recurrence. Mechanistically, E6*I overexpression promotes E-cadherin internalization, likely through the Rab11-associated trafficking pathway. Splice-switching oligonucleotides that shifted endogenous E6 splicing toward E6FL reduced invasion in two HPV+ OPSCC models. Together, these findings position HPV E6 splicing as a regulatory axis underlying aggressive HPV+ OPSCC biology and support E6 splicing and membrane-to-cytoplasmic ECAD localization as candidate risk-stratification biomarkers.

## Introduction

Oropharyngeal squamous cell carcinoma (OPSCC), a subtype of head and neck SCC (HNSCC), has surpassed cervical cancer to become the most common HPV-positive (HPV+) malignancy in the United States and other developed countries(*1, 2*). While standard chemoradiation achieves high cure rates, treatment-related toxicities such as dysphagia and xerostomia substantially diminish quality of life(*3, 4*). De-intensification efforts have not yet been established as a standard approach for unselected patients, and some of these approaches have resulted in inferior tumor control(*2, 5–7*). Moreover, 25-30% of patients experience tumor recurrence or persistence following standard therapy(*8–10*), even patients with early stage (I/II) disease(*11, 12*). The lack of robust pre-treatment biomarkers for risk stratification and treatment selection remains a major challenge; current candidates have not reliably predicted aggressive disease at diagnosis, limiting clinical utility(*2, 4*).

Unlike other HNSCC, HPV+ OPSCC harbors few actionable genetic alterations(*13, 14*), underscoring the importance of exploring alternative mechanisms such as splicing regulation(*15, 16*). HPV16 is the predominant genotype driving OPSCC, with its viral oncoproteins E6 and E7 playing central roles in cancer development(*4*). Besides disrupting host cell cycle regulation and differentiation, E6 and E7 also influence immune evasion, genomic instability, and epithelial to mesenchymal transition (EMT) to drive tumor progression(*17*). Importantly, full-length HPV16 E6 (E6FL) transcript undergoes splicing to generate truncated variants known as E6*(*18*). Among all E6 spliced variants, E6*I, resulting from exclusion of 183 bp from E6FL, is the most abundant E6 splice variant in OPSCC(*19–23*). Despite its abundance, the impact of differential E6 splicing on biology and clinical outcomes in OPSCC remains poorly studied. Previous studies investigating E6 splice variants have been limited and relied on models of cervical cancer or other cancers(*24–28*) that differ significantly from OPSCC both clinically and biologically(*2, 29*). Our earlier bioinformatics study in 68 HPV+ HNSCC patients (>75% OPSCC) suggests that a host transcriptional program associated with higher spliced E6* relative to E6FL transcript composition may predict poorer overall survival and larger tumors(*30*), implicating E6 splicing as a potential regulatory switch for aggressive HPV+ OPSCC.

In HPV+ OPSCC, invasion of tumor cells into adjacent tissues and regional lymph nodes has been linked to locoregional and distant recurrences(*31–34*). However, the molecular mechanisms underlying invasion are poorly characterized. EMT, a driver of tumor invasion, was once viewed as a binary process where epithelial cells lost cell-cell adhesion and apical-basal polarity while acquiring mesenchymal characteristics. Recent studies highlighted the significance of intermediate states known as partial EMT (p-EMT)(*35, 36*). Tumor cells in p-EMT states retain some epithelial markers and tumor-initiating capabilities but gain stemness and motility(*36–38*). A key mechanism driving p-EMT is the relocalization of epithelial proteins away from cell-cell junctions to reduce cell-cell adhesion(*39, 40*). In oral cavity SCC, p-EMT promotes collective migration and is strongly correlated with nodal metastasis and invasion(*41*).

Significant heterogeneity in p-EMT states has also been identified in HPV+ OPSCC(*42*), but its molecular drivers and prognostic significance remain unclear.

In this study, we explored the impact of HPV16 E6 splicing on OPSCC progression by integrating multi-institutional patient cohorts, disease-relevant functional models, and mechanistic studies. Our findings support a model in which a shift of splicing balance from HPV E6FL toward E6*I promotes aggressive invasion and recurrence via p-EMT, characterized by the cytoplasmic relocalization of cell-cell junction epithelial protein, E-cadherin (ECAD). We established an ECAD localization score to quantify the membrane and cytoplasmic distribution of ECAD in patients’ tumor tissues. We developed antisense splice-switching oligonucleotides (SSOs) to inhibit splicing of E6FL to E6*I, thereby attenuating invasive phenotypes. Taken together, our findings establish HPV16 E6 splicing as a mechanistic driver for aggressive OPSCC and provide biological rationale for nominating E6 splicing and ECAD cytoplasmic localization as potential biomarkers for risk stratification in future studies.

## Results

### 1. HPV16+ OPSCC cell lines express varying proportions of E6FL and E6*I

In functional and mechanistic studies, we focused on HPV16, which accounts for more than 80% of HPV+ OPSCC(*43*). HPV16 E6FL is primarily spliced into E6*I and less abundantly to E6*II (*18, 20, 44*) (Fig. 1A). Recent studies in human HPV+ head and neck squamous cell carcinoma (HNSCC) have shown that E6*I is the most abundant among all E6 transcript variants, followed by E6FL and E6*II (*19, 21, 22, 45*). However, the functional and mechanistic impact of E6 spliced variants have not been systematically characterized in pre-clinical models of OPSCC.

**Fig. 1:**
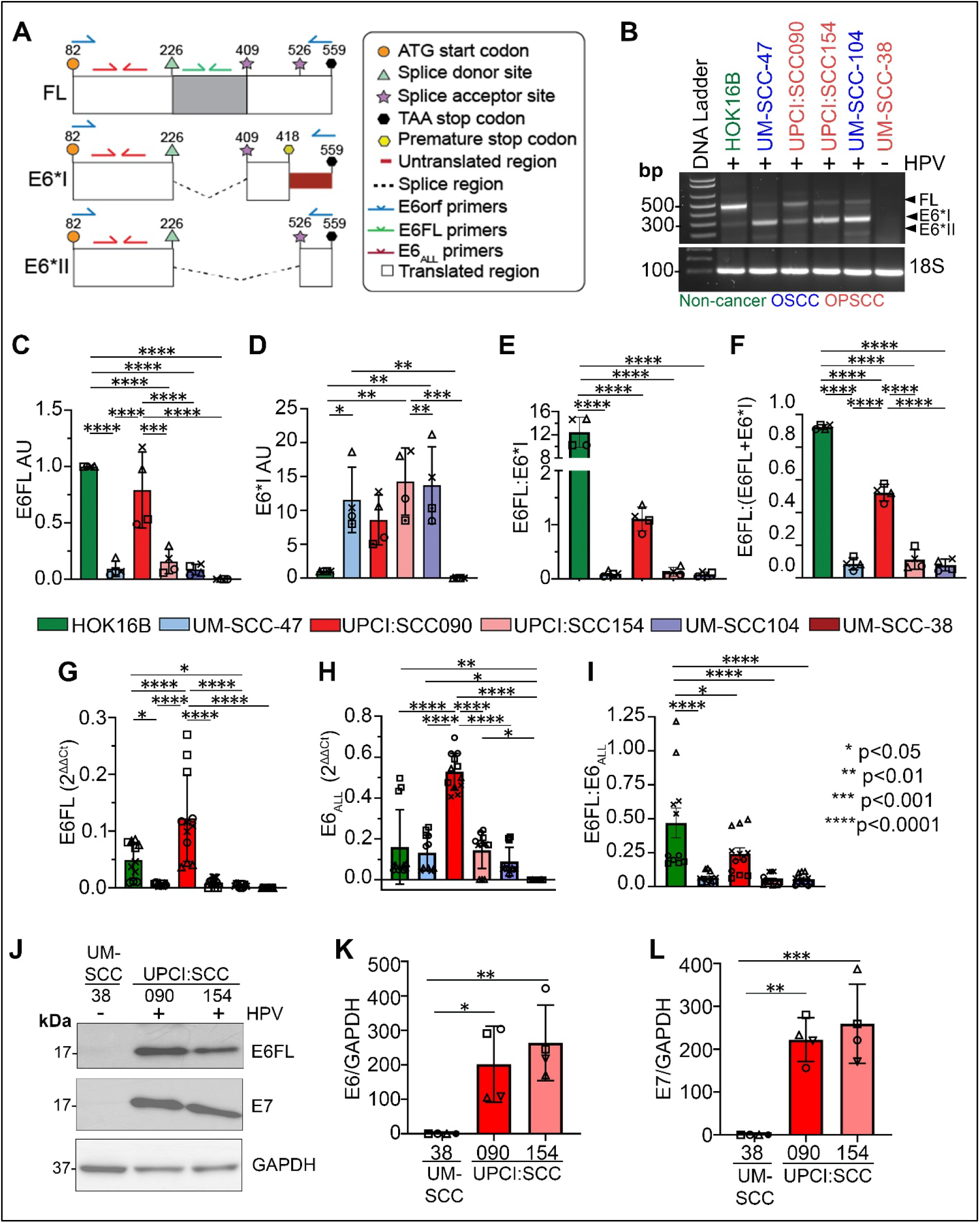
HPV16 E6*I and E6FL transcripts are differentially expressed in HPV+ cell lines. (A) Schematic indicating primers used to detect E6FL, E6*I and E6*II. (B) Representative agarose gel from 4 independent experiments detecting E6 variants in cDNA from a panel of oral cavity SCC, OPSCC, and non-cancer (keratinocyte) cell lines, using primers against E6 open reading frame (E6orf). Expected sizes for E6FL, E6*I and E6*II are 477 bp, 295 bp and 178 bp respectively. (C-F) Quantification of four independent experiments (including (B)) by densitometric analysis of E6FL and E6*I normalized to 18S (AU = arbitrary units). Each shape shows an independent experiment. (G-I) RT-qPCR of E6FL and all E6 variants (E6_ALL_) normalized to β actin. Each shape represents an independent experiment; three points of the same shape indicate three replicates. (J-L) Immunoblot of HPV E6FL and E7. GAPDH is used as a control (J). Densitometries of E6FL (K) and E7 (L) was normalized to GAPDH. For all graphs, data are represented as mean ± SD. *p<0.05, **p<0.01, ***p<0.001, ****p<0.0001 (one-way ANOVA post-hoc Tukey test)

To establish disease-relevant pre-clinical models for interrogating the impact of differential E6 splice variant composition, we first examined the expression of E6 spliced variants across a panel of HPV16+ OPSCC and oral cavity squamous cell carcinoma (OSCC) cell lines using complementary reverse transcription-polymerase chain reaction approaches. In the first approach, primers spanning the open reading frame of the E6 (E6orf) gene were used to amplify all E6 splice variants (Fig. 1A; blue primers). Agarose gel electrophoresis resolved distinct bands corresponding to E6FL and E6*I according to their expected molecular sizes, with only a faint band corresponding to E6*II (Fig. 1B). This is consistent with previous studies reporting that E6FL and E6*I transcripts are the most abundance splice variants in OPSCC (*19, 21–23*). Quantitative analysis revealed that HPV+ cancer cell lines had lower E6FL and higher E6*I transcript expression than HOK16B, an HPV-immortalized non-cancer keratinocyte cell line (Fig. 1C, 1D). As expected, the HPV(-) OPSCC cell line, UM-SCC-38, did not express any E6 variant and served as a negative control (Fig. 1B-1D).

Because absolute transcript abundance does not directly reflect relative splice composition, we calculated: (i) ratio of unspliced E6FL to spliced E6*I (Fig. 1E); and (ii) ratio of E6FL to the total of the two main variants, E6FL and E6*I (Fig. 1F). E6*II was excluded due to negligible transcript expression. HPV+ OPSCC and OSCC cell lines have lower E6FL:E6*I or E6FL:(E6FL+E6*I) ratios compared to non-cancer, HPV-immortalized HOK16B, consistent with increased relative splicing of E6FL toward E6*I(Fig. 1E, 1F).

We independently validated the above RT-PCR results with a complementary approach using isoform-specific primers that selectively amplify E6FL or total E6 (E6_ALL_) transcripts (Fig. 1A). Using this approach, HPV+ cancer cell lines again express varying proportions of E6FL and E6_ALL_ with significant variation in the E6FL:E6_ALL_ ratio (Fig. 1G-1H). Notably, the two HPV+ OPSCC cell lines show different splicing profiles, with UPCI:SCC154 having significantly lower E6FL mRNA expression and lower E6FL:E6_ALL_ ratio than UPCI:SCC090 (Fig. 1E, 1F, 1I). As expected, UM-SCC-38, the HPV(-) OPSCC cell line, does not express either variant.

Immunoblotting detected a band at the expected molecular weight of E6FL, approximately 17 kDa, in UPCI:SCC090 and UPCI:SCC154, but not in HPV-negative UM-SCC-38 cells (Fig. 1J–K). A band corresponding to the expected size of E6*I was not detected. E7 was also detected in both HPV16+ OPSCC cell lines but not in UM-SCC-38 cells (Fig. 1J, 1L). Although UPCI:SCC154 had substantially less E6FL transcript than UPCI:SCC090, the expression of the E6FL protein band did not differ significantly between the two cell lines.

Together our findings indicate that E6*I expression is higher in HPV+ cancer cell lines than in HOK16B, HPV-immortalized non-cancer cells. Also, HPV+ cell lines express varying proportions of E6FL and E6*I transcripts, and E6*II expression is minimal. Because E6*I protein could not be detected by immunoblotting, subsequent experiments evaluate composition of E6 splice transcript variants and associated phenotypes rather than E6*I protein expression.

### 2. Endogenous variations in E6 spliced transcript ratio are associated with migration and ECAD redistribution in HPV+ OPSCC cells

Since spliced E6*I is the most abundant E6 isoform in HPV+ cell lines (Fig. 1B), and in HPV+ OPSCC from patients (*19, 21, 45*), we investigated whether the two main E6 isoforms, E6FL and E6*I, differentially contribute to oncogenic phenotypes in UPCI:SCC154, an HPV+ OPSCC cell line with endogenous E6 and E7. To do this, we leveraged the cellular heterogeneity found in parental HPV+ OPSCC cell lines (*42*) and isolated 29 single-cell clones from the HPV+ UPCI:SCC154 cell line with varying E6FL:E6*I transcript ratios (Fig. 2A, 2B). Low E6FL:E6*I transcript ratio indicates increased splicing of E6FL to E6*I, and vice versa. Across 4 consecutive passages, the 29 clones had a wide range of E6FL:E6*I transcript ratios, although in later passages, E6FL:E6*I was progressively more homogeneous between some clones (Fig. 2B). To minimize passage-dependent effects, all functional studies were performed only within the first four passages, focusing on clones with consistently lower (#3, 6, 11) or higher (#24, 25, 27) E6FL:E6*I transcript ratio (Fig. 2B; Fig. S1A). To assess whether E6FL:E6*I ratio correlates with total E6 transcript expression (E6_ALL_), we quantified the total expression of E6FL and E6*I transcripts. E6_ALL_ mRNA levels did not differ between the clones with high and low ratio (Fig. S1B). Therefore, changes in oncogenic phenotypes between the clones are unlikely due to changes in total E6 transcript expression.

**Fig. 2:**
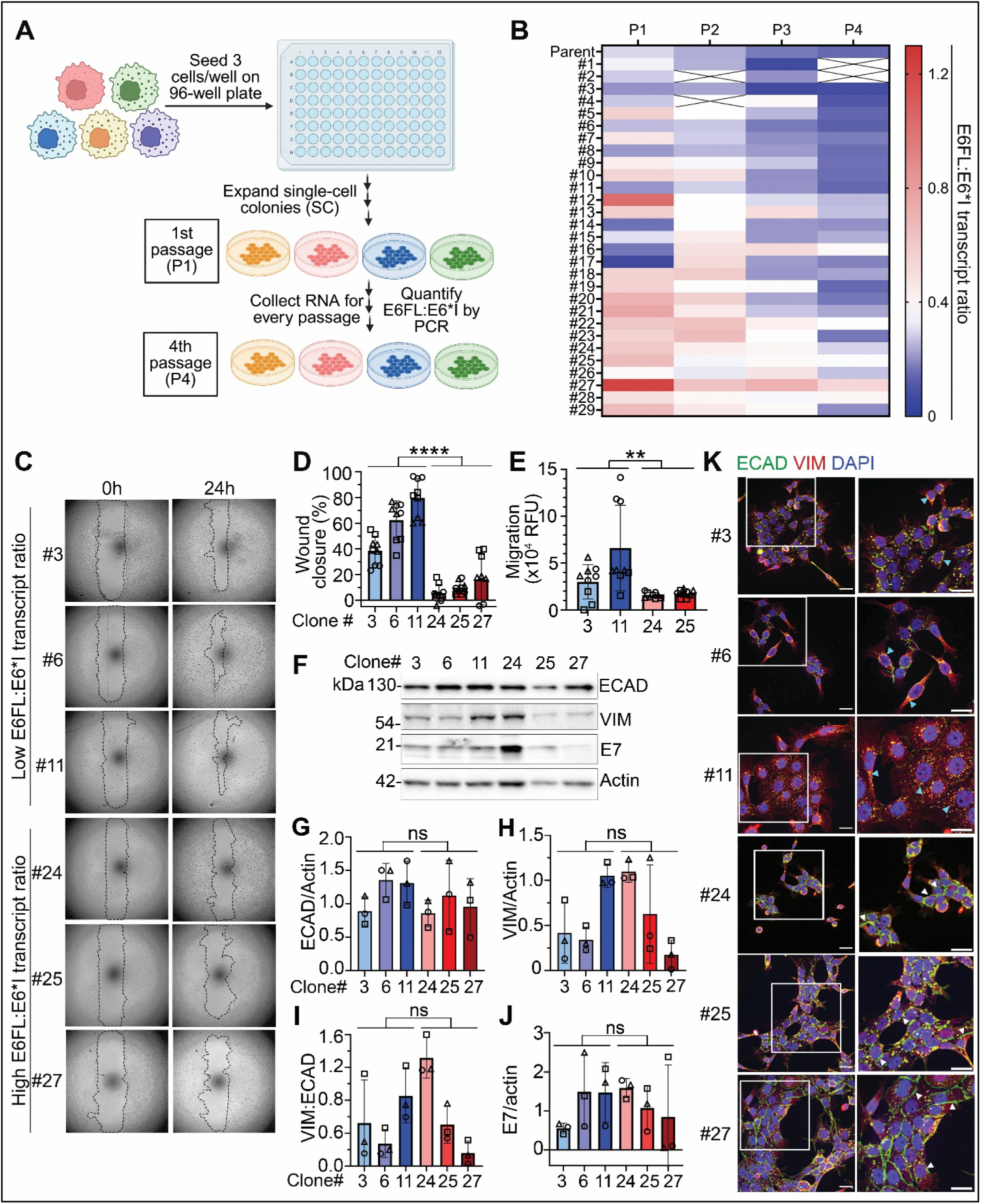
Low E6FL:E6*I transcript ratio is associated with increased migration in HPV+ OPSCC UPCI:SCC154 single-cell clones. (A) Schematic illustrating single-cell (SC) clone isolation and expansion. Created with Biorender.com. (B) Heat map showing the values of E6FL:E6*I for each clone quantified from RT-PCR. X indicates undetectable expression of E6FL or E6*I. (C) Representative images for wound healing assays from three independent experiments. (D) Migration is quantified from 3 independent experiments including (C) and represents percent (%) wound closure after 24 h. (E) Fluoroblok transwell migration assay. Migration was calculated by normalizing with relative fluorescence unit (RFU) at 0h. For data in D&E, each shape represents an independent experiment (n=3), with three replicates per experiment. ** p<0.01, **** p<0.0001 (one-way ANOVA post-hoc Tukey) (F) Representative immunoblot of ECAD and VIM in UPCI:SCC154SC clones from three independent experiments. Actin was used as a loading control. (G-J) Densitometry of ECAD (G) and VIM (H), were normalized to actin. VIM:ECAD (I) was quantified to assess mesenchymal phenotype. Densitometry of E7 (J) was normalized to actin. Each shape represents an independent experiment (n=3); ns= non-significant; ** p<0.01, ****p<0.0001 (unpaired t test). (K) Immunofluorescence staining of DAPI (blue), ECAD (green) and VIM (red) in clones. Right panel shows enlarged images from boxed area in left panel Scale bar = 100um (both left and right). Cyan arrowheads highlight examples of cytoplasmic ECAD; white arrowheads show examples of membrane ECAD. Quantification is provided in Fig. S1I-S1K

Clones with low E6FL:E6*I transcript ratio (higher splicing to E6*I) had significantly more wound closure and proliferation than clones with high E6FL:E6*I (Fig. 2C,2D; S1C). This finding was independently validated using a complementary transwell migration assay, in which we randomly selected two clones with low E6FL:E6*I and two clones with high E6FL:E6*I (Fig. 2E). Consistent with these functional differences, colonies from #3 and #11 (low E6FL:E6*I; higher splicing to E6*I) exhibited a more diffuse and scattered growth pattern that indicates a more mesenchymal phenotype, whereas colonies from #24, #25 and #27 (high E6FL:E6*I), remained compact and epithelial-like (Fig. S1D), suggesting altered cell-cell adhesion and motility.

To determine whether these phenotypic differences reflect EMT, we assessed protein expression of epithelial (E-cadherin, ECAD) and mesenchymal (vimentin, VIM) markers. Immunoblot analysis revealed no significant differences in total ECAD or VIM levels between low- and high-ratio clones (Figs. 2F-2H), nor in the VIM:ECAD ratio commonly used to assess EMT(*46*) (Fig. 2I). Thus, the phenotypic differences were not accompanied by a conventional complete EMT pattern characterized by reduced total ECAD and increased VIM.

Although E6*I may enhance translation of E7(*47, 48*), other studies reported conflicting findings(*49, 50*). Therefore, we investigated if altered E7 expression or activity accounts for the enhanced migration in clones with low E6FL:E6*I(3, 6, 11) compared to those with higher ratios (#24, 25, 27). Immunoblot analysis showed no significant differences in E7 protein expression between clones with low and high E6FL:E6*I transcript ratios (Fig. 2F, 2J). Since E7 degrades retinoblastoma (Rb) and enhances p16 expression, we also investigated Rb and p16 to assess E7 activity (Fig. S1E). Protein expression of Rb and p16, both downstream targets of E7, also did not differ between clones with low and high E6FL:E6*I transcript ratios (S1E-S1G). Therefore, changes in migration associated with E6FL:E6*I transcript ratios are unlikely due to changes in E7 activity.

We then examined the subcellular localization of ECAD and VIM. Immunofluorescence analysis showed that SC clones #3, #6, #11 (low E6FL:E6*I) demonstrated less membrane localization and higher cytoplasmic localization of ECAD than #24, #25, #27 (high E6FL:E6*I) (Fig. 2K; Fig. S1H, S1I), although total ECAD and VIM expression did not differ significantly between the clones (Fig. S1J, S1K). Therefore, lower E6FL:E6*I was associated with redistribution of ECAD away from the cell-cell junctions, consistent with a p-EMT phenotype (*39, 51*).

Together, these findings associate lower E6FL:E6*I transcript ratio with greater proliferation and migration, a more dispersed cellular morphology, and reduced membrane ECAD localization in HPV+ UPCI:SCC154 cells. Because the single-cell clone analysis establishes an association but does not distinguish a causal effect of E6 splice transcript composition from other clone-specific differences, we next experimentally modulated endogenous E6 splicing using splice-switching oligonucleotides (SSOs) in two HPV16+ OPSCC models

### 3. SSO-mediated modulation of HPV16 E6 splicing alters invasion and ECAD localization

To determine whether experimental modulation of endogenous E6 splicing alters the phenotypes associated with the E6FL:E6*I transcript ratio and to control for potential clonal artifacts, we designed seven splice-switching oligonucleotides (SSOs) to inhibit this splicing (Fig. 3A; Table S1). Screening SSOs in UPCI:SCC154 identified two, SSO-1 and SSO-2, that increased E6FL:E6*I transcript ratio more than four-fold compared to non-targeting SSO (NC), suggesting that they block splicing (Fig. 3B; blue box). In contrast, SSO-5 and SSO-6 modestly reduced E6FL:E6*I compared to NC, suggesting that they enhance splicing to E6*I (Fig. 3B; red box). Similar effects on E6FL:E6*I transcript ratio were noted in a second HPV+ OPSCC cell line, UPCI:SCC090, in which SSO-1 and SSO-2 increased the ratio and SSO-5 and SSO-6 decreased the ratio relative to NC (Fig. S2A). Importantly, the total transcript expression of E6 (E6_ALL_; contains E6FL and E6*I) was similar across all SSO conditions (Fig. 3B, S2A). Therefore, the downstream effects of SSOs are unlikely due to changes in total E6 transcript abundance.

**Fig. 3:**
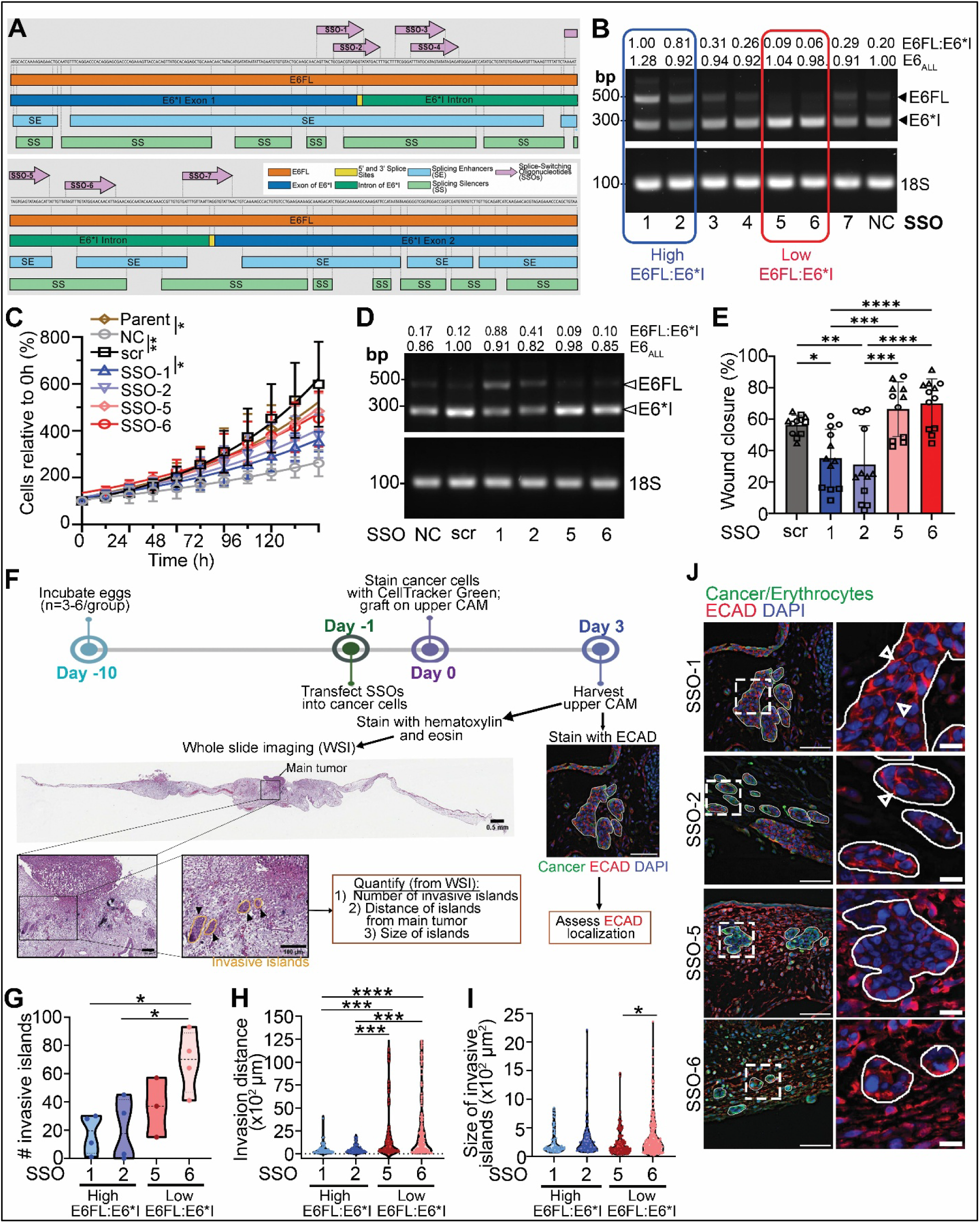
Inhibition of E6FL splicing suppresses invasion of HPV16+ OPSCC. (A) Schematic illustrating position of SSOs (purple) in HPV16 E6 pre-mRNA. (B) Representative agarose gel from 3 independent experiments showing expression of E6 isoforms in cDNA of UPCI:SCC154 transiently transfected with SSOs. E6FL:E6*I ratios and E6_ALL_ (sum of E6FL and E6*I) were quantified by densitometry. E6_ALL_ transcript was normalized with 18S and expressed as fold change from negative control (NC) SSO. Blue box represents inhibition of splicing (high E6FL:E6*I) compared to NC; red box represents the reverse. (C) Proliferation assays. Viable cells are quantified from 3 independent experiments and expressed as percent (%) of 0h. *p<0.05; **p<0.01; no asterisk= not-significant (one-way ANOVA post-hoc Tukey). (D) Agarose gel showing independent validation of effects of SSOs selected from red and blue box in (B) on E6FL:E6*I transcript ratio in UPCI:SCC154. E6_ALL_ transcript was normalized with scrambled SSO control. (E) Wound closure after 18h is expressed as a percent of wound area at 0h in UPCI:SCC154 transfected with SSOs. Each shape represents an independent experiment; each point represents one replicate. Representative images are in Fig. S2E. (F) Schematic of chick chorioallantoic membrane (CAM) experiment. Invasive islands (orange circle, black arrowheads) are identified as cancer cells that are detached from the main tumor bulk. (G-I) Quantification of invasive islands across whole-slide CAM images. Number of invasive islands (G), distance from invasive islands to the nearest tumor bulk (H), and size of invasive islands (I) were quantified from the entire CAM section. Each point in (G) represents a different egg. Each point in (H) and (I) represent an invasive tumor island. (J) Immunofluorescence staining of ECAD (red) in tumor islands (outlined in white) from upper CAM; cancer cells in green; DAPI in blue. Right panel is enlarged for boxed area in left panel. White arrowheads point to examples of membrane ECAD localization. Scale= 50μm (left); 10μm right). For all graphs, data are represented as mean ± SD. *p<0.05, **p<0.01, ***p<0.001, ****p<0.0001 (One-way ANOVA post-hoc Tukey test).

We next assessed whether SSOs affected cell growth. UPCI:SCC154 transfected with NC SSO exhibited minimal growth compared to untransfected parent cells, suggesting non-specific toxicity of NC SSO (Fig. 3C). Therefore, we generated a scrambled SSO for subsequent functional studies. The scrambled SSO did not substantially alter E6FL:E6*I transcript ratio (Fig. 3D) and showed comparable growth to untransfected parental UPCI:SCC154 cells(Fig. 3C). Interestingly, UPCI:SCC154 transfected with SSO-1, that enhances E6FL:E6*I transcript ratio (reduced splicing) were less proliferative than the scrambled control (Fig. 3C). SSO-2, which also reduced splicing, trended toward lower proliferation compared to scrambled control (Fig. 3C), although not significant. In contrast, SSO-5 and SSO-6, which enhanced splicing, did not have significant difference in cell growth compared to scrambled control (Fig. 3C).

Because the original non-targeting SSO reduced cell growth independently of E6 splicing, subsequent functional experiments used a scrambled SSO control that did not alter E6FL:E6*I or cell growth. We evaluated whether SSO-mediated changes in E6 splicing altered E6FL and E7 protein expression. In UPCI:SCC154, SSOs that increased the E6FL:E6*I transcript ratio, particularly SSO-1 and SSO-2, increased E6FL protein expression significantly compared to SSO-5, that shifted E6 transcript composition towards E6*I (Fig. S2B, S2C). E6FL protein expression in UPCI:SCC154 transfected with SSO-1 and SSO-2 is increased compared to SSO-5 and scrambled control, although the difference was insignificant (Fig. S2B, S2C). In contrast, E7 protein expression was not significantly altered between the SSO conditions (Fig. S2B, S2D). These findings are consistent with greater relative E6FL transcript and lower E6*I transcript expression in cells transfected with SSO-1 and SSO-2 compared to scrambled SSO (Fig. 3D), without a corresponding detectable change in E7 protein expression.

To determine if splicing modulation of E6 impacts aggressive phenotypes, we assessed migration and invasion in HPV16+ OPSCC cells transfected with SSOs. In a wound healing assay, SSO-1 and SSO-2 significantly decreased migration compared to scrambled control or SSOs that favor E6 splicing (SSO-5, SSO-6) in UPCI:SCC154 (Fig. 3E; S2E). This suggests that shifting splicing away from E6*I reduced migration, whereas conditions favoring E6*I splicing maintained higher migration.

In human HPV+ OPSCC, aggressive keratinizing tumors may exhibit invasive islands at the periphery of the main tumor(*34, 52, 53*). Furthermore, tumor budding is a prognostic feature in HPV+ OPSCC(*31*). Therefore, we tested whether shifting the E6 splice isoform balance toward or away from E6*I alters local invasion using the chick chorioallantoic membrane (CAM) model as described(*54*) (Fig. 3F). Because SSO-1/SSO-2 and SSO-5/SSO-6 shifted the E6FL:E6*I ratio in opposite directions, we used this system to compare invasion under high versus low E6FL:E6*I conditions (Fig. 3F; S2F, S2G). As expected, UPCI:SCC154 and UPCI:SCC090 with low E6FL:E6*I transcript ratios (SSO-5, SSO-6; increased splicing to E6*I) showed more invasive islands than tumors with high E6FL:E6*I transcript ratios (SSO-1, SSO-2; block splicing) (Fig. 3G; S2H). Notably, UPCI:SCC090 transfected with SSO-1 showed complete suppression of invasion, with no invasive islands observed, while SSO-2 induced only minimal invasion (Fig. S2H).

Since aggressive invasion in OPSCC is reflected by tumor islands that invade a great distance from the main tumor(*52, 53*), we quantified the distance of invasive islands from the main tumor in CAM. Since invasive islands were absent or minimal in UPCI:SCC090 following SSO-1 or SSO-2 transfection (Fig. S2H), quantification of the distance from the main tumor was not possible in this cell line. Therefore, distance of invasive islands from the main tumor was quantified only in UPCI:SCC154, where SSO-5 and SSO-6 produced invasive islands that were significantly further from the main tumor bulk compared to SSO-1 and SSO-2 (Fig. 3H). There was no significant difference in the size of islands between the high (SSO-1, SSO-2; block splicing) vs low (SSO-5, SSO-6; more splicing) E6FL:E6*I groups (Fig. 3I).

Our results using single-cell clones of UPCI:SCC154 suggested that cells with low E6FL:E6*I had reduced membrane localization of ECAD (Fig. 2K; S1I). We validated this finding in vivo by performing immunofluorescence on UPCI:SCC154 tumors on the CAM (Fig. 3J). As expected, UPCI:SCC154 with high E6FL:E6*I (SSO-1, SSO-2) demonstrated higher membrane and lower cytoplasmic localization of ECAD than those with lower E6FL:E6*I (SSO-5, SSO-6) (Fig. 3J).

Together, the use of independent SSOs that shifted E6FL:E6*I in opposite directions, while preserving total E6 transcript abundance, supports a splice-composition-dependent effect on migration, invasion, and ECAD localization. Across the single-cell clone and SSO models, lower E6FL:E6*I transcript ratios were consistently associated with increased migration and invasion and redistribution of ECAD from the membrane to the cytoplasm, consistent with a p-EMT phenotype. These findings suggest a novel role of E6 splice transcript composition in regulating aggressive phenotypes in HPV16+ OPSCC.

### 4. Overexpression of E6*I induces invasion and ECAD relocalization *in vitro and in vivo*

To independently isolate the effects of E6FL and E6*I variant independent of other viral genes, including E7, we generated constructs to express HPV16 E6*I or E6FL (Fig. 4A) and stably overexpressed them in UM-SCC-38, an HPV(-) OPSCC cell line.

**Fig. 4:**
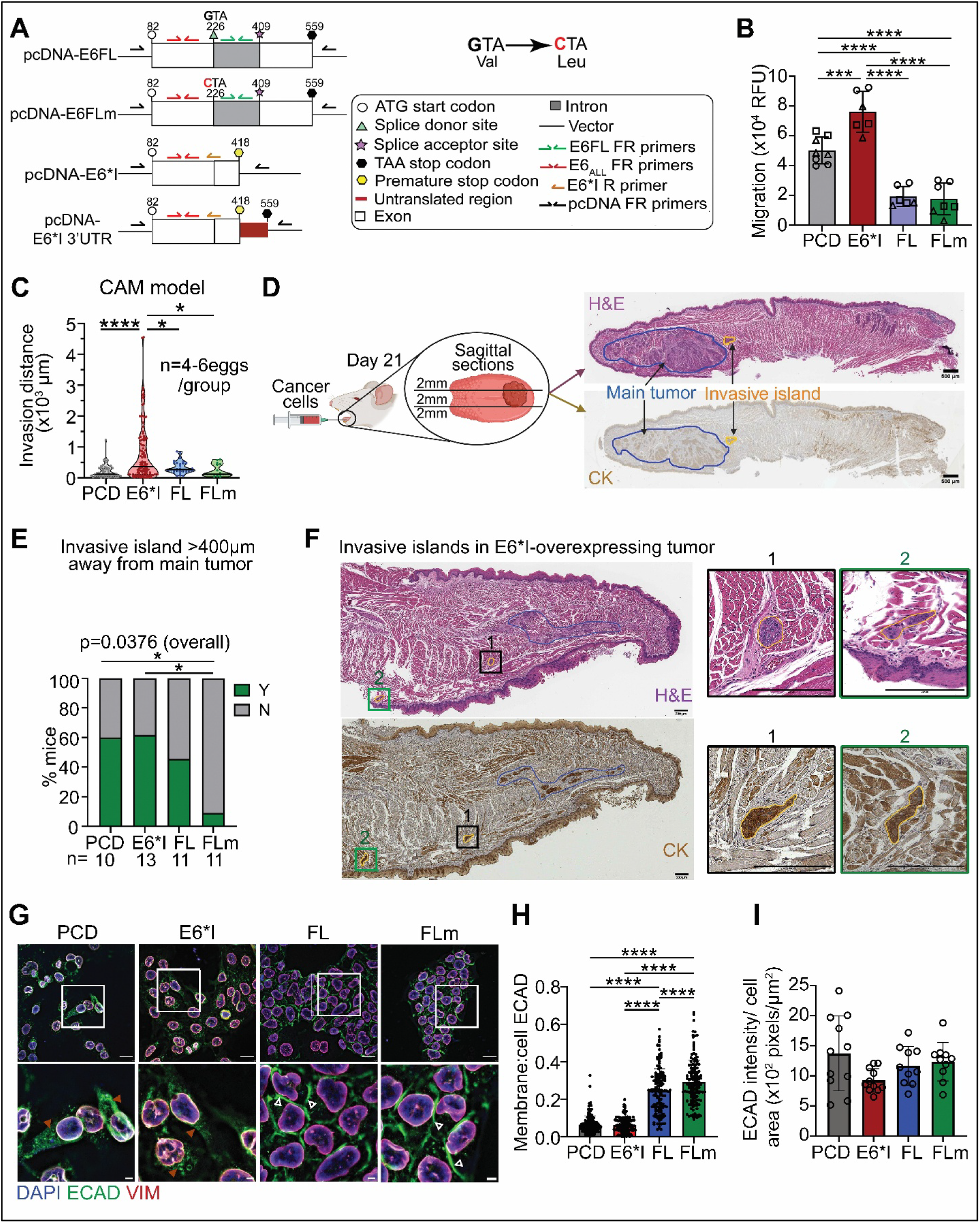
Overexpression of HPV16 E6*I promotes invasion and redistribution of E-cadherin localization away from cell membrane. UM-SCC-38 cells stably expressing empty vector or E6 variants were used. (A) Schematic of constructs expressing E6FL and E6*I. (B) Fluoroblok transwell migration assays. Migration was calculated by normalizing with relative fluorescence unit (RFU) at 0h. Each shape represents an independent experiment (n=3), identical shapes indicate replicates. (C) CAM assay. The distance from each invasive tumor island to the tumor bulk was measured in whole-slide images of the upper CAM. Each point represents an invasive tumor island; 4–6 eggs per group. (D) Orthotopic mouse tongue tumor model and representative sagittal tongue sections collected 21 days after tumor injection. Serial sections were stained with hematoxylin and eosin (H&E) or cytokeratin (CK). Blue outlines and yellow outlines indicate main tumor and invasive tumor island, respectively. (E) Percentage of mice with an invasive island >400μm from the periphery of the main tumor. (F) Representative E6*I-expressing tumor stained with H&E and CK, showing invasive tumor islands separated from the main tumor. Blue outlines indicate the main tumor, and yellow outlines indicate invasive islands. Boxed regions 1 and 2 are shown at higher magnification. Additional examples are provided in Fig. S5B. Scale bar =200µm (G) Representative ECAD (green), vimentin (VIM, red) and DAPI (blue, nucleus) immunofluorescence in cell lines. Lower panels are enlarged images from boxed area. Scale bar=100µm (upper panel) and 20µm (lower panel). Brown arrowheads indicate cytoplasmic ECAD; white arrowheads indicate membrane ECAD. (H) Quantification of membrane:cell ECAD from >120 cells in 12 fields including (G). Lower membrane:cell ECAD shows increased cytoplasmic localization of ECAD and vice versa. Each dot represents one cell. (I) Total ECAD intensity in cell areas from 10 fields per condition including G. Each dot represents one field. All data are presented as mean ± SD. For (B), (C), (H), and (I), statistical significance was assessed by one-way ANOVA post-hoc Tukey’s test. Fisher’s exact test was used in (E). ∗p<0.05, ∗∗p<0.01, ∗∗∗p<0.001, ∗∗∗∗p<0.0001.

Splicing of E6FL to E6*I results in a premature stop codon at nucleotide 418. To distinguish potential effects of transcript structure, we generated two constructs for E6*I; one terminating at the premature stop codon (termed E6*I) and the other retaining the native 3’ untranslated region (termed E6*I 3’UTR) (Fig. 4A). To prevent splicing of E6FL, we generated E6FLm by mutating G to C at nucleotide 226 (Fig. 4A), which introduces an amino acid substitution (V42L) at the splice donor site, as described(*19*).

Stable transfection into UM-SCC-38 was confirmed at genomic level using primers specific to vector (pcDNA), which allows detection of all transfected constructs, including the vector, based on molecular size (Fig. S3A). To verify isoform transcript expression, we performed reverse transcription-polymerase chain reaction with a combination of isoform-specific primers: E6FL-specific (green primers in Fig. 4A) and E6_ALL_ (red primers in Fig. 4A) that detect only E6FL or all E6 variants, respectively; and E6*I-specific primers (orange primer in Fig. 4A; E6*I reverse primer that binds to the exon-exon region to detect E6*I, not E6FL). As expected, cells overexpressing E6*I and E6*I 3’UTR showed transcripts with E6*I and E6_ALL_ primers, but none with E6FL primers (Fig. S3B-S3D). In contrast, transcripts in cells overexpressing E6FL and E6FLm were detected with both E6_ALL_ and E6FL primers (Fig. S3B-S3C). There was a low level of E6*I transcripts in cells overexpressing E6FL, but not E6FLm (Fig. S3C), showing that cells overexpressing E6FL had partial splicing, but none in E6FLm.

Since E6FL, but not E6*I, degrades p53 (*25, 55, 56*), we interrogated p53 protein expression in stably transfected cells as a functional readout for E6FL and E6*I activity. Overexpression of E6FL and E6FLm suppressed p53 in UM-SCC-38 compared to E6*I (Fig. S3E, S3F). In contrast, both E6*I and E6*I 3’UTR had similar p53 expression as the empty vector (PCD) control (Fig. S3E, S3F). This shows that overexpression of E6FL and E6FLm, but not spliced E6*I isoforms, downregulate p53 protein expression in OPSCC, similar to findings in cervical cancer(*55*). Together these findings verify that the stably transfected cell lines express corresponding E6 isoforms.

To assess the functional impacts of E6FL and E6*I in OPSCC, we performed functional assays with UM-SCC-38 stably expressing E6 isoforms and the empty vector (PCD). All E6 isoforms induced proliferation in UM-SCC-38 compared to control (PCD) (Fig. S3G). Moreover, UM-SCC-38 overexpressing E6FL (expresses both E6FL and E6*I) demonstrated higher proliferation than E6FLm (non-spliceable; does not express E6*I), E6*I 3’UTR, and E6*I (Fig. S3G), suggesting that co-expression of E6FL and E6*I enhances proliferation than either isoform alone. Because overexpression of E6*I or E6*I 3’UTR yielded similar proliferation (Fig. S3G) and E6*I expression is higher than E6*I 3’UTR (Fig. S3C, S3D), E6*I was used for subsequent functional studies.

We next assessed migration, which is associated with aggressive disease and poor clinical outcomes in HNSCC(*11, 57–61*). In orthogonal wound healing and transwell migration assays, UM-SCC-38 overexpressing E6*I exhibited greater migration (Fig. 4B, S3H) than cells overexpressing PCD, E6FL or E6FLm. Thus, in this HPV(-) OPSCC cell line model, ectopic E6*I expression was sufficient to increase migration relative to vector control, whereas E6FL and E6FLm were associated with less migration than E6*I.

To assess local invasion, we used our previously established CAM model (*54*). E6*I-expressing tumors exhibited small invasive islands located at greater distances from the main tumor (Figs. 4C; S4A-S4C). These islands were smaller than those observed in the E6FL and PCD groups (Fig. S4B), and E6*I-expressing tumors generated more invasive islands than E6FLm-expressing tumors (Fig. S4C).

The CAM is primarily composed of type IV collagen and laminin and disruption of the basal laminin (de-lamination) facilitates tumor invasion (58). Therefore, we evaluated laminin staining in our CAM assay. E6FLm-expressing tumors showed the least disruption of the laminin-positive basement membrane, although this difference did not reach statistical significance (Figs. S4D, S4E). Collectively, the CAM findings indicate that E6*I is associated with a more invasive phenotype than E6FLm.

We assessed local invasion using orthotopic mouse model as we described (*62*). UM-SCC-38 stably expressing E6 variants or vector (PCD) were injected in the tongue of athymic nude mice (Fig. 4D). Tongues, harvested 21 days later, were sectioned and stained with H&E and cytokeratin (highlights epithelial cells) to identify the main tumor and invasive islands (Fig. 4D). Tumors were observed in all mice injected with UM-SCC-38-E6*I or -PCD (control) cells, and in 80% of mice injected with UM-SCC-38-FL or -FLm (Fig. S5A).

A higher proportion of mouse tongue tumors overexpressing E6*I exhibited locally invasive islands located > 400um from the main tumor than E6FLm-expressing tumors (Fig. 4E). However, the E6*I group did not differ significantly from E6FL-(express both E6FL and E6*I transcripts) or PCD-(represents HPV(-) OPSCC background) groups (Fig. 4E, 4F; S5B). Thus, our findings in mouse and CAM models support that overexpression of E6*I induces a more locally invasive phenotype compared to non-spliceable E6FLm. The mouse model suggests that E6*I did not increase invasion relative to the HPV-negative vector-control.

Since ectopic expression of E6*I induces migration and invasion compared to non-spliceable E6FLm in vitro and in vivo, we assessed if overexpression of E6 spliced variants impacts the expression and localization of epithelial-to-mesenchymal transition (EMT) markers. Immunofluorescence data in vitro (Fig. 4G; S6A) showed that ECAD, which facilitates cell-cell adhesion(*63, 64*), was mostly localized at cell junctions in UM-SCC-38-E6FL or -E6FLm; however, in -E6*I or -PCD cells, ECAD mainly localized in the perinuclear/cytoplasmic region, consistent with p-EMT (*39, 51*). Quantification of membrane:cell ECAD confirmed reduced membrane ECAD localization in UM-SCC-38-E6*I or -PCD compared to -E6FL or -E6FLm (Fig. 4H). Total ECAD and VIM intensity showed no difference between E6 variants and control (Fig. 4I, S6B). Consistent with their perinuclear/cytoplasmic ECAD localization, control- and E6*I-overexpressing cells were more isolated than FL- and FLm-overexpressing cells that were more tightly packed (Fig. 4G). Immunoblotting similarly showed no significant differences in total ECAD or VIM expression (Fig. S6C-S6E).

Taken together, these findings demonstrate that E6 variants differentially regulate proliferation, migration, invasion, and ECAD localization. Overexpression of E6*I increases migration and invasion relative to unspliced E6FLm. Overall, our findings in UM-SCC-38 expressing E6 spliced variants support the differential effects of HPV16 E6 spliced variants in migration and invasion in OPSCC.

### 5. E6*I expression promotes ECAD internalization and alters ECAD trafficking

Having established that a shift from E6FL to E6*I is associated with higher cytoplasmic and lower membrane localization of ECAD in OPSCC models, we investigated the underlying mechanism. Since redistribution of epithelial proteins via endocytosis disrupts epithelial junctions and is essential for cancer spread(*65*), we hypothesized that the expression of the E6 spliced variants differentially regulate ECAD internalization. To test this, we performed an internalization assay, as described (*66*), using an antibody targeting the extracellular domain of ECAD (HECD-1) (Fig. 5A). UM-SCC-38-E6*I had more internalized ECAD after 30 min of internalization period than -E6FL or -E6FLm; while control (PCD) cells had similar internalized ECAD (Fig. 5B, 5C). This suggests that the E6FL and E6FLm constructs reduce ECAD internalization relative to E6*I and PCD, consistent with the ECAD localization pattern observed in the stably overexpressed cells in Fig. 4G and 4H.

**Fig. 5:**
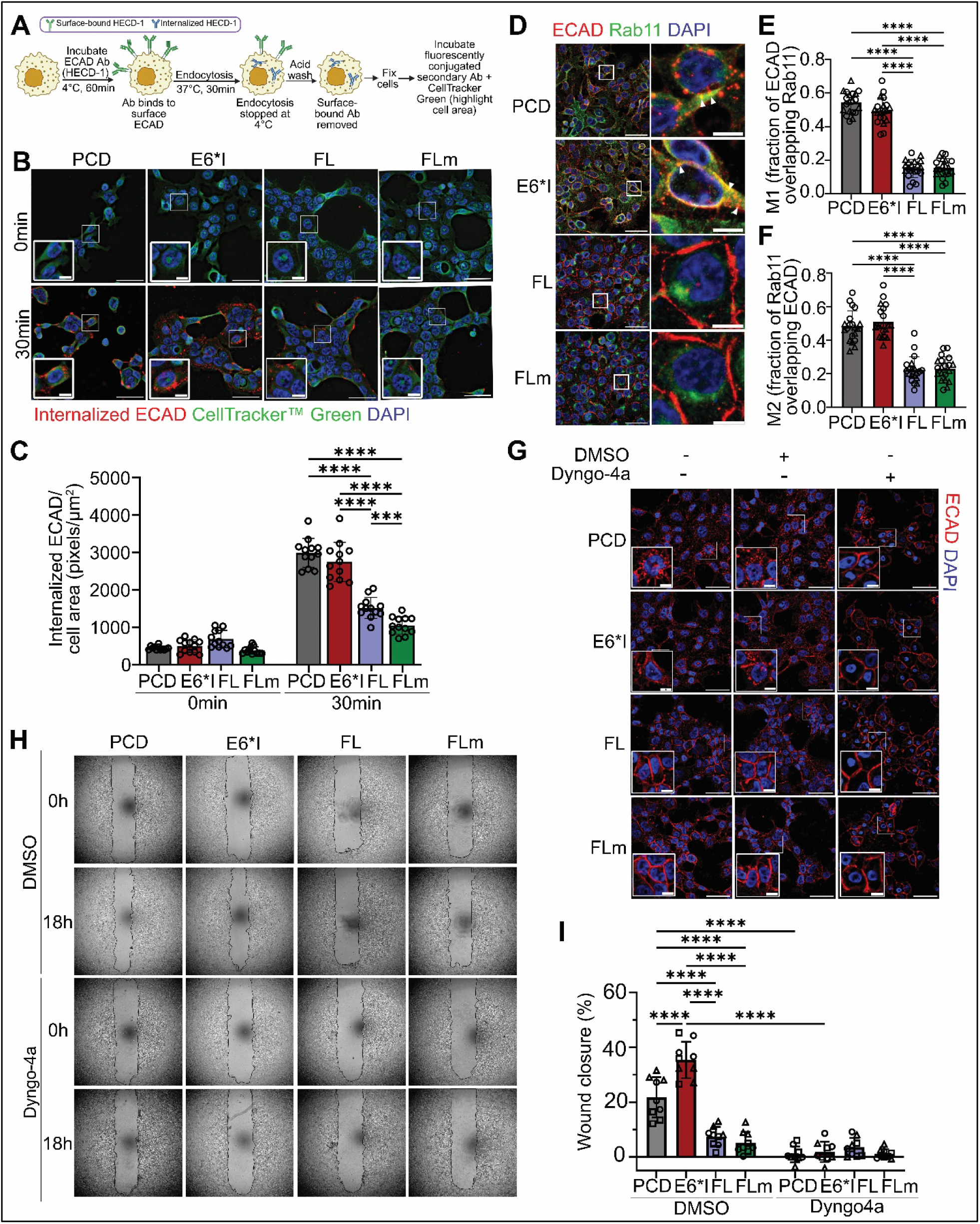
E6 splice variants differentially regulate ECAD internalization, trafficking, and OPSCC cell migration. (A) Schematic of antibody internalization assay. Created with Biorender.com (B) Representative immunofluorescence images of antibody internalization assay performed on UM-SCC-38 overexpressing PCD or E6 isoforms. Red = internalized ECAD, green = CellTracker™ Green (whole cells). Blue = DAPI (nucleus). Enlarged images of boxed areas are showed in lower left. Scale Bars: Outside=50μm, inside=10μm (C) Quantification for (B). Fluorescence intensity of internalized ECAD (red) was quantified within cell areas marked by CellTracker™ Green. *** p<0.001; **** p<0.0001 (two-way ANOVA) (D) Maximum projection of images showing co-localization of ECAD (red) and Rab11 (green) in UM-SCC-38 expressing PCD or E6 isoforms. DAPI (nucleus) is in blue. Right panel shows enlarged image of boxed area in left panel. Scale bar= 50μm (left) and 10μm (right). (E&F) Co-localization of ECAD and Rab11 was quantified from 12 z slices, 5 fields/duplicate. Manders’ coefficients, i.e., M1, fraction of ECAD overlapping Rab11 (E) and M2, fraction of Rab11 overlapping ECAD (F), were quantified using ImageJ JaCoP plugin. **** p<0.0001 (one-way ANOVA post-hoc Tukey). (G) Immunofluorescence on UM-SCC-38 with PCD or E6 isoforms treated with Dyngo-4a or DMSO vehicle. Cells were fixed 24h after treatment and stained with ECAD (red) and DAPI (blue). Enlarged images of boxed areas are showed in lower left. Scale Bars: Outside=50μm, inside=10μm. (H) Wound healing assay for UM-SCC-38 cells expressing PCD or E6 isoforms treated with either Dyngo-4a or DMSO vehicle. (I) Migration is quantified from (H) and represents percent (%) wound closure after 18h. For all graphs, data are represented as mean ± SD. *p<0.05, **p<0.01, ***p<0.001, ****p<0.0001 (One-way ANOVA post-hoc Tukey test).

Since vesicular trafficking is central to ECAD internalization and recycling (*65*), we investigated co-localization of ECAD with EEA1 or Rab11, markers for early and recycling vesicles, respectively. Co-localization of ECAD and EEA1 did not change significantly between E6 variants (Fig. S7A-S7C). In contrast, ECAD was co-localized with Rab11 more extensively in cells with overexpression of E6*I compared to FL and FLm (Fig. 5D; S7D). This co-localization of ECAD in Rab11-positive components in UM-SCC-38 overexpressing E6*I compared to E6FL or E6FLm, is consistent with previous studies implicating Rab11-mediated trafficking in ECAD redistribution in other cancer models(*39, 64, 67*).

To determine the functional relevance of E6*I-mediated ECAD internalization, we pharmacologically inhibited dynamin-mediated internalization of membrane proteins with Dyngo-4a(*68, 69*). Dyngo-4a enhanced membrane-localized ECAD in cells overexpressing E6*I and PCD, consistent with reduced dynamin-dependent ECAD internalization (Fig. 5E). In cells overexpressing FL or FLm, ECAD was mostly localized at the membrane regardless of Dyngo-4a treatment (Fig. 5E), suggesting that less internalization of ECAD occurred in these cells. Functionally, Dyngo-4a significantly reduced wound closure in cells overexpressing E6*I and PCD, but not in cells overexpressing FL and FLm (Fig. 5F, 5G), consistent with their ECAD localization patterns (Fig. 5E). Together, these findings support an association between dynamin- and Rab11-associated trafficking pathways, ECAD internalization and migration in OPSCC cells stably overexpressed with E6*I compared to E6FL or E6FLm constructs.

### 6. Lower E6FL:E6_ALL_ influence score in HPV+ OPSCC correlates with adverse clinical features and p-EMT in multi-institutional human cohorts

Having established that E6FL and E6*I transcripts have differential roles in migration and invasion in multiple preclinical models, we next investigated whether the biological influence of E6 isoform transcripts was associated with clinical outcomes in HPV+ OPSCC patients. Direct quantification of individual HPV E6 splice isoforms from retrospectively assembled RNA-sequencing datasets is technically challenging because of variable RNA quality, inconsistent transcript coverage, and differences in sequencing platforms (*70–72*). We therefore used our previously developed E6FL:E6_ALL_ influence score (also referred to as E6FL:E6_ALL_ score) as an indirect measure for E6 splicing-associated host transcriptional biology(*30*). This score is derived from 168 host genes that significantly correlated with the relative abundance of E6FL among all E6 transcripts (includes E6FL and truncated E6*) in 68 HPV+ HNSCC patients (75% OPSCC)(*30*). A lower E6FL:E6_ALL_ influence score corresponds to a host transcriptional pattern associated with lower relative contribution of E6FL and higher contribution of E6* and vice versa. Because E6*I is the predominant splice E6 transcript variant in HPV+ HNSCC (*19, 20, 45*), this pattern is biologically compatible with higher influence of E6*I transcript. However, this score does not directly quantify E6*I expression, E6FL:E6*I ratio or splicing rate.

In previous study, we correlated E6FL:E6_ALL_ influence score with overall survival in two cohorts, one from the University of Michigan (termed as UM18) and the other from The Cancer Genome Atlas (TCGA). In that previous analysis. low E6FL:E6_ALL_ influence score, reflecting higher E6*I-associated biology, was associated with overall survival(*30*), while its association with recurrence was not studied.

To establish the clinical relevance of E6 splicing specifically in OPSCC, we expanded our analysis to 230 HPV+ OPSCC patients from five non-overlapping cohorts across three institutions (Table 1; Fig. 6A). These include the TCGA and UM18 that were studied in our initial analysis, and three additional cohorts including UM67, Human Virome Consortium (HVC) cohorts and a newly introduced UM20 cohort (Table 1). Tumors that did not originate from the oropharynx were excluded. Among these cohorts, patients from TCGA, UM18, UM20 and UM67 have overall survival information (Table 1), while patients from the three UM cohorts (UM20, UM67 and UM18) had available recurrence information.

**Figure 6.**
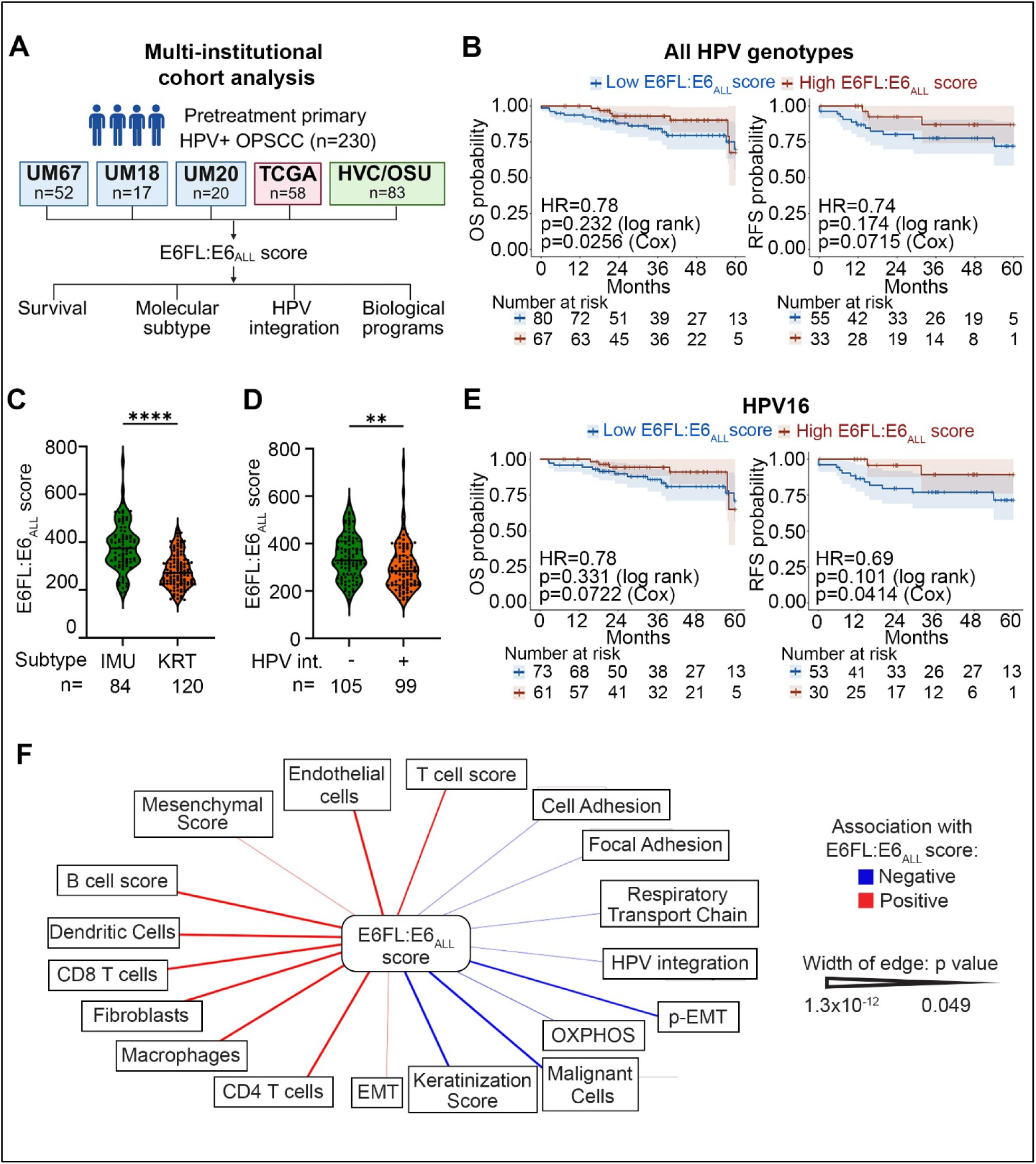
Clinical and molecular associations of the E6FL:E6_ALL_ score in patients. (A) Schematic of the multi-institutional cohort analysis. Created with Biorender.com (B) Kaplan-Meier curves showing overall survival (OS; left) and recurrence-free survival (RFS; right) among patients with tumors classified as having low or high E6FL:E6_ALL_ scores across HPV genotypes, stratified by mean. Shaded regions indicate 95% confidence intervals. Hazard ratio (HR) per 50-point increase, log-rank p and cox p (adjusted for cohort, age, overall stage) using E6FL:E6_ALL_ scores as a continuous variable are indicated. (C-D) Distribution of E6FL:E6_ALL_ scores in immune (IMU) and keratinization (KRT) molecular subtypes (C) and according to integration status (D). Each point represents one patient. Solid line represents the median. ** p<0.01; **** p<0.0001 (unpaired t test) (E) Kaplan–Meier estimates of OS (left) and RFS (right) among patients with HPV16+ tumors classified as having low or high E6FL:E6_ALL_ scores, stratified by mean. Shaded regions indicate 95% confidence intervals. Log-rank p and cox p (adjusted for cohort, age, overall stage) are indicated. (F) Network representation of associations between the E6FL:E6_ALL_ score and tumor- or microenvironment-related biological programs. Only statistically significant associations (p<0.05) are shown. Red edges indicate positive associations, and blue edges indicate negative associations. For categorical associations, including associations with T stage, HPV integration status, and subtype, logistic regression was employed with cohort as a covariate.

**Table 1:**
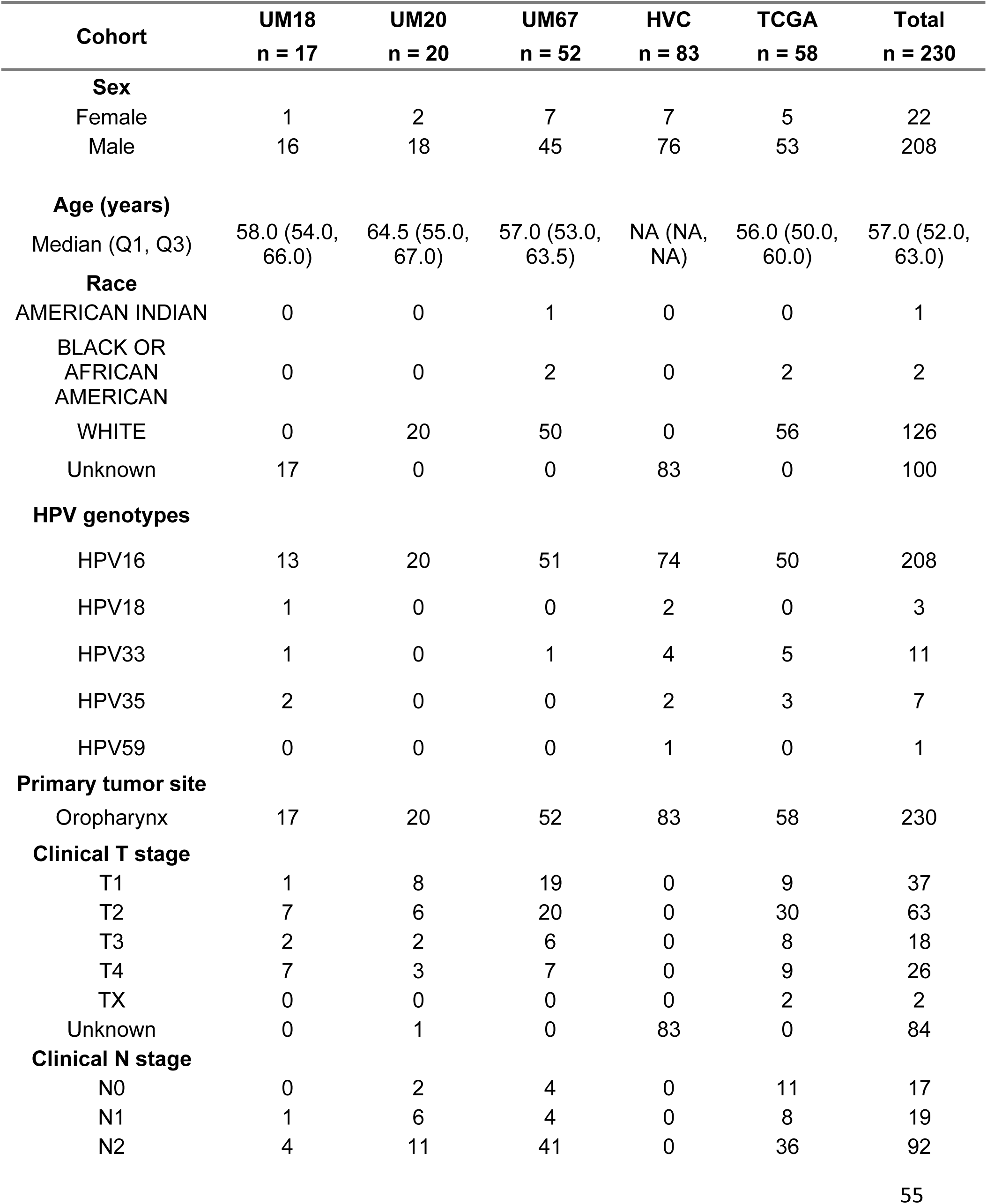
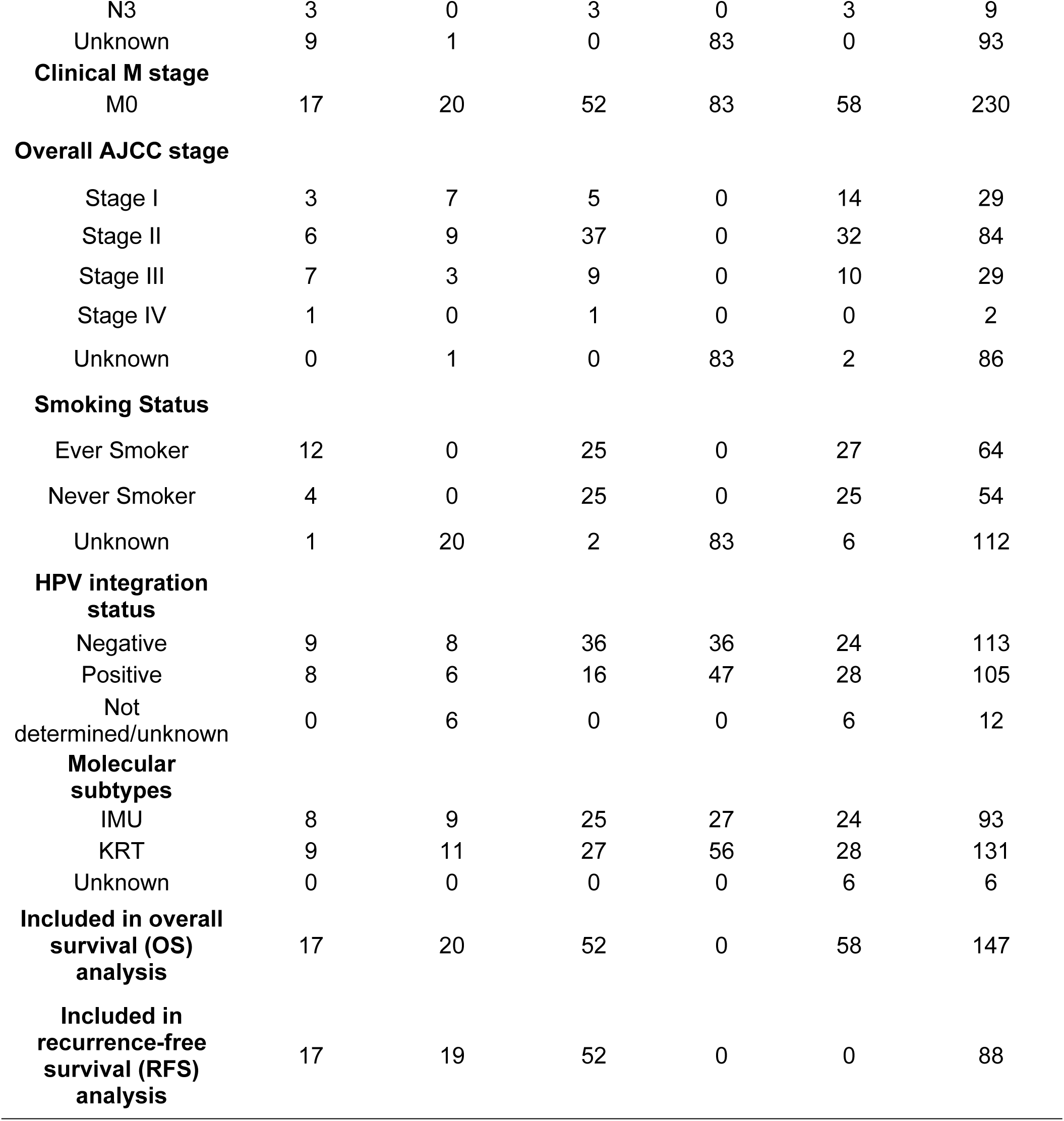
Study characteristics of RNA-sequencing patient cohorts.

| **Cohort** | **UM18**<br>**n = 17** | **UM20**<br>**n = 20** | **UM67**<br>**n = 52** | **HVC**<br>**n = 83** | **TCGA**<br>**n = 58** | **Total**<br>**n = 230** |
| --- | --- | --- | --- | --- | --- | --- |
| **Sex** |  |  |  |  |  |  |
| Female | 1 | 2 | 7 | 7 | 5 | 22 |
| Male | 16 | 18 | 45 | 76 | 53 | 208 |
| **Age (years)** |  |  |  |  |  |  |
| Median (Q1, Q3) | 58.0 (54.0, 66.0) | 64.5 (55.0, 67.0) | 57.0 (53.0, 63.5) | NA (NA, NA) | 56.0 (50.0, 60.0) | 57.0 (52.0, 63.0) |
| **Race** |  |  |  |  |  |  |
| AMERICAN INDIAN | 0 | 0 | 1 | 0 | 0 | 1 |
| BLACK OR AFRICAN AMERICAN | 0 | 0 | 2 | 0 | 2 | 2 |
| WHITE | 0 | 20 | 50 | 0 | 56 | 126 |
| Unknown | 17 | 0 | 0 | 83 | 0 | 100 |
| **HPV genotypes** |  |  |  |  |  |  |
| HPV16 | 13 | 20 | 51 | 74 | 50 | 208 |
| HPV18 | 1 | 0 | 0 | 2 | 0 | 3 |
| HPV33 | 1 | 0 | 1 | 4 | 5 | 11 |
| HPV35 | 2 | 0 | 0 | 2 | 3 | 7 |
| HPV59 | 0 | 0 | 0 | 1 | 0 | 1 |
| **Primary tumor site** |  |  |  |  |  |  |
| Oropharynx | 17 | 20 | 52 | 83 | 58 | 230 |
| **Clinical T stage** |  |  |  |  |  |  |
| T1 | 1 | 8 | 19 | 0 | 9 | 37 |
| T2 | 7 | 6 | 20 | 0 | 30 | 63 |
| T3 | 2 | 2 | 6 | 0 | 8 | 18 |
| T4 | 7 | 3 | 7 | 0 | 9 | 26 |
| TX | 0 | 0 | 0 | 0 | 2 | 2 |
| Unknown | 0 | 1 | 0 | 83 | 0 | 84 |
| **Clinical N stage** |  |  |  |  |  |  |
| N0 | 0 | 2 | 4 | 0 | 11 | 17 |
| N1 | 1 | 6 | 4 | 0 | 8 | 19 |
| N2 | 4 | 11 | 41 | 0 | 36 | 92 |
| N3 | 3 | 0 | 3 | 0 | 3 | 9 |
| Unknown | 9 | 1 | 0 | 83 | 0 | 93 |
| **Clinical M stage** |  |  |  |  |  |  |
| M0 | 17 | 20 | 52 | 83 | 58 | 230 |
| **Overall AJCC stage** |  |  |  |  |  |  |
| Stage I | 3 | 7 | 5 | 0 | 14 | 29 |
| Stage II | 6 | 9 | 37 | 0 | 32 | 84 |
| Stage III | 7 | 3 | 9 | 0 | 10 | 29 |
| Stage IV | 1 | 0 | 1 | 0 | 0 | 2 |
| Unknown | 0 | 1 | 0 | 83 | 2 | 86 |
| **Smoking Status** |  |  |  |  |  |  |
| Ever Smoker | 12 | 0 | 25 | 0 | 27 | 64 |
| Never Smoker | 4 | 0 | 25 | 0 | 25 | 54 |
| Unknown | 1 | 20 | 2 | 83 | 6 | 112 |
| **HPV integration status** |  |  |  |  |  |  |
| Negative | 9 | 8 | 36 | 36 | 24 | 113 |
| Positive | 8 | 6 | 16 | 47 | 28 | 105 |
| Not determined/unknown | 0 | 6 | 0 | 0 | 6 | 12 |
| **Molecular subtypes** |  |  |  |  |  |  |
| IMU | 8 | 9 | 25 | 27 | 24 | 93 |
| KRT | 9 | 11 | 27 | 56 | 28 | 131 |
| Unknown | 0 | 0 | 0 | 0 | 6 | 6 |
| **Included in overall survival (OS) analysis** | 17 | 20 | 52 | 0 | 58 | 147 |
| **Included in recurrence-free survival (RFS) analysis** | 17 | 19 | 52 | 0 | 0 | 88 |

We first analyzed the association of E6FL:E6_ALL_ influence score with overall survival in patients with available information. Lower E6FL:E6_ALL_ score (higher E6*I-associated biology) associated with significantly worse overall survival, after accounting for age, overall stage, and cohort (n=147; Cox p=0.0256) (Fig. 6B). Lower E6FL:E6_ALL_ score also trended toward worse recurrence-free survival (n = 88; Cox p = 0.0715) (Fig. 6B). These findings suggest that a host transcriptional state associated with higher E6*I influence relative to E6FL is associated with poorer clinical outcomes in HPV+ OPSCC.

We next evaluated whether the E6FL:E6_ALL_ influence score was associated with previously described molecular features of aggressive HPV+ OPSCC. We previously identified two molecular subtypes of HPV+ HNSCC, keratinized (KRT) and immune-rich (IMU), based on gene expression (*38, 73*). In our current analysis, KRT tumors, that are associated with worse clinical outcomes (*38, 73*), had significantly lower E6FL:E6_ALL_ influence scores (associated with higher E6*I impact) than IMU tumors (Fig. 6C). Because HPV integration positivity had been associated with aggressive disease and inferior clinical outcomes in HPV+ OPSCC(*2, 74, 75*), we assessed the relationship between E6FL:E6_ALL_ influence scores and HPV integration status, E6FL:E6_ALL_ influence scores were significantly lower in integration-positive than integration-negative tumors (Fig. 6D). Together, these findings support that lower E6FL:E6_ALL_ score is associated with molecular features linked to aggressive HPV+ OPSCC.

Because HPV genotypes may impact patient prognosis(*76, 77*), we further restricted our survival analysis to only HPV16+ OPSCC cases (Fig. 6E). In this HPV16-only subset, lower E6FL:E6_ALL_ influence score trended towards worse overall survival (Cox p=0.0722) and was significantly associated with worse recurrence-free survival (Cox p=0.0414), after accounting for age, overall stage, and cohort (Fig. 6E). These findings suggest that the relationship between lower E6FL:E6_ALL_ influence score and poor outcome is retained within HPV16+ OPSCC, particularly with respect to recurrence.

To gain insight into biological processes associated with differential E6 splicing, we correlated the E6FL:E6_ALL_ influence score with curated gene expression signatures by pooling all five cohorts. E6FL:E6_ALL_ score associated positively with cell adhesion and focal adhesion programs (Fig. 6F), suggesting that tumors with greater E6FL influence retain stronger adhesion-associated transcriptional features. Unexpectedly, E6FL:E6_ALL_ score also correlated positively with complete EMT (Fig.6F), which is conventionally associated with increased invasion and poorer prognosis(*78, 79*). This apparent paradox prompted us to examine whether E6 splicing preferentially associates with intermediate EMT states rather than complete EMT, given emerging evidence that intermediate EMT states rather than full EMT drive invasion in HNSCC(*80, 81*). Using an established p-EMT signature(*80*), we observed significant inverse correlation between E6FL:E6_ALL_ score and p-EMT (Fig. 6F), indicating higher p-EMT activity in tumors with greater E6* influence.

Previous studies associated reduced oxidative stress and heightened immune response with improved prognosis in OPSCC (*45, 82–85*). Our analysis demonstrated that E6FL:E6_ALL_ influence score is directly correlated with immune infiltration and negatively with oxidative phosphorylation and respiratory electron transport chain (Fig. 6F). Collectively, these findings suggest that greater E6*I influence is associated with a p-EMT transcriptional state and other tumor-associated metabolic and immune features in HPV+ OPSCC. These patient-level findings align with our preclinical data and support a model in which splicing shift toward E6*I promotes epithelial plasticity and aggressive tumor behavior.

### 7. Reduced membrane ECAD localization is associated with recurrences and lower E6FL:E6_ALL_ influence scores in HPV+ OPSCC

Our preclinical OPSCC models suggested that a shift from E6FL towards E6*I promotes p-EMT-associated redistribution of ECAD from membrane to cytoplasm relative to E6FL without markedly reducing total ECAD protein expression. We therefore investigated whether ECAD membrane-to-cytoplasmic localization was associated with E6 splicing and clinical outcomes in patients with HPV+ OPSCC.

We evaluated ECAD localization in two independent, non-overlapping cohorts of 75 FFPE primary HPV+ OPSCC, comprising discovery (Cohort 1; n=47) and validation (Cohort 2; n=28) cohorts (Fig. 7A). The discovery cohort consists of 8 patients from UM18, and 39 patients from UM67, while the validation cohort consists of 3 patients from UM18 (n=3), UM67 (n=5) and UM20 (n=20) (Fig 7A; Table S2). Tissue specimens with insufficient material were excluded. All patients in Cohort 1 have both overall survival and recurrence information. All patients in Cohort 2 have overall survival information, and all but one patient have recurrence information (Table S3).

**Fig. 7:**
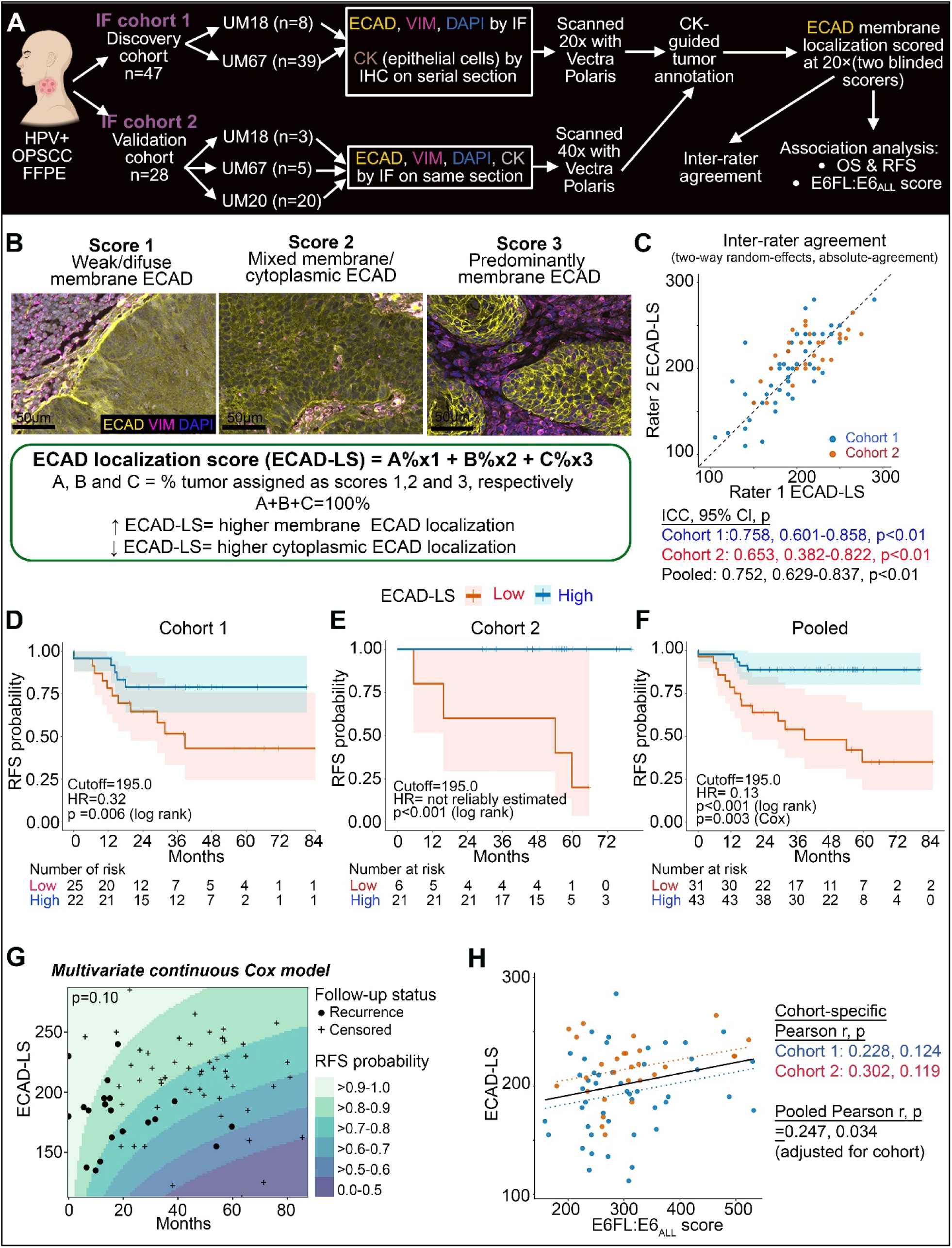
Reduced membrane E-cadherin localization associated with recurrences in HPV+ OPSCC. (A) Study workflow created with Biorender.com. Formalin-fixed, paraffin-embedded (FFPE) HPV+ OPSCC specimens were divided into discovery (cohort 1) and validation (cohort 2) cohorts. ECAD membrane localization was independently evaluated by two blinded raters (B) Representative images of tumors with weak/diffuse (Score 1), mixed membrane/cytoplasmic (Score 2), or predominantly membrane ECAD localization (Score 3). ECAD is in yellow, VIM in magenta and DAPI in blue. The E-cadherin localization score (ECAD-LS) is a weighted composite of the percentage of tumor assigned to each pattern; higher scores indicate greater membrane localization. Scale bars, 50 μm. (C) Inter-rater agreement for ECAD-LS using a two-way random-effects, absolute-agreement intraclass correlation coefficient (ICC). (D–F) Recurrence-free survival according to low versus high ECAD-LS, dichotomized at the median from Cohort 1 (195.0) as the locked threshold. Shaded areas indicate confidence intervals, tick marks indicate censored observations, and numbers at risk are shown below each plot. (G) Multivariable continuous Cox model relating ECAD-LS to RFS over time. Background colors represent model-estimated RFS-probability intervals; filled circles indicate observed recurrences and plus signs indicate censored observations (H) Correlation between ECAD-LS and the E6FL:E6_ALL_ score. Points represent individual tumors from cohort 1 (blue) and cohort 2 (orange); dotted lines show cohort-specific linear fits and the solid black line shows the pooled fit. For cohort-specific correlation, pearson correlation was analyzed. For pooled cohort, a partial pearson adjusting for cohort was used.

Multiplex immunofluorescence was performed in Cohorts 1 and 2 separately using ECAD, VIM, and DAPI (Fig. 7A; S8A). Cytokeratin staining, performed either by immunohistochemistry on a serial section (Cohort 1) or as part of the multiplex panel (Cohort 2), was used to guide annotation of tumor epithelial regions (Fig. 7A; Fig. S8A). VIM helped delineate the cytoplasm and distinguish membrane versus non-membrane ECAD signal, and DAPI identified the nuclei (Fig. S8A). This approach enabled ECAD localization to be evaluated specifically within viable tumor regions while excluding stromal, inflammatory. Epithelial cells in non-neoplastic superficial epithelium l were excluded on histologic review.

ECAD localization varies substantially among tumors (Fig 7B). Some HPV+ OPSCC showed strong, crisp, continuous membranous ECAD staining, whereas others showed weak membrane ECAD with diffuse cytoplasmic staining. To quantify this pattern, viable tumor cells were assigned one of these three localization categories: Score 1, weak or diffuse membrane ECAD; Score 2, mixed discontinuous membrane and cytoplasmic ECAD; and Score 3, predominantly strong, crisp and continuous membrane ECAD (Fig. 7B). An ECAD localization score (ECAD-LS) was then calculated as:

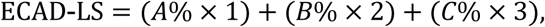

where A, B, and C represented percentages of viable tumor cells assigned scores 1, 2, and 3, respectively, and added up to 100%. Thus, ECAD-LS ranges from 100 to 300, with higher values indicating greater membrane localization. Percentages of viable tumor cells were visually estimated. A higher ECAD-LS reflects greater membrane localization, whereas a lower ECAD-LS reflects reduced membrane and increased cytoplasmic ECAD localization.

Two investigators blinded to clinical and molecular data independently scored each tumor; one scorer is a board-certified pathologist. All tumor regions were assessed. Necrotic regions, stroma, immune cells, non-neoplastic epithelium, tissue folds, and regions with inadequate tissue or staining quality were excluded. Inter-rater agreement was between moderate and good, with intraclass correlation coefficients of 0.758 in Cohort 1, 0.653 in Cohort 2, and 0.752 across the pooled cohort (all p<0.01; Fig. 7C). The reproducibility of the ECAD-LS across two independent scorers, including a board-certified pathologist, supports the feasibility of this scoring approach. The average ECAD-LS from both scorers was therefore used for subsequent analyses.

We next examined the association between ECAD localization and recurrence-free survival (RFS). In Cohort 1, patients were dichotomized using the cohort median ECAD-LS of 195.0 (Fig. S8B; Fig. 7D); patients with low ECAD-LS had significantly shorter RFS than those with high ECAD-LS (HR=0.32 for high versus low ECAD-LS; log-rank p=0.006; Fig. 7D). Next, we tested this association in Cohort 2 as an independent and non-overlapping cohort. Because one patient in Cohort 2 lacked recurrence information, this patient was excluded from the RFS analysis, leaving 27 patients for analysis. (Table S3). To test the association of ECAD-LS with RFS without deriving a new cutoff, we used median ECAD-LS derived from cohort 1 (195.0) and applied it as a locked cutoff to Cohort 2 (Fig. 7E). Because all events occurred in a single group, the hazard ratio was not estimable (Fig. 7E; Table S3). Nevertheless, the difference in RFS was significant by log-rank analysis (p<0.001) (Fig. 7E). These findings support the direction of the discovery-cohort result but should be interpreted cautiously given the limited number of events.

As a sensitivity analysis, we also dichotomized Cohort 2 using its own median ECAD-LS of 218.75 (Fig. S8B; S8C, left). This analysis again (Fig. S8C, left) showed shorter RFS among patients with low ECAD-LS, although the difference did not reach statistical significance (log-rank p=0.055). The hazard ratio remained non-estimable because no recurrences occurred in the high-ECAD-LS group (Fig. S8C; Table S3). When the two cohorts were pooled and dichotomized using the pooled-cohort median (205.0; Fig. S8B), low ECAD-LS was significantly associated with shorter RFS by log-rank analysis (p<0.01; Fig. S8C, right). This association remained significant in a Cox proportional hazards model adjusted for age, AJCC stage, and cohort (p=0.036). A pooled analysis using the locked Cohort 1 cutoff produced a similar association (Fig.7F).

Because ECAD-LS is intrinsically continuous, we additionally evaluated it without dichotomization. In a multivariable Cox model of the pooled cohort adjusted for age, AJCC stage, and cohort, lower continuous ECAD-LS showed a trend toward a greater risk of recurrence, but the association was not statistically significant (p=0.10; Fig. 7G). Thus, the RFS analyses consistently showed the same direction of effect, although the strength of the association depended on the analytical approach and was limited by the small number of recurrence events, particularly in Cohort 2.

We also evaluated the association between ECAD localization and overall survival (OS). Low ECAD-LS was associated with shorter OS in unadjusted log-rank analyses using either the locked Cohort 1 cutoff (Fig. S8D, left) or the pooled-cohort median (Fig. S8D, right). In Cox models adjusted for age, AJCC stage, and cohort, continuous ECAD-LS was also not significantly associated with OS in the multivariable Cox model (p=0.65; Fig. S8E). The clinical association of ECAD localization was therefore more evident for recurrence than for overall survival. These findings support ECAD localization as a candidate prognostic biomarker for pretreatment and primary HPV+ OPSCC.

Since our preclinical studies showed that splicing of E6FL into E6*I promotes the redistribution of ECAD from membrane to cytoplasm, without altering ECAD protein expression, we tested whether ECAD localization was associated with E6FL:E6_ALL_ score. In the pooled cohort, ECAD-LS was positively correlated with the E6FL:E6_ALL_ score, even after adjusting for cohort differences (Partial pearson r=0.247, p=0.034; Fig. 7H). The individual cohorts showed associations in the same direction, although neither reached statistical significance independently. Thus, tumors with lower E6FL:E6_ALL_ scores, reflecting a relatively greater contribution from E6* isoforms, tended to have less membrane (more cytoplasmic ECAD), which is consistent with our preclinical findings.

Finally, we investigated whether the observed associations reflected ECAD localization rather than overall ECAD or VIM expression. Total ECAD and VIM intensities were quantified within annotated tumor regions using Akoya inForm image analysis software. This digital measurement provided an orthogonal assessment of total protein expression specifically in the tumor region and was distinct from the pathology-based ECAD-LS, which was designed to capture subcellular localization. Stromal cells were excluded from the quantification. Total ECAD protein intensity was not associated with the E6FL:E6_ALL_ score in Cohort 1 but was inversely associated with the score in Cohort 2 (Fig. S9A). Total VIM protein expression was not significantly associated with the E6FL:E6_ALL_ score in either cohort (Fig. S9B). Moreover, neither total ECAD nor total VIM expression was significantly associated with RFS or OS in either cohort (Fig. S9C, S9D). Therefore, ECAD localization showed a more consistent relationship with recurrence than did total ECAD or VIM expression.

Together, these findings extend our mechanistic observations to HPV+ OPSCC patient specimens. Reduced membrane ECAD localization is associated with lower E6FL:E6_ALL_ scores and shorter recurrence-free survival, whereas total ECAD and VIM abundance do not consistently associate with clinical outcome. These results support a model (Fig. 8) in which altered E6 splicing promotes redistribution of ECAD away from the membrane, consistent with p-EMT, rather than uniformly reducing ECAD expression, and potentially contributes to enhanced invasion and recurrence in HPV+ OPSCC.

**Figure 8.**
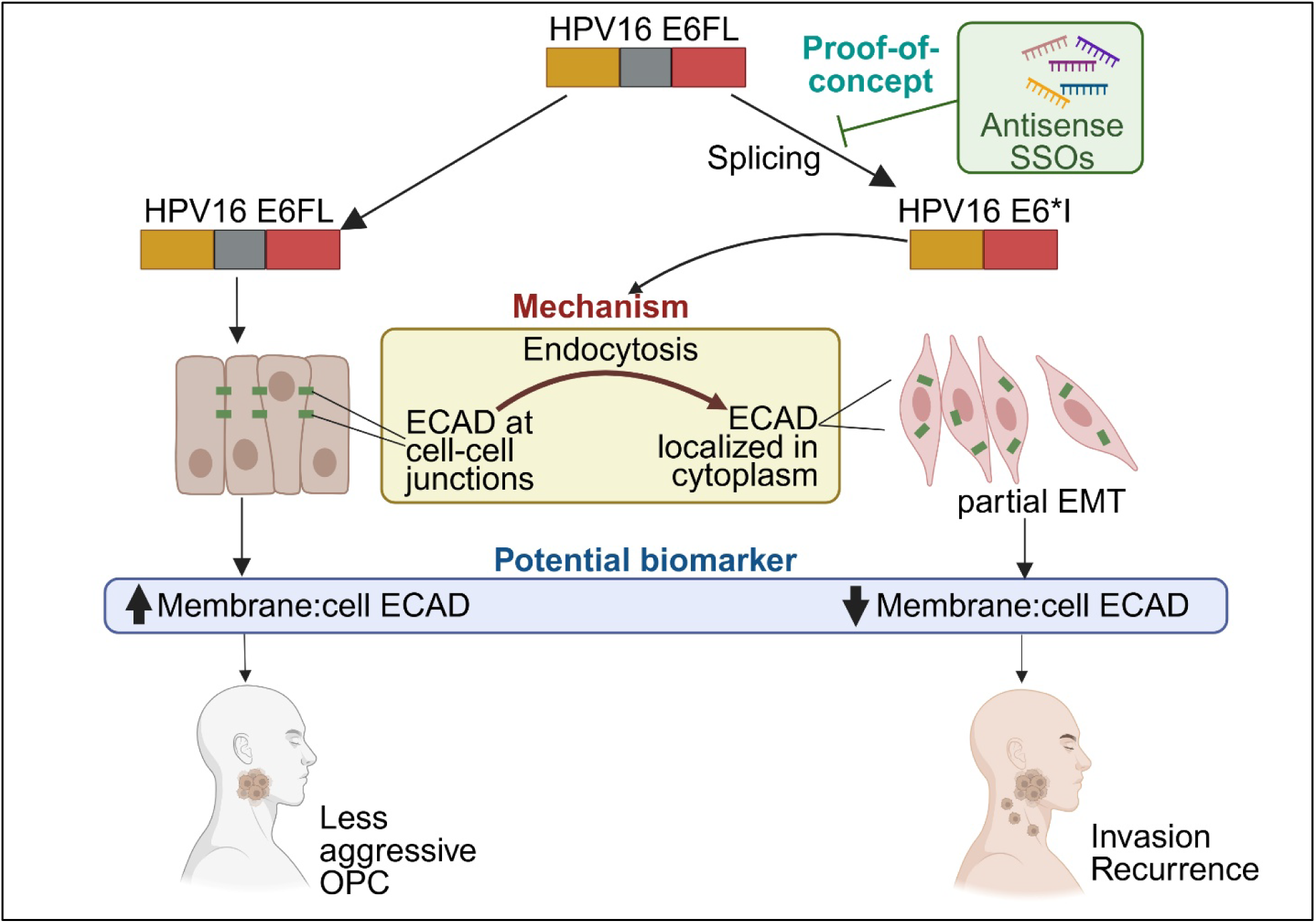
Schematic diagram of study

## Discussion

Approximately one-quarter of HPV+ OPSCC recurs or persists after treatment but the underlying mechanisms are unclear. When HPV+ OPSCC recurs, salvage therapy offers limited benefit, resulting in two-year overall survival of only approximately 50%(*9*). A few biomarkers have been proposed but either failed to reliably identify this subset of patients with aggressive cancer at diagnosis or require further external validation(*2, 4*). By integrating complementary preclinical models with multi-institutional patient cohorts, our study suggests HPV16 E6 RNA splicing as an underappreciated determinant associated with epithelial plasticity and invasion. Our findings support a model in which a splicing shift toward E6*I is associated with redistribution of E-cadherin (ECAD) away from cell-cell junctions, thereby facilitating p-EMT and tumor spread. Pathology-based assessment of ECAD localization may provide a clinically accessible pretreatment marker of this biology and recurrence risk in patients with primary HPV+ OPSCC.

Although E6 and E7 are well-established drivers of HPV+ cancer, the role of viral RNA splicing in tumor progression has received substantially less attention. Studies investigating E6FL and E6*I in OPSCC are limited by conflicting results(*30, 45, 82, 86*), while others use cervical cancer or non-oropharyngeal models that do not fully reflect the unique biology of OPSCC (*18, 24–28*). Using orthogonal OPSCC-specific models, we demonstrated that invasive phenotypes are influenced by the relative balance between E6FL and E6*I transcripts. *In vivo*, tumors with a relative shift toward E6*I exhibited locally invasive tumor islands located farther away from the main tumor than tumors with reduced E6 splicing; this corresponds to an aggressive clinical presentation of HPV+ OPSCC, where invasive tumor cells are found a distance away from the periphery of the main tumor (*31, 34, 87*).

Our findings suggest that E6 splice composition may have distinct effects on proliferation and local invasion. In the overexpression model, splice-competent E6FL, which produced both E6FL and E6*I transcripts, enhanced proliferation more strongly than either E6*I or splice-deficient E6FLm alone. This finding suggests that co-expression of E6FL and E6*I may support proliferation. By contrast, among single-cell clones with comparable total E6 transcript abundance, lower E6FL:E6*I transcript ratios, indicating a relative shift toward E6*I, were associated with increased proliferation. In SSO-treated HPV+ OPSCC cells, SSOs that increased the E6FL:E6*I transcript ratio reduced or trended toward reduced proliferation, whereas SSOs that favored E6*I did not significantly alter proliferation relative to the scrambled control. These findings indicate that the relationship between E6 splice composition and proliferation may depend on experimental context or other features of the endogenous viral and cellular background.

Across complementary models (overexpression, SSO, single-cell clone), a relative shift toward E6*I was consistently associated with migration and local invasion. In the endogenous clones and SSO models, these phenotypic differences occurred without substantial changes in total E6 transcript abundance. This interpretation is consistent with our previously published finding that total E6 and E7 expression did not differ significantly between the prognostically distinct immune-rich (IMU) and keratinized (KRT) subtypes, whereas proportion of E6FL among all E6 transcripts differed (*45*). Together, these observations suggest that relative transcript composition may capture information not reflected by total E6 transcript expression.

The observation that E6*I promoted invasion without reducing p53 also suggests that this phenotype may not require canonical E6FL-mediated p53 degradation This is consistent with previously published reports(*18, 82*),. Nevertheless, p53 protein abundance alone does not establish the functional transcriptional activity of stabilized p53. Future studies will be needed to determine how HPV16 E6 splice composition affects p53 activity and whether this contributes to invasive phenotype in OPSCC.

Although some studies report that E6*I promotes E7 translation(*47, 48*), others did not find a consistent correlation(*49, 50*). Thus, comparable total E6 transcript levels do not necessarily predict increased E7 protein as E6FL:E6*I transcript ratios decrease. In the SSO model, total E6 transcript abundance was comparable across conditions, whereas SSO-1 and SSO-2 increased the endogenous E6FL:E6*I transcript ratio and increased E6FL protein abundance. However, SSO-mediated modulation of E6 splicing did not significantly alter E7 protein expression. Similarly, E7 protein and its downstream markers did not differ substantially between the low- and high-ratio single-cell clones. These findings argue against altered E7 activity as the principal explanation for the observed phenotypes, although they do not exclude more subtle effects (such as post-transcriptional or translational) of E6 splicing on E7 regulation.

Interestingly, E6*I-overexpression and HPV-negative vector-control conditions shared selected features, including reduced junctional ECAD and increased ECAD internalization relative to E6FL- and E6FLm-expression conditions. This observation raises the possibility that a splice shift toward E6*I promotes an epithelial-plasticity state resembling selected features of HPV-negative HNSCC, which generally has poorer clinical outcomes than HPV+ OPSCC(*88*). However, these comparisons were based on a single HPV-negative cell line. Additional studies are therefore needed to determine whether this similarity extends beyond ECAD localization and trafficking. Furthermore, although an E6*I protein exists(*18, 89–91*), endogenous E6*I protein could not be reliably detected by immunoblotting. Therefore, the present study does not establish whether the observed effects are mediated directly by E6*I protein. Rather, the preclinical models show that alterations in the E6FL:E6*I transcript ratio are associated with proliferation, migration, local invasion, and ECAD localization.

Altered ECAD trafficking provides a potential and novel mechanism connecting E6 splicing to invasion. When ECAD protein is localized at adherens junctions, it maintains adhesion to neighboring cells; consequently, its redistribution away from the membrane weakens cell-cell adhesion. This phenotype occurs without significantly altering total protein expression, supporting a p-EMT state rather than a conventional complete EMT phenotype(*40*). Such an intermediate state may enable tumor cells to acquire motility while retaining epithelial characteristics required for collective invasion and survival.

The co-localization of ECAD and Rab11 in cells with higher E6*I transcript relative to E6FL thus suggests the contribution of endosomal trafficking, consistent with previous studies(*63*). Inhibition of dynamin-dependent endocytosis also increased membrane ECAD and reduced migration. However, static colocalization cannot distinguish ECAD recycling, retention, or subsequent trafficking, and dynamin inhibition affects multiple membrane proteins and cellular processes. Future studies using live-cell trafficking, genetic perturbation of specific endocytic regulators, and ECAD-directed rescue experiments will be needed to determine whether a splice shift toward E6*I alters trafficking machinery directly or acts through other host signaling pathways.

Since direct quantification of HPV splice variants, including E6FL and E6*I transcripts, from RNA-sequencing datasets can be confounded by technical challenges including non-linear sensitivity to RNA quality, fragment read lengths and platform variability(*70–72*), we used our previously developed E6FL:E6_ALL_ influence score. This host gene expression-based score is associated with the relative abundance of E6FL among E6 transcripts that leverage high-quality data from TCGA and UM18 cohorts(*30*). It is not a direct measure of E6FL:E6*I transcript ratio, E6*I protein expression, or splicing rate. Rather, the score summarizes a host transcriptional program associated with relative E6FL versus E6*I transcript abundance in previously defined UM18 and TCGA cohorts and can be applied across heterogeneous transcriptomic datasets. Because UM18 and TCGA contributed to the original development of the score in our previous study(*30*), overall survival analyses that include these cohorts should not be interpreted as fully independent validation. However, recurrence-free survival was not evaluated in the original study(*30*), and recurrence outcomes were not used to derive, select, or weight the genes included in the influence score. The recurrence analysis reported here therefore represents a new evaluation of the previously defined score in our present study.

In our expanded meta-analysis in multi-institutional cohorts, lower E6FL:E6_ALL_ influence scores (associated with higher E6*I impact), were associated with poorer overall survival and trended toward worse recurrence-free survival, and vice versa. When the analysis was restricted to HPV16+ OPSCC, a lower score was significantly associated with worse recurrence-free survival and showed a trend toward worse overall survival. Lower scores (associated with higher E6*I impact) were also observed in keratinized and HPV integration-positive tumors, both of which have been associated with aggressive disease (*2, 38, 73–75*). Collectively, these findings support the clinical relevance of E6 splice-associated biology. Future studies using targeted isoform-specific RNA in situ hybridization, single-cell analysis, or spatial profiling will be important for validating the influence score and the role of E6 splice balance in tumor progression.

A major translational implication of our study is that the subcellular localization of ECAD may provide more clinically relevant information than total ECAD expression in HPV+ OPSCC. Loss of total ECAD is a well-established feature of complete EMT and has been associated with invasion and poor prognosis across many cancers(*92*). However, tumor cells undergoing p-EMT can retain ECAD while repositioning it away from cell junctions, thereby weakening cell-cell adhesion and acquiring invasive properties without losing epithelial identity(*39, 40*). In this setting, measurements of total ECAD protein may not distinguish junctional ECAD from non-junctional or cytoplasmic ECAD. Consistent with this concept, total ECAD and VIM expression were not consistently associated with clinical outcomes or E6FL:E6_ALL_ score, in contrast to reduced membrane ECAD. Therefore, localization of ECAD may capture a functionally important state of epithelial plasticity that is not adequately represented by conventional measurements of EMT-marker abundance.

Previous studies have investigated the prognostic relevance of total membrane ECAD in HNSCC(*93–98*). However, these studies focused on tumors of the oral cavity, larynx and hypopharynx rather than OPSCC and the HPV status of the tumors was not clarified(*93–98*). Our study focuses on HPV+ OPSCC and links ECAD membrane-to-cytoplasm redistribution to HPV E6*I-associated invasion and recurrence. Therefore, ECAD localization may serve as an easily deployable readout of epithelial plasticity associated with HPV E6 splicing. To quantify ECAD localization, we developed tissue-based ECAD-LS that measures the extent of tumor cell involvement with varying ECAD localization patterns. Because ECAD-LS relies on interpretation of ECAD localization in routinely collected pathology specimens, it could potentially be incorporated into established diagnostic workflows without requiring sequencing infrastructure or specialized bioinformatics. If adapted and validated using conventional ECAD immunohistochemistry, ECAD-LS may provide an affordable and scalable approach to recurrence-risk assessment, particularly in regions where molecular profiling is unavailable, costly or technically challenging (*99, 100*). Pathologist-guided scoring may also be feasible in settings without automated image-analysis platforms.

In this study, ECAD-LS should be considered as a candidate prognostic marker rather than a clinically validated test. Although the locked discovery-cohort-cutoff analysis in the second non-overlapping cohort supported the results in the discovery cohort, there were only four recurrences in the Cohort 2, limiting precise estimation of the association. Dichotomization of the Cohort 2 by its own median showed a trend toward worse RFS among patients with low ECAD-LS, whereas continuous ECAD-LS was not significantly associated with recurrence-free survival after multivariable adjustment in the pooled cohort. Clinical translation will therefore require larger, geographically diverse cohorts, standardized staining and scoring protocols, assessment of inter-laboratory reproducibility, and determination of whether ECAD-LS adds prognostic value beyond stage, smoking history, treatment, HPV and p16 status, and established histopathologic features. Direct comparison between multiplex immunofluorescence and conventional immunohistochemistry will be particularly important before implementation in settings with fewer resources.

Our SSO studies provide proof of concept that endogenous E6 splicing can be experimentally modulated to alter invasive phenotypes in preclinical models. However, shifting the splice balance toward E6FL could enhance canonical E6FL functions, including p53 degradation; the net effect on tumor growth, treatment response and survival is unclear. Therapeutic development will require isoform-specific pharmacodynamic studies, off-target assessment, delivery optimization and evaluation of tumor control. Our findings therefore provide a rationale for investigating E6 splicing as a potential therapeutic target but do not yet establish that minimizing E6 splicing will provide clinical benefit.

In conclusion, our comprehensive investigation identifies HPV16 E6 splice composition as a determinant associated with epithelial plasticity and aggressive behavior in HPV+ OPSCC. SSOs that shifted E6 spliced transcript composition away from E6*I reduced migration and invasion in preclinical OPSCC models, providing proof of concept for further therapeutic investigation. ECAD-LS is a candidate prognostic marker with strong potential to be scalable and clinically actionable due to its accessibility even in resource-limited settings. However, further external validation and adaption to conventional pathology workflow is needed. Together, our findings provide the foundation for future mechanistic and translational studies of E6 splicing, biomarker validation, and targeted therapy development in HPV+ OPSCC.

## Materials and Methods

### Study design

This study was designed to investigate the oncogenic impact of HPV E6FL and its spliced isoform E6*I in HPV+ OPSCC. We hypothesized that splicing of HPV E6FL oncogene into its shorter E6*I isoform promotes aggressive behavior in OPSCC.

E6FL and E6*I isoform expression was manipulated using three complementary approaches: stable isoform overexpression, single-cell clone isolation with endogenous variations of E6FL and E6*I, and SSOs to shift endogenous E6 splicing. RT-PCR, RT-qPCR were performed to confirm expression of E6 isoforms. Immunoblot revealed effects on the protein expression of HPV E6 and E7, as well as their downstream targets p53, Rb and p16, and on ECAD and VIM for EMT. Oncogenic phenotypes were determined by proliferation, wound healing, and transwell assays. In all wound healing assays, mitomycin C was added to minimize contribution of proliferation. Immunofluorescence revealed cellular localization of ECAD and VIM for EMT.

For rigor and reproducibility, in vitro studies (except immunofluorescence) included at least three independent experiments using orthogonal approaches in multiple cell lines. The number of experiments and replicates are stated in each figure. Immunofluorescence was performed once or twice; for each experiment, with capture of at least 4-6 images from each duplicate, for a total of 8-12 images per condition in each experiment. Representative images are shown in figures. Both CAM and mouse experiments were performed to study invasion *in vivo*. For CAM assays, each group had 3-6 eggs. Sample size for CAM was selected by prior experience; no power analysis was performed. Eggs were randomized into groups before grafting cancer cells. For mouse experiments, 50 mice were randomized into 4 groups, with 10-14 mice per group based on power analysis. Investigators (YXL, ML) who assessed the results were not blinded to the intervention. Data collection was stopped at experiment endpoint as stated in Materials and Methods.

Transcriptomic and clinical data from 230 patients with HPV+ OPSCC were obtained from five cohorts, as described below. Formalin-fixed, paraffin-embedded (FFPE) tumor specimens from 75 patients were used for multiplex immunofluorescence and ECAD localization analyses.

### Cell culture

HOK16B (RRID: CVCL_B405) was obtained from Dr. No-Hee Park and cultured as described (*101, 102*). UPCI:SCC090 (RRID: CVCL_1899) and UPCI:SCC154 (RRID: CVCL_2230) were purchased from American Type Culture Collection (Manassas, VA, USA; ATCC CRL-3239 and CRL-3241 respectively) and cultured according to ATCC instructions. UM-SCC-38 (RRID: CVCL_7749), UM-SCC-47 (RRID: CVCL_7759) and UM-SCC-104 (RRID: CVCL_7712) were obtained from Dr. Thomas Carey and cultured as described (*103*). All cell lines were genotyped at the University of Michigan Sequencing Core or GeneCopeia (Rockville, MD, USA). The genotypes for UM-SCC cell lines were validated against published sequences (*104, 105*). The genotypes for ATCC cell lines were validated against provided sequences.

### Patient cohorts

Five patient cohorts were used in this study, including (1) our previously well-characterized University of Michigan cohort of 17 HPV+ HNSCC (referred to as UM18; GEO #GSE74956) (*45*), (2) 58 HPV+ HNSCC from The Cancer Genome Atlas (TCGA) (*13*), (3) 83 HPV+ OPSCC from HPV Virome Consortium (*21, 106*) (referred to as HVC; European Genome-phenome Archive EGAD00001004366), (4) 52 HPV+ OPSCC from the UM67 cohort (EGA Dataset from EGAS50000000893) (*73*), and a newly introduced cohort of 20 HPV+ OPSCC from the University of Michigan (referred to as UM20). We are in the process submitting the information from UM20 dataset under the same study as UM67 (EGAS50000000893). The clinical and demographic information of each cohort are summarized in Table 1.

For the UM20 cohort, tumor blocks were collected from patients diagnosed with previously untreated primary OPSCC between 2015 and 2018. This cohort was from the Michigan Medicine tumor biorepository collected in collaboration with the UM Rogel Comprehensive Cancer Center. Recurrence and overall-survival outcomes were abstracted using the same protocol as for UM67 (*73*). For both the UM67 and UM20 cohorts, written informed consents were obtained, and the study was approved by the UM Institutional Review Board. Epidemiologic, demographic and follow-up information for survival were collected.

### Plasmid construction and SSOs

Antisense SSOs used in this study were designed by Integrated DNA Technologies (Coralville, IA, USA) and synthesized as 2’-O-methoxyethyl modified oligonucleotides with phosphorothioate (PS) backbone modification by Genscript (Piscataway NJ, USA). Sequences of SSOs are listed in Table S1.

E6 plasmids were designed and constructed by the University of Michigan Vector Core (RRID:SCR_026696). Primers that were used to amplify DNA for subcloning are listed in Table S4. Briefly, polymerase chain reaction (PCR) was performed on pLentiLox HPV16 E6-E7 IRES-PURO to amplify a fragment encoding E6FL using E6-BamHI-F as the forward primer and E6FL-EcoRI-R as the reverse primer. To amplify E6*I, PCR was performed on the same plasmid using E6-BamHI-F as the forward primer and E6*I-EcoRI-R as the reverse primer. PCR fragments were subcloned into the BamHI/EcoRI sites of pcDNA3 vector to generate pcDNA3-E6FL and pcDNA3-E6*I.

To generate pcDNA3-E6FLm, PCR was performed on p1321-HPV16 E6/E7 (Addgene #8641) using E6FLm-BamHI-F and E6-splice-mutant-mega-R as the forward and reverse primers, respectively to generate a megaprimer (*107*). This PCR product was gel purified and used as a forward primer for megaprimer mutagenesis (*107*), where E6FLm-EcoRI-R primer was used as the reverse primer. The resulting PCR product containing the splice mutation that changed Val42 to Leucine was cloned into BamHI/EcoRI digested pcDNA3. A silent mutation to Arg40 codon was also observed in the final pcDNA3-E6FLm construct. To generate pcDNA3-E6*I 3’UTR, PCR was performed on p1321-HPV16 E6/E7 using E6*I-3’UTR-BsmGI-F and E6*I-3’UTR-EcoRI-R as forward and reverse primers, respectively. Amplified product was cloned into the BsmGI/EcoRI site of pcDNA3-E6*I to generate pcDNA3-E6*I 3’UTR.

### Cell transfection

For UM-SCC-38, empty vector (pcDNA3), E6*I, E6*I 3’UTR, E6FL and E6FL mutant plasmids were transfected using Amaxa Cell Line Nucleofector kit V (Lonza, Basel, Switzerland, # VCA-1003) with the U20 program according to the manufacturer’s instructions.Stably transfected cells were generated in the presence of Geneticin (G418; Gibco). UM-SCC-38-pcDNA3, UM-SCC-38-E6*I, UM-SCC-38-E6*I 3’UTR were maintained in 250 µg/ml G418, while UM-SCC-38-E6FL and UM-SCC-38-E6FLm were maintained in 150 µg/ml G418. G418 was removed from culture media for all experiments.

For SSO transfection, cells at 40% confluence were transfected with SSOs using Lipofectamine RNAimax (Invitrogen, Waltham, MA, USA, # 13778075). 24 h after transfection, cells were trypsinized and plated for subsequent functional assays.

### Single-cell clone isolation and expansion

UPCI:SCC154 cells were passed through a round bottom polystyrene test tube, with cell strainer snap cap (Corning, Corning, NY, USA) to separate the cell suspension into single cells. Cells were diluted to 30 cells/ml; 100µl of the cell suspension (3 cells/well) were added to each well in a 96-well plate. 24 hours after attachment, the plate was scanned under a bright-field microscope; wells with one attached cell were recorded. Single cells were maintained in EMEM media with 10% FBS and 100 μg/ml penicillin and streptomycin until growth into a colony (>50 cells), which was expanded. Passage 0 was designated when cells reached 50-70% confluence in a 100 mm dish, then split into three; one fraction was seeded in a 100 mm plate (passage 1), the second fraction was frozen, and the third fraction was harvested for subsequent RT-PCR. This process was continued till passage 4.

### Reverse transcription-polymerase chain reaction (RT-PCR) and quantitative PCR (RT-qPCR)

RNA was extracted with the miRNA isolation kit (QIAGEN, Hilden, Germany, # 217084). cDNA was synthesized using SuperScript II reverse transcriptase (Invitrogen #18064014). PCR was performed in the Proflex base PCR System (Applied Biosystems, Waltham, MA, USA) using Taq DNA polymerase (New England Biolabs, Ipswich, MA, USA, # M0273S). PCR products were analyzed on an agarose gel (1.6-2.0% in Tris-Borate-EDTA buffer). Raw images of agarose gels for this study are in Fig. S10, S11, and S12. qPCR was performed with Power SYBR Green Master Mix (Applied Biosystems #43-676-59) on a StepOne Plus Real-Time PCR System (Applied Biosystems # 4376600). Data were analyzed by the relative quantification method with normalization to actin. Primers used for PCR and qPCR are listed in Table S5.

### Western blot (Immunoblot) Analysis

For immunoblotting of p53, Rb, p16, ECAD, VIM, and actin, cells were grown to 60-80% confluence, washed with phosphate-buffered saline, and lysed in 1% Nonidet P-40 lysis buffer containing 50 mmol/L Tris-HCl, pH 7.4, 200 mmol/L NaCl, 2 mmol/L MgCl₂, 10% glycerol, and protease inhibitors. Lysates were sonicated and centrifuged, and supernatants were collected. Protein concentration was determined using the Bio-Rad protein assay (Bio-Rad, Richmond, CA, USA). Equal amounts of protein were separated on 4–20% Tris-Glycine gels (Invitrogen) and transferred to nitrocellulose membranes.

Membranes were immunoblotted with primary antibodies against p53 (Cell Signaling Technology, Danvers, MA, USA; #2524; RRID: AB_331743), Rb (Cell Signaling Technology; #9309; RRID: AB_823629), p16 (Santa Cruz Biotechnology, Dallas, TX, USA; sc-1661; RRID: AB_628067), ECAD (BD Biosciences, Franklin Lakes, NJ, USA; #610182; RRID: AB_397581), VIM (Proteintech, Rosemont, IL, USA; #10366-1-AP; RRID: AB_2273020), and actin (BD Biosciences; #612656; RRID: AB_2289199).

For immunoblotting of HPV16 E6FL and E7, cells were lysed separately in radioimmunoprecipitation assay buffer and stored at −80°C overnight. Lysates were then thawed, centrifuged, and supernatants were collected. Equal amounts of protein were separated on 4-20% Tris-Glycine gels and transferred to nitrocellulose membranes that were immunoblotted with antibodies against HPV16 E6 (GeneTex, Irvine, CA, USA; GTX132686; RRID: AB_2886710) and HPV16 E7 (Santa Cruz Biotechnology; sc-65711; RRID: AB_831413). The E6FL band was quantified for E6FL protein analysis using ImageJ. GAPDH was used as the loading control for E6FL and E7 immunoblots.

After overnight incubation with primary antibodies at 4°C, membranes were incubated with horseradish peroxidase-conjugated goat anti-rabbit IgG or goat anti-mouse IgG secondary antibodies (Cell Signaling Technology; #7074 and #7076, respectively; RRID: AB_2099233 and AB_330924). Immunoreactive bands were visualized using SuperSignal West Pico chemiluminescent substrate (Pierce, #34577) on a ChemiDoc Imaging System (Bio-Rad; RRID: SCR_019684).

Band intensities were quantified using ImageJ software (National Institutes of Health, Bethesda, MD, USA; RRID: SCR_003070), expressed as arbitrary densitometric units, and normalized to the corresponding loading control, actin or GAPDH, as indicated. Raw immunoblot images are provided in Fig. S13.

### Proliferation assay

Cells were seeded at equal densities (3 × 10^4^ for UM-SCC-38, UM-SCC-47, UM-SCC104, UPCI:SCC090, and UPCI:SCC154) on a 24-well plate (4-6 replicate wells per condition). After overnight attachment, cell cultures were incubated in a BioTek BioSpa 8 Automated Incubator (Agilent Technologies, Santa Clara, CA, USA; RRID: SCR_019728), at 37°C and 5% CO_2_. Cell cultures were transferred from the BioSpa 8 incubator to a Cytation 5 cell imaging multimode reader (RRID: SCR_019732) every 12h. 4-6 high contrast brightfield images were captured per well and image pre-processing was performed to obtain cell number at each time point according to the manufacturer’s instructions.

### Wound healing assay

Cells were grown to 95-100% confluence in triplicate. After 24-48h, cells were treated with mitomycin C (Sigma #M4287) for 2h in serum-free media (0.5ug/ml for UPCI:SCC154; 1ug/ml for UM-SCC-38). After mitomycin C treatment, the serum-free media was replaced with complete media with 10% FBS. A scratch wound was created using BioTek Autoscratch tool (Agilent Technologies), and cell cultures were incubated in a BioSpa 8 incubator at 37°C and 5% CO_2_. Images of the wound were taken every 4-6h. The area of the wound at the indicated time point was quantified using ImageJ (RRID:SCR_003070; National Institute of Health). The percentage of wound closure was computed as: Percent wound closure = 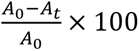, where *A*_0_is wound area at baseline and *A_t_*is wound area at time *t*.

### Fluoroblok migration assay

Cells were plated for 24-48h and then stained with CellTracker™ Green CMFDA Dye (Invitrogen # C7025) for 1.5h. Fluoroblok migration assays were performed as described (*108*). Briefly, labelled cells were trypsinized, resuspended in serum-free media, and added to the top chamber of the Fluoroblok transwell insert (8µm pore size, Corning #351152). Media with 10% FBS was added to the lower chamber of the transwell insert as a chemoattractant. Relative fluorescence units (RFU) at 492/517 nm were determined in a SpectraMax i3x Multi-Mode Microplate Reader (Molecular Devices, San Jose, CA, USA) using Softmax Pro (Molecular Devices; RRID: SCR_014240) software. Migration was quantified by normalizing to RFU of an uncoated transwell insert (RFU_m_) with the background RFU at 0h (RFU_b_).

### In vivo chick chorioallantoic membrane (CAM) and mouse experiments

Animal studies were performed in compliance with the National Institutes of Health guidelines for animal research and approved by the Institutional Animal Care and Use Committee of the University of Michigan.

For overexpression CAM studies, UM-SCC-38 stably transfected with E6*I, FL, FLm, or vector control were stained with CellTracker™ Green CMFDA Dye (Invitrogen # C7025) for 2h then grafted on the CAM (day 10), as described(*54*). For SSO CAM studies, UPCI:SCC154 transfected with SSO-1, SSO-2, SSO-5, or SSO-6 for 24h, were stained with CellTracker™ Green CMFDA Dye (Invitrogen # C7025) and grafted on the CAM (day 10)(*54*). CAMs were harvested at day 13, sectioned and embedded on slides by the University of Michigan Dental School Histology Core (RRID:SCR_026746). Slides were stained with H&E or immunostained with laminin B1 (Abcam, #ab229025; RRID: AB_3083735) and 4′,6-diamidino-2-phenylindole (DAPI) to highlight basement membrane and nuclei, respectively, as described (*54*). Fields of immunofluorescently stained CAM sections were imaged =using Leica Thunder Imager. Scoring of CAM basement membrane degradation was done for all eggs by an investigator blinded by groups according to the rubrics described in (*109*).

For mouse experiments, UM-SCC-38 cells stably transfected with PCD (control) or E6 isoforms were injected at the tongue tip in athymic nude mice (Crl:NU(NCr)-*Foxn1^nu^*; Charles River Laboratories; age 6-7 wk, male, n = 10-14/group) at 1x 10^6^ cells/mouse. Tumors were monitored for 21 days after which mice were euthanized. Tumors were harvested, processed and sectioned. Tissue sections (5μm, FFPE) were stained with H&E or cytokeratin AE1/AE3 mouse monoclonal antibody (EMD Millipore, IHCR2025–6) to highlight epithelium according to our previously established protocol (*110*).

Whole slide imaging of H&E (CAM and mice) and CK-stained slides (mice) was performed on a Leica Slide Scanner AT2. Fluorescence images for CAM slides were taken on a THUNDER imager (Leica Microsystems, Wetzlar, Germany; RRID: SCR_023794)]. The number and size of invasive tumor islands, and their distance from the main tumor (tumor bulk), were quantified from H&E images using Halo Image analysis platform v3.1 (Indica labs, Albuquerque, NM, USA) or Qupath v0.5.1(*111*).

### Immunofluorescence in vitro and in vivo

Cells were plated on poly-L-lysine coated glass coverslips for 24h, fixed with 4% paraformaldehyde (PFA) and permeabilized with 1% bovine serum albumin (BSA) and 0.2-0.3% TritonX-100 before incubation with primary antibodies. For co-detection of ECAD and VIM, cells were labelled with antibodies to detect VIM (Proteintech, #103661-1-AP; RRID: AB_2273020) and ECAD (BD Biosciences, #610182; RRID: AB_397581) diluted in 1% BSA and 0.3% TritonX-100 overnight at 4°C. Cells were then incubated for 2h at room temperature with goat anti-mouse Alexa Fluor 488 (Invitrogen #A11029; RRID: AB_2534088) and donkey anti-rabbit Alexa Fluor 568 (Invitrogen #A10042; RRID: AB_2534017) diluted in 1% BSA with 0.3% Triton X-100 to detect primary antibodies ECAD and VIM respectively. For detection of ECAD on Dyngo-4a-treated cells, cells were labelled with antibodies against ECAD (BD Biosciences, #610182; RRID: AB_397581) overnight at 4°C, before being incubated with donkey anti-mouse Alexa Fluor 594 (Invitrogen # A32744; RRID: AB_2762826) for 2h at room temperature. For co-detection of ECAD and Rab11 or EEA1, cells were incubated with antibodies against ECAD (BD Biosciences, #610182; RRID: AB_397581) and either EEA1 (Cell Signaling #3288; RRID: AB_2096811) or Rab11 (Cell Signaling #5589; RRID: AB_10693925) overnight at 4°C. Cells were then incubated with goat anti-rabbit Alexa Fluor 488 (Invitrogen # # A11034; RRID: AB_2576217) and donkey anti-mouse Alexa Fluor 594 (Invitrogen # A32744; RRID: AB_2762826) to detect EEA1 or Rab11, and ECAD respectively for 2h at room temperature.

Coverslips were mounted on slides using ProLong™ Gold Antifade mounting medium with DAPI (Thermo Fisher Scientific, Waltham MA, USA; #P36931). 4-6 fields per duplicate in each condition were imaged at 63x [with oil immersion using the THUNDER Imager (Leica). For co-detection of EEA1 or Rab11 and ECAD on 12 z-slices of 0.27um thick (default settings) were taken and degree of co-localization was quantified using Just Another Colocalization Plugin on ImageJ.

### Antibody Internalization

Internalization assay was performed based on an established protocol (*66*). Briefly UM-SCC-38 cells were seeded on poly-L-lysine-coated glass coverslips for 24h. The cell media was changed to binding buffer (Hanks’ balanced salt solution (HBSS) supplemented with 5mM Ca2+ and 50mg/ml BSA) for 1h at 37°C. Cells were then incubated with an antibody against the extracellular domain of ECAD (HECD-1, Invitrogen #13-1700; RRID: AB_2533003) diluted in binding buffer for 1h at 4°C to allow antibodies to bind to surface ECAD. Then cells were washed twice with PBS and incubated with pre-warmed binding buffer for 0 or 30min at 37°C to allow internalization of surface-bound ECAD antibodies. Endocytosis was stopped by immediately transferring cells to 4°C. Cells were washed twice with ice-cold PBS, followed three times with acid wash buffer (0.5M acetic acid with 0.5M NaCl in PBS; 5min per wash) to remove surface-bound ECAD. To visualize internalized ECAD, cells were fixed with 4% PFA, permeabilized with 0.2% TritonX-100, and incubated with CellTracker™ Green CMFDA Dye (Invitrogen # C7025) and secondary antibody donkey anti-mouse Alexa Fluor 594 (Invitrogen # A32744; RRID: AB_2762826) diluted in 1% BSA and 0.2% TritonX-100 for 2h at room temperature. Coverslips were mounted on slides using ProLong™ Gold Antifade with DAPI and imaged using Thunder Imager (63x oil immersion, 6 fields per duplicate per condition). Signal intensity of internalized ECAD was quantified using ImageJ. Images were split into channels and the green (CellTracker™ Green) and red (internalized ECAD) channels were analyzed. Thresholding was performed to create a binary mask on the green channel. The green channel mask was used to generate the cell area of interest to calculate total internalized ECAD (red) intensity within cells. Results were represented as internalized ECAD intensity over cell area. *Treatment with Dyngo-4a*

For endocytosis inhibition, cells were serum-starved for 2h before they were incubated with 50μm of Dyngo-4a (Abcam ab120689). DMSO was used as a vehicle control. Cells were either subjected to immunofluorescence at 24h after treatment, or wound healing assay after Dyngo-4a was added.

### Sample acquisition and pre-processing of patient samples

For UM18, UM67 (EGAS50000000893), TCGA and HVC, clinical and RNA sequencing data were downloaded from respective sources (*13, 21, 45, 106*). Sample acquisition and pre-processing of these datasets have been described in our previous published manuscripts.

For the UM20 cohort, library preparation was performed using the ribo-depletion method with the FastSelect kit and sequenced on the NovaSeq 6000 at the UM Advanced Genomics Core, resulting in 150bp paired-end reads. Raw reads were trimmed using Cutadapt (v3.4) and aligned to hg38. HPV read preprocessing for UM20 was performed as described in (*112*).

### Association tests between E6FL:E6_ALL_ influence scores and biological signatures

E6FL:E6_ALL_ influence scores were computed for the five cohorts as described (*30*). Integration status was defined as positive for patients having one or more expressed HPV insertion events, detected using SurVirus with default settings (*113*).

Pathway scores characterizing the tumor immune microenvironment, differentiation state and oxidative phosphorylation were computed across the four cohorts. These scores were derived as sample-wise pathway scores aggregated by the rank of gene expression levels, as previously described (*45*). Samples were ranked based on the expression levels of each gene. The ranks of all genes within the same pathway were summed for each sample, and the resulting values were centered by mean and scaled by standard deviation across samples to determine the final pathway scores. The following signature scores utilized corresponding Gene Ontology (GO) terms: T cell activation (GO: 0042110), B cell activation (GO:0042113), keratinocyte differentiation (GO: 0030216), mesenchymal cell differentiation (GO:0048762), cell matrix adhesion (GO:0007160), epithelial cell migration (GO:0010632), focal adhesion (KEGG: M7253), respiratory electron transport chain (GO:0022904), and oxidative phosphorylation (KEGG:M19540). For EMT (*114*) and p-EMT (*80*) scores, negatively regulated genes were ranked in descending order before summation and scaling. Positively regulated genes were ranked in ascending order before summation, scaling, and addition with negatively regulated genes. Cell type deconvolution utilized CIBERSORTx with single-cell RNAseq HNSCC as the reference signature matrix (*115*). Due to the distinct origins of the signature matrix and the mixture matrix obtained from single-cell RNAseq and bulk RNAseq platforms, respectively, B-mode batch correction was activated to eliminate technical variations. By default, quantile normalization and absolute mode were disabled. The number of permutations was set to 100, meeting the recommended minimum to achieve a robust deconvolution p-value. The resulting cell type deconvolution matrix included proportions of the following cell types for all input samples: T cells CD8, T cells CD4, fibroblasts, macrophages, B cells, malignant cells, mast cells, dendritic cells, myocytes, and endothelial cells.

To generate the association network in Fig. 6F, associations between the E6FL:E6_ALL_ influence score and all other clinical and molecular variables were calculated; only statistically significant (p<0.05) associations are included in the network For categorical associations, including associations with T stage, HPV integration status, and subtype, logistic regression was employed with cohort as a covariate. For all other associations (which are continuous), a linear model was used with cohort as a covariate. Cytoscape version 3.9.1 (*116, 117*) was utilized to create the network graph. Statistical details are in Table S3.

### Multiplex immunofluorescence staining and whole-slide imaging

Two non-overlapping cohorts were used to determine ECAD localization in patient tissues. Cohort 1 served as the discovery set, and Cohort 2 served as a non-overlapping internal validation set. Specimens with insufficient tissue were excluded from the analysis. In Cohort 1, eligible UM18 and UM67 specimens were divided into non-overlapping discovery and validation subsets, and the subsequently assembled UM20 specimens were assigned to the validation cohort. Clinical and epidemiological characteristics are provided in Table S2.

5μm sections of FFPE tumor tissue were mounted on Superfrost Plus charged glass slides by the University of Michigan Dental School Histology Core. For Cohort 1, one tissue section was stained with a mouse monoclonal antibody against cytokeratin AE1/AE3 (EMD Millipore, IHCR2025-6) to identify epithelial regions, following our previously established protocol (*110*). An adjacent section was stained for ECAD and VIM by multiplex immunofluorescence. Briefly, tissue sections were incubated at 55-60°C overnight, deparaffinized in xylene, and rehydrated through a graded ethanol series (100%, 95%, and 70%) followed by Milli-Q water. Endogenous peroxidase activity was quenched by incubation with 1% hydrogen peroxide in methanol for 30 min at room temperature. Antigen retrieval was performed in 10 mM sodium citrate dihydrate buffer (pH 6.0) at 92°C for 15 min. Slides were then incubated for 1 h at room temperature in blocking solution containing 5% goat and donkey serum and 0.3% Triton X-100.

Primary antibodies against VIM (Cell Signaling Technology, #5741; RRID: AB_10695459) and ECAD (BD Biosciences, #610182; RRID: AB_397581) were diluted in 1% bovine serum albumin containing 0.3% Triton X-100 and applied overnight at 4°C. The following day, ECAD and VIM were detected using goat anti-mouse Alexa Fluor 568 (Invitrogen, A11031; RRID: AB_144696) and goat anti-rabbit Alexa Fluor 647 (Abcam, ab150079; RRID: AB_2722623), respectively. Slides were coverslipped using ProLong Gold Antifade Mountant with DAPI.

For Cohort 2, ECAD and VIM staining were performed using the same protocol. After incubation with the corresponding secondary antibodies, the same sections were incubated with an Alexa Fluor 488-conjugated mouse anti-cytokeratin antibody (Novus Biologicals, NBP2-33200AF488) for 2.5 h in the dark. Slides were then counterstained with DAPI and coverslipped.

Whole-slide images from both cohorts were acquired using a Vectra Polaris imaging system by the University of Michigan Tissue and Molecular Pathology Shared Resource Core Facility (RRID:SCR_026803).

### ECAD localization score and inter-rater analysis

ECAD localization was evaluated in multiplex immunofluorescence-stained sections by two investigators, including a board-certified pathologist, who independently scored each specimen while blinded to clinical outcomes, the E6FL:E6_ALL_ influence score, and other molecular data. Before formal scoring, the raters jointly reviewed representative staining patterns to establish a common interpretation of the scoring categories. Formal scores were subsequently assigned independently, without knowledge of the other rater’s assessment.

Tumor regions were identified using cytokeratin staining and corresponding histomorphology. Only viable tumor cells were evaluated. Stromal tissue, inflammatory cells, non-neoplastic epithelium, necrotic regions, tissue folds, and regions with insufficient tissue or inadequate staining quality were excluded. For each specimen, the raters estimated the percentage of viable tumor cells exhibiting each of three ECAD localization patterns:

Score 1: absent or weak ECAD, with predominantly cytoplasmic localization;

Score 2: mixed membrane and cytoplasmic ECAD localization; and

Score 3: predominantly crisp, continuous ECAD localization at cell–cell junctions.

The percentages assigned to the three patterns were required to sum to 100%. ECAD-LS was calculated separately for each rater as:

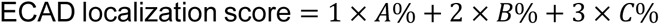

where A, B, and C denote the percentages of tumor cells in ECAD localization categories 1, 2, and 3, respectively. Scores therefore ranged from 100 to 300, with lower scores indicating greater cytoplasmic localization and higher scores indicating greater membrane localization. For downstream analyses, the final ECAD localization score for each sample was calculated as the average of the two raters’ scores.

Inter-rater agreement was assessed using the intraclass correlation coefficient (ICC) with 95% confidence intervals. ICCs were calculated separately for Cohort 1, Cohort 2, and the pooled cohort using a two-way random-effects model assessing absolute agreement. ICC values were interpreted as poor (<0.50), moderate (0.50-0.75), good (0.75-0.90), or excellent (>0.90) agreement.

### Quantification of total ECAD and VIM in patient specimens

Whole-slide fluorescence images were analyzed using InForm software v3.0 (Akoya Biosciences). For each slide, 4-8 regions of interest (ROIs) were marked and trained to segment ROIs into tumor, non-tumor and blank (no tissue) areas. Tumor classification was guided by cytokeratin staining and histomorphology. Following this, adaptive cell segmentation was trained using the same ROIs to segment cells into cytoplasm (VIM and cytokeratin), membrane (ECAD), and nucleus (DAPI). Stromal, inflammatory, necrotic, non-neoplastic epithelial, folded, and technically inadequate regions were excluded. Cell-level ECAD and VIM fluorescence intensities were extracted from identified tumor cells. For each patient, total ECAD and VIM expression was summarized as the mean whole-cell fluorescence intensity across all evaluable tumor cells. Image-analysis parameters were established by an investigator blinded to clinical outcomes or E6FL:E6_ALL_ influence scores.

### Survival analysis

For Kaplan-Meier analysis of E6FL:E6_ALL_ influence score, patients were classified as having either high or low E6FL:E6_ALL_ influence scores, based on the mean. Overall survival (OS) was defined as time in months from diagnosis to death and analyzed for 147 patients for whom outcome data is available (UM18, UM67, UM20 and TCGA; Table 1). Recurrence-free survival (RFS) was defined as the time in months from diagnosis to either a recurrence or death, whichever occurred first, and analyzed in 88 patients with available recurrence data (UM18, UM67 and UM20; Table 1). Using the R packages survival and survminer, Kaplan-Meier curves were created, and a p value was calculated using a univariate log-rank test for all 147 patients. A Cox proportional hazards regression model was used in 137 patients (10 removed due to missing data) after adjusting for age, overall stage, and cohort, and in 87 patients (1 removed due to missing data) for recurrence-free survival. Analysis was repeated after restricting patients with HPV16+ OPSCC. TNM staging for all samples was ascertained from medical chart data and reviewed by attending clinicians and converted to overall AJCC tumor stage 8^th^ edition.

For Kaplan-Meier analysis of the ECAD-LS, patients for Cohort 1 and Cohort 2 (Table S2) were dicohotomized according to the median ECAD-LS in Cohort 1 (195.0) as a locked cutoff. As a sensitivity analysis, Cohort 2 was also dichotomized using its own median ECAD-LS of 218.8. The two cohorts were additionally pooled and analyzed using either the pooled-cohort median of 205.0 or the Cohort 1 median of 195.0 as a locked cutoff. Kaplan–Meier curves were generated for RFS and OS, and survival distributions were compared using two-sided log-rank tests. When no events occurred in one ECAD-LS group, the conventional Cox hazard ratio was considered non-estimable and was not reported. Cox proportional hazards regression was performed in the pooled cohort to adjust for the age, overall stage and tissue cohort.

Because ECAD-LS is an intrinsically continuous measurement, we also evaluated it as a continuous variable in the pooled cohort using Cox proportional hazards regression. Multivariable models were adjusted for age at diagnosis, overall clinical stage, and tissue cohort. Associations of ECAD-LS with RFS and OS were evaluated separately.

Total ECAD and VIM fluorescence intensities were analyzed separately. Because staining and image processing were performed separately for the two tissue cohorts, total ECAD and VIM intensities were analyzed within each cohort. For survival analyses, patients were classified as having high or low total ECAD or VIM expression using the fluorescence intensity within each cohort as the cutoff. Kaplan–Meier curves were generated separately for Cohort 1 and Cohort 2, and associations with RFS and OS were assessed using two-sided log-rank tests.

### Statistical analysis

For all statistical analysis, a p value of less than 0.05 was considered statistically significant. Statistical comparisons between two groups were performed using unpaired t tests, while comparisons between more than two groups were with one-way ANOVA post-hoc Tukey test. Fisher’s Exact test was used to statistically compare the degree of laminin-stained basement membrane degradation in the CAM assay. Statistical tests for Kaplan-Meier curve and bioinformatics analysis are described above.

## Acknowledgements

This study makes use of data generated by Drs. Gillison, Symer and Akagi in the HPV Virome Consortium, formerly at The Ohio State University Comprehensive Cancer Center and now at University of Texas MD Anderson Cancer Center. Research reported in this publication was supported by the National Institute of Health/National Cancer Institute P30CA046592 (University of Michigan Cancer Center Shared Resources: Cancer Data Sciences) and National Institute of Health/National Cancer Institute (Proteogenomics of Cancer Training Program). We are grateful for services provided by University of Michigan Research Histology and Immunohistochemistry Core and School of Dentistry Histology Core. We are grateful to Dr Timothy Frankel for providing use of InForm software version 3.0 for morphometric analysis.

## Funding

National Institute of Health/National Cancer Institute CA250214 (NJD/MAS/LR) National Institute of Health/National Institute of Dental and Craniofacial Research R35 DE027551 (NJD)

Postdoctoral Fellows Small Grant Program of the University of Michigan Rogel Cancer Center (YXL).

## Author contributions

Conceptualization (NJD, YXL, ML, LSR, MAS)

Data curation (BFG, SL, MAS, SX, LR)

Formal analysis (YXL, BFG, AF, SL, JC, QL, VB)

Funding acquisition (NJD, MAS, LSR, YXL)

Investigation (YXL, ML, BFG, SL, JC, QL, JBH, GTW, MAS, NJD)

Methodology (YXL, ML, BFG, HK, RH, TL, LGM, MCM, LSR, MAS, AW, NJD)

Supervision (NJD, MAS, LSR, VB)

Validation (YXL, ML, BFG, AF, JC, AW, QL)

Visualization (YXL, ML, BFG, NJD, JC, AW, QL)

Writing-original draft (YXL, ML, BFG, MLM, MAS, NJD)

Writing-review and editing (all authors).

Corresponding author and lead contact (NJD)

## Competing interests

The authors declare no competing interests

## Data availability

All data is present in this paper or Supplementary Information. Information of UM18 and HVC cohorts were provided in GEO #GSE74956 and European Genome-phenome Archive EGAD00001004366 respectively. Information of UM67 cohort dataset is in European Genome-phenome Archive EGAD50000001306. We are in the process submitting the information from UM20 dataset under the same study as UM67 (EGAS50000000893).

## Supplementary Figures and Tables

**Fig. S1:**
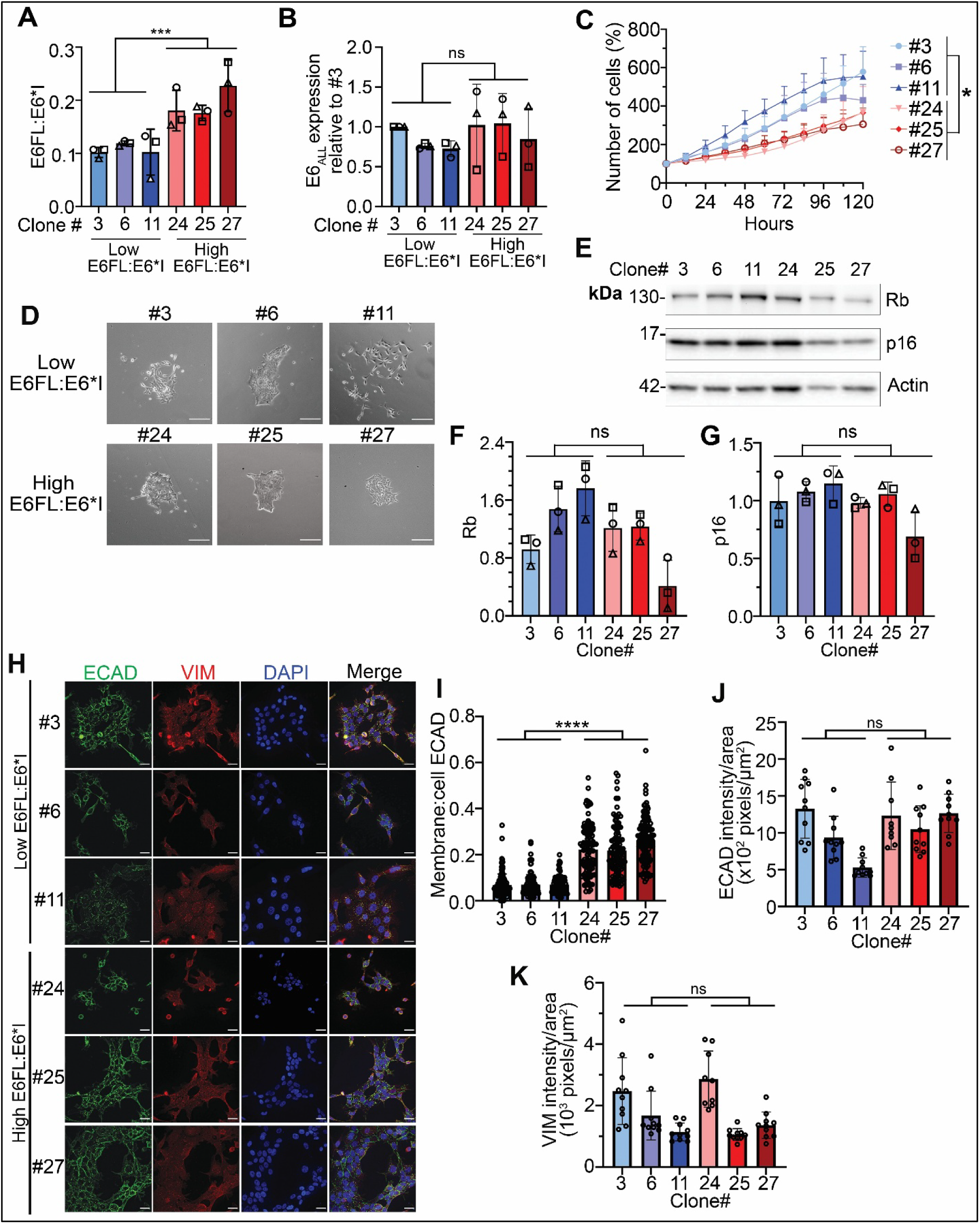
Validation and functional phenotypes of UPCI:SCC154 single-cell clones. (A-B) Validation of E6FL:E6*I (A) and E6_ALL_ (B) transcript expression in selected clones. E6_ALL_ was normalized with 18S mRNA expression and expressed as a fold change relative to clone #3. (C) Proliferation assay performed on UPCI:SCC154 single-cell clones. Viable cells were quantified and expressed as percent (%) of 0h. Three independent experiments were quantified, with 4 replicates per experiment. (D) Phase contrast images of colonies after 15 days of growth. Scale bar = 500μm (E) Representative immunoblot showing Rb and p16 expression from three independent experiments. Actin was used as the loading control. (F&G) Densitometric quantification of Rb (F) and p16 (G), normalized to actin. Each shape shows one experiment (three). ns= not significant (unpaired t test). (H) Individual channels for immunofluorescence staining of DAPI (blue), E cadherin (ECAD, green) and vimentin (VIM, red) in UPCI:SCC154 single-cell clones. (I) Membrane:cell ECAD was quantified from (H). Each dot represents one cell. At least 120 cells were quantified. (J&K) ECAD (J) and VIM (K) was quantified per field and normalized by total cell area in the field. Each dot represents one field. For all graphs, data are represented as mean ± SD. * p<0.05; ns= not significant (One-way ANOVA post-hoc Tukey)

**Fig. S2:**
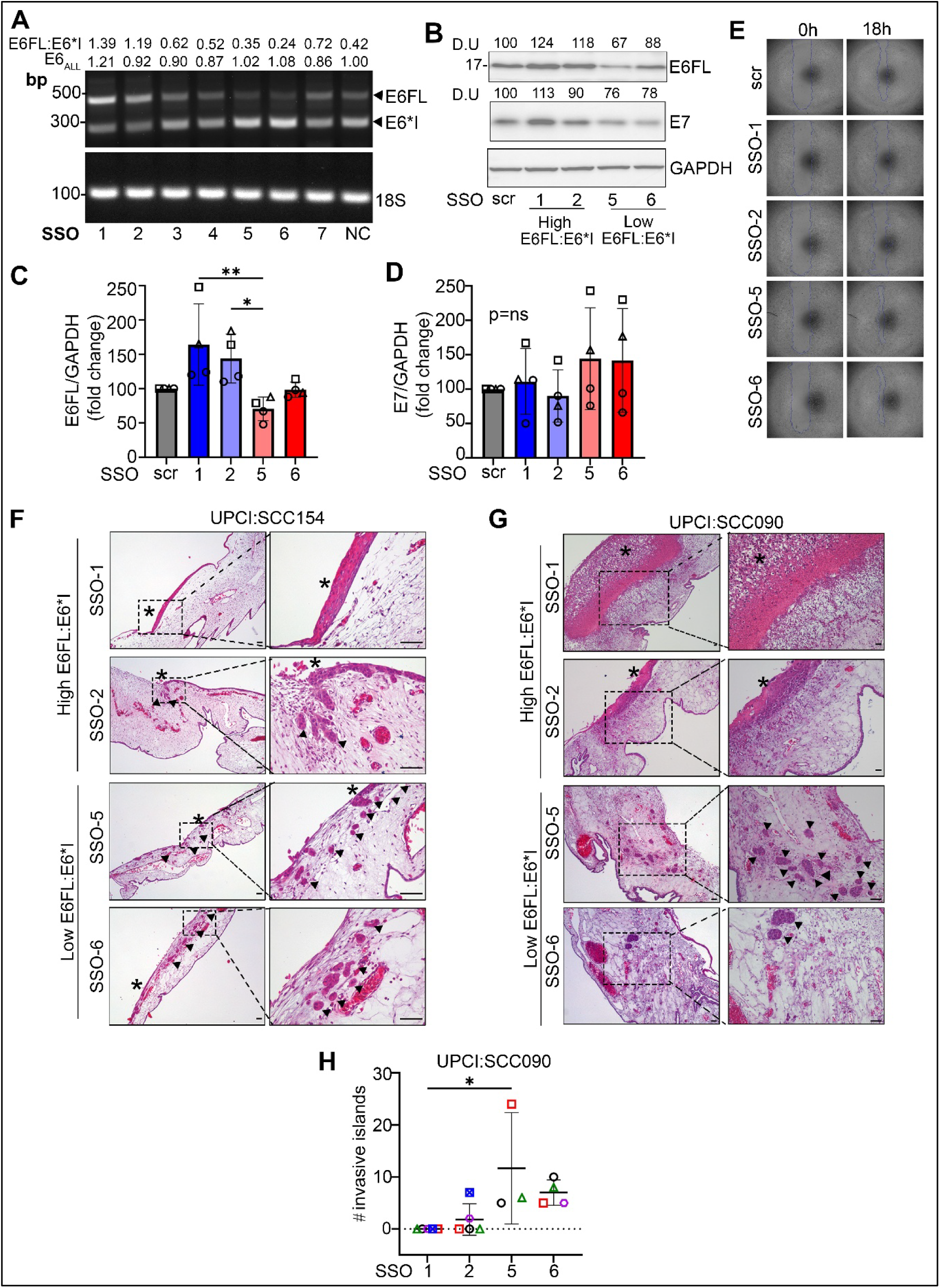
Functional studies performed in HPV+ OPSCC cell lines transfected with SSOs. (A) Representative agarose gel from 3 independent experiments validating expression of E6 isoforms in cDNA of UPCI:SCC090 transiently transfected with SSOs. NC represents non-targeting control, scr represents scrambled SSO control. E6FL:E6*I ratios were quantified by densitometry. E6_ALL_ transcript expression was quantified by summing E6FL and E6*I and normalizing by 18S of the same sample, then expressed as a fold change relative to NC (B) Representative immunoblot of E6FL and E7 in UPCI:SCC154 transfected with SSOs. GAPDH is used as a loading control (C&D). Densitometry of E6FL (C) and E7 (D) of 4 independent experiments including (B) (E) Wound healing images of UPCI:SCC154 transfected with SSOs. Quantification is in Fig.4E (F-G) Representative H&E images from CAM assays performed in UPCI:SCC154 (F) and UPCI:SCC090 (G) transfected with SSOsArrows represent example of invasive islands that detach from the main tumor bulk. Asterisk represents the main tumor. In the images showing UPCI:SCC090 transfected with SSO-1 and SSO-2, no invasive island was identified. In images showing CAM from UPCI:SCC090 transfected with SSO-5 and SSO-6, the main tumor bulk could not be identified. (H) Number of invasive islands detached from main tumor in CAM with UPCI:SCC090 injected with SSOs. For all graphs, data are represented as mean ± SD. * p<0.05; ** p<0.01, ns= not significant (One-way ANOVA post-hoc Tukey)

**Fig. S3:**
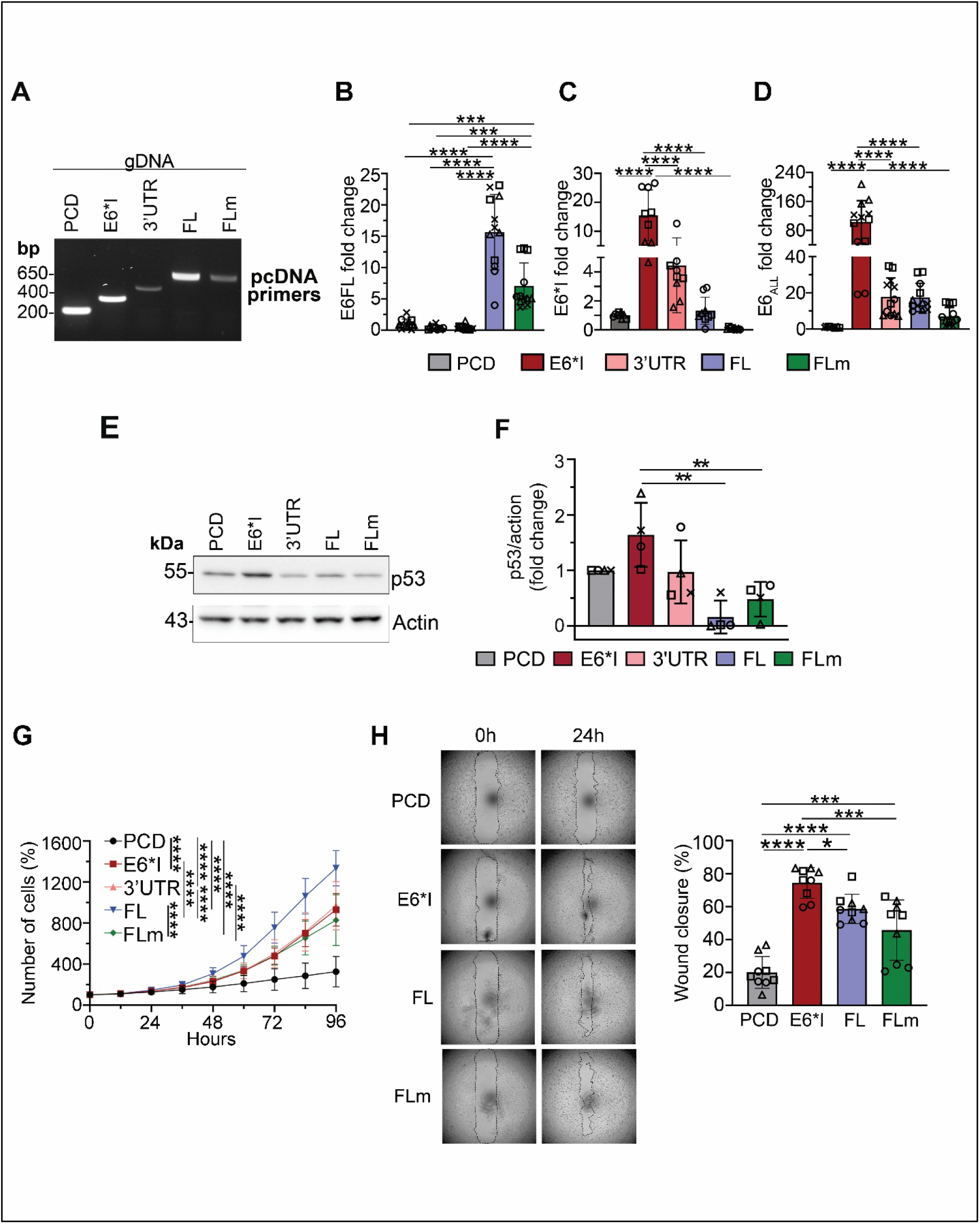
Overexpression of HPV16 E6 isoforms in HPV-negative OPSCC cell line, UM-SCC-38. (A) Agarose gel showing overexpression of E6 isoforms in genomic DNA with PCR using primers against pcDNA vector. Expected size: PCD= 198 bp; E6*I=356 bp; E6*I 3’UTR (3’UTR)=497 bp; FL/FLm=679 bp) (B-D) RT-qPCR of cell lines using primers against full-length E6 (E6FL) (B), E6*I (C) and all E6 isoforms (E6_ALL_) (D). Each shape represents an independent experiment, with three replicates per experiment. Transcript expression was represented as fold change relative to PCD control. (E) Representative immunoblot for p53, the downstream target of E6, in lysates. Actin was used as a loading control. (F) Densitometry for p53 was normalized to actin and expressed relative to PCD control from (C) and three other experiments (total four experiments). Each shape represents an independent experiment (four). Data is represented as mean ± SD. (G) Proliferation assay performed on stable cells. Number of cells were quantified at indicated time points and normalized as a percent (%) to 0h. (H) Representative images of wound healing assay of stable cells. Wound closure is represented as the percent of wound closure relative to 0h. Each shape is an independent experiment; same shapes represent replicates of the same experiment. One-way ANOVA with post-hoc Tukey test was performed for all assays. **p<0.01, ***p<0.001, ****p<0.0001

**Figure S4.**
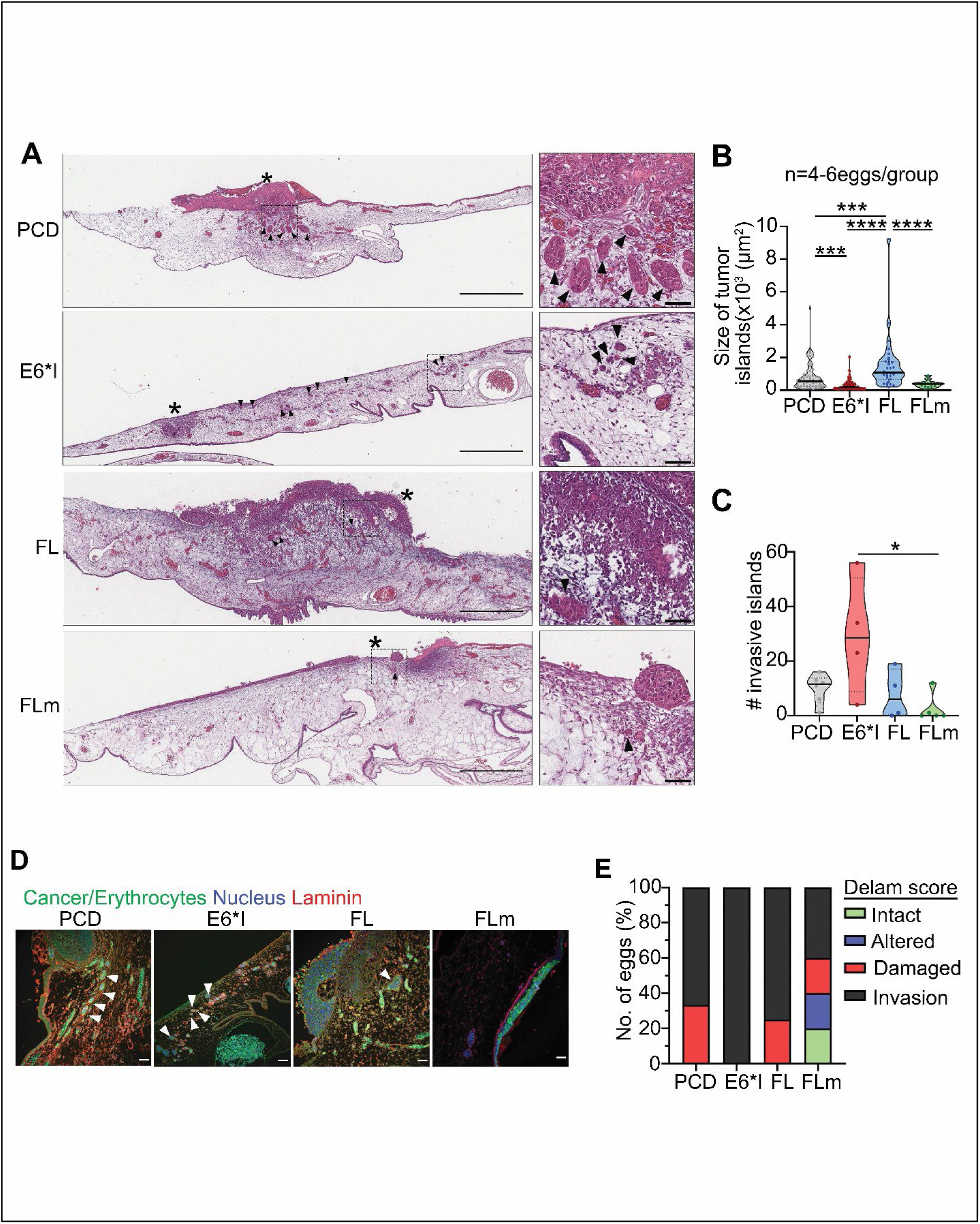
CAM assay performed in UM-SCC-38 overexpressing E6 isoforms. (A) UM-SCC-38 labeled with CellTracker™ Green CMFDA Dye were seeded on the upper CAM that was harvested 3 days later, sectioned and stained with hematoxylin and eosin (H&E). Whole-slide imaging of CAM sections was performed. Invasive tumor islands are indicated by black arrowheads. Tumor bulk is indicated by an asterisk (*). Right panel shows enlarged images from boxed areas in left panels. Scale bar= 500μm (left), 50μm (right). (B&C) Size of invasive islands (B) and number of invasive islands (C) were quantified from the entire CAM section shown in (A). N=4-6 eggs/group. Each point in (B) and (C) represents an invasive tumor island; each point in (D) represents a different egg. Data are represented as mean ± SD. *p<0.05, ***p<0.001, ****p<0.0001 (One-way ANOVA post-hoc Tukey test). (D) Representative images of upper CAM stained for laminin (red; basement membrane) and DAPI (blue; nucleus). Green signals highlights fluorescently-labelled tumor cells with some non-specific staining observed in erythrocytes. Arrowheads indicate invasive tumor islands. Scale bar =50μm. (E) Basement membrane degradation of immunofluorescence images including (D) was scored according to the CAM-Delam scoring system reported by Palaniappan et al. (2020) (Fisher’s Exact Score; p=ns).

**Fig. S5:**
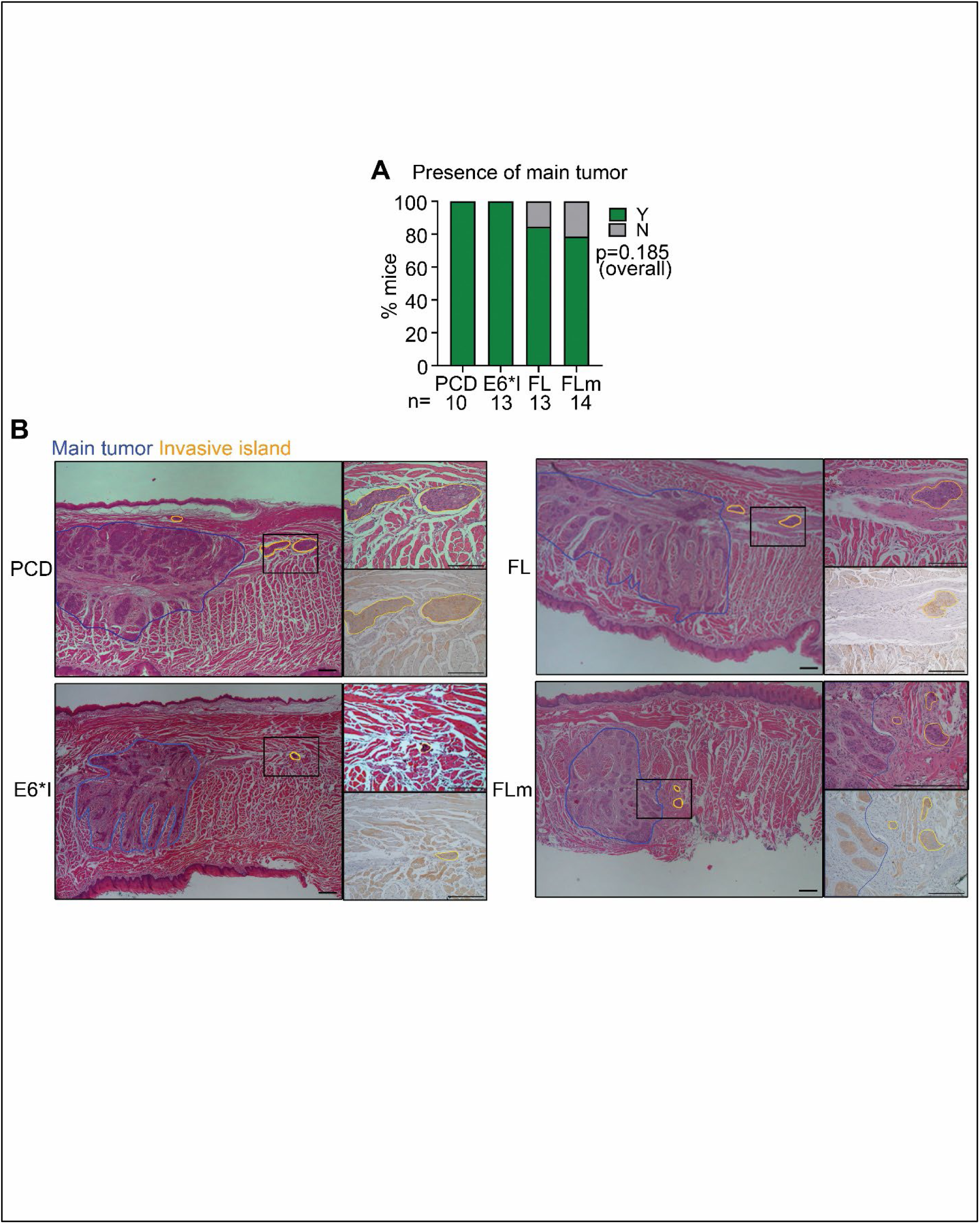
Mouse tumors. (A) Percentage (%) of mice showing presence of main tumor (Y=yes, N=no). (B) H&E and CK-stained images of main tumor bulk (blue outline) and invasive islands (yellow outline) in mouse tongue. Right panels are high magnification H&E and CK-stained images of boxed area in left panels. Scale bar =200μm in both panels

**Fig. S6:**
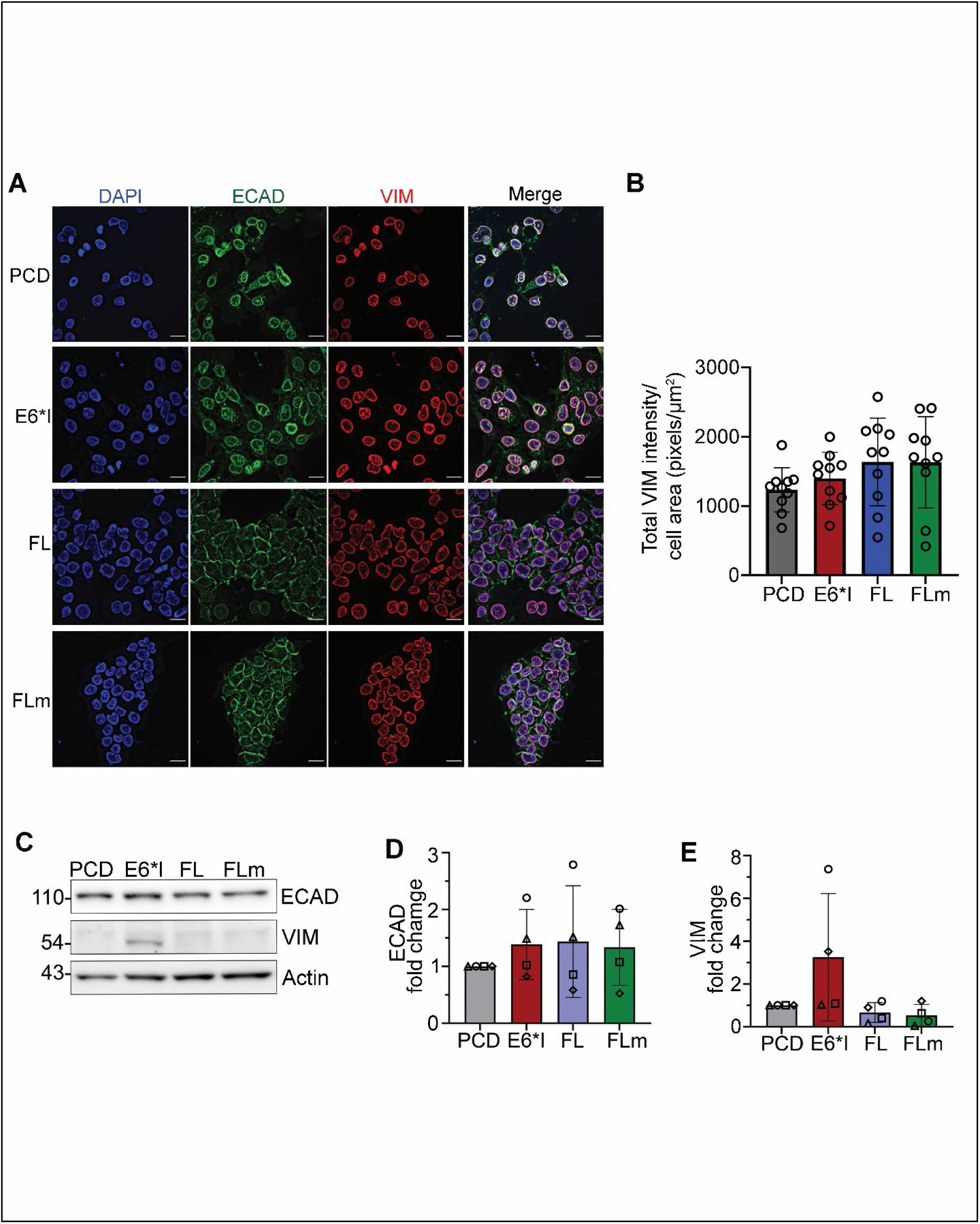
EMT markers in UM-SCC-38 overexpressing E6 isoforms. Individual channels of Immunofluorescence of DAPI (blue), E cadherin (ECAD, green) and vimentin (VIM, red) in UM-SCC-38 stably overexpressing E6 isoforms. Scale bar =100 µM. Quantification of total VIM intensity normalized to cell area over 10 independent fields including (A) (one data point represents one field). One-way ANOVA post-hoc Tukey comparison; no statistical significance. Representative Immunoblot of ECAD and VIM. Actin was a loading control. (D&E) Densitometry of immunoblots for ECAD (D) and VIM (E) normalized to actin and expressed relative to PCD. Each data point is an independent experiment. For all graphs in this figure, data are represented as mean ± SD (one-way ANOVA post-hoc Tukey test; no asterisk=not significant)

**Fig. S7:**
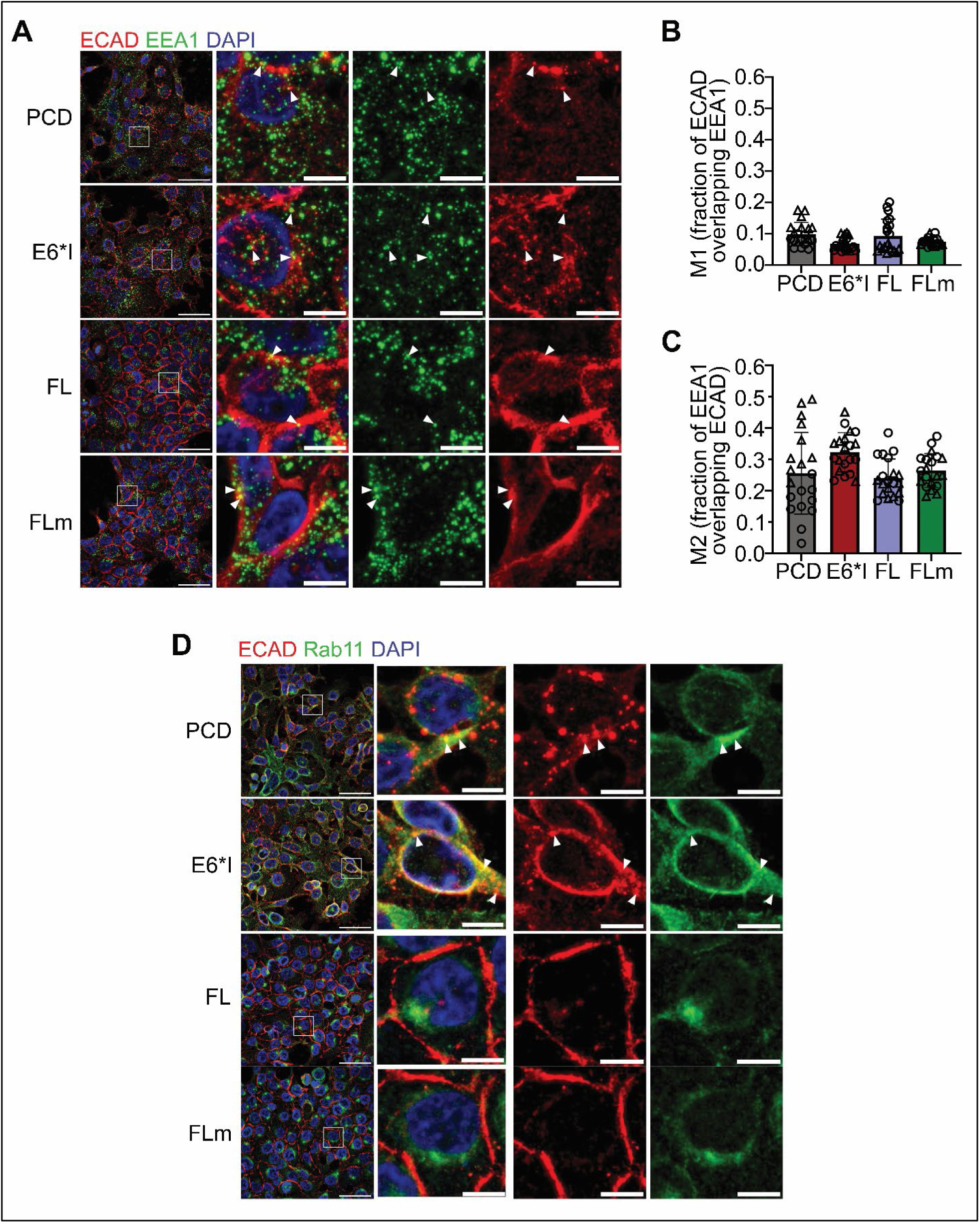
Co-localization of ECAD with markers for endocytosis vesicles, EEA1 and Rab11. (A) Maximum projection images of ECAD (red) and EEA1 (green) in UM-SCC-38 expressing E6 isoforms or vector control (PCD). Second panel shows merged image of boxed area in first panel. Third and fourth panels show individual channels for ECAD (red) and EEA1 (green), respectively. Arrowheads show co-localization of ECAD and EEA1. Scale bar= 50μm (B&C) Co-localization of ECAD and EEA1 was quantified from 12 z slices, 5 fields/duplicate. Manders’ coefficients, i.e. fraction of ECAD overlapping EEA1 (B) and fraction of EEA1 overlapping ECAD (C), were quantified using ImageJ JaCoP plugin. Each point represents a different field. Data are represented as mean ± SD. *p<0.05, one-way ANOVA post-hoc Tukey) (D) Maximum projection images of ECAD (red) and Rab11 (green) in UM-SCC-38 with PCD or E6 isoforms. Second panel shows enlarged image of boxed area in first panel. Third and fourth panels are ECAD (red) and Rab11 (green), respectively. DAPI (blue) = nucleus. Scale bar = 50μm (left) and 10μm (right).

**Figure S8.**
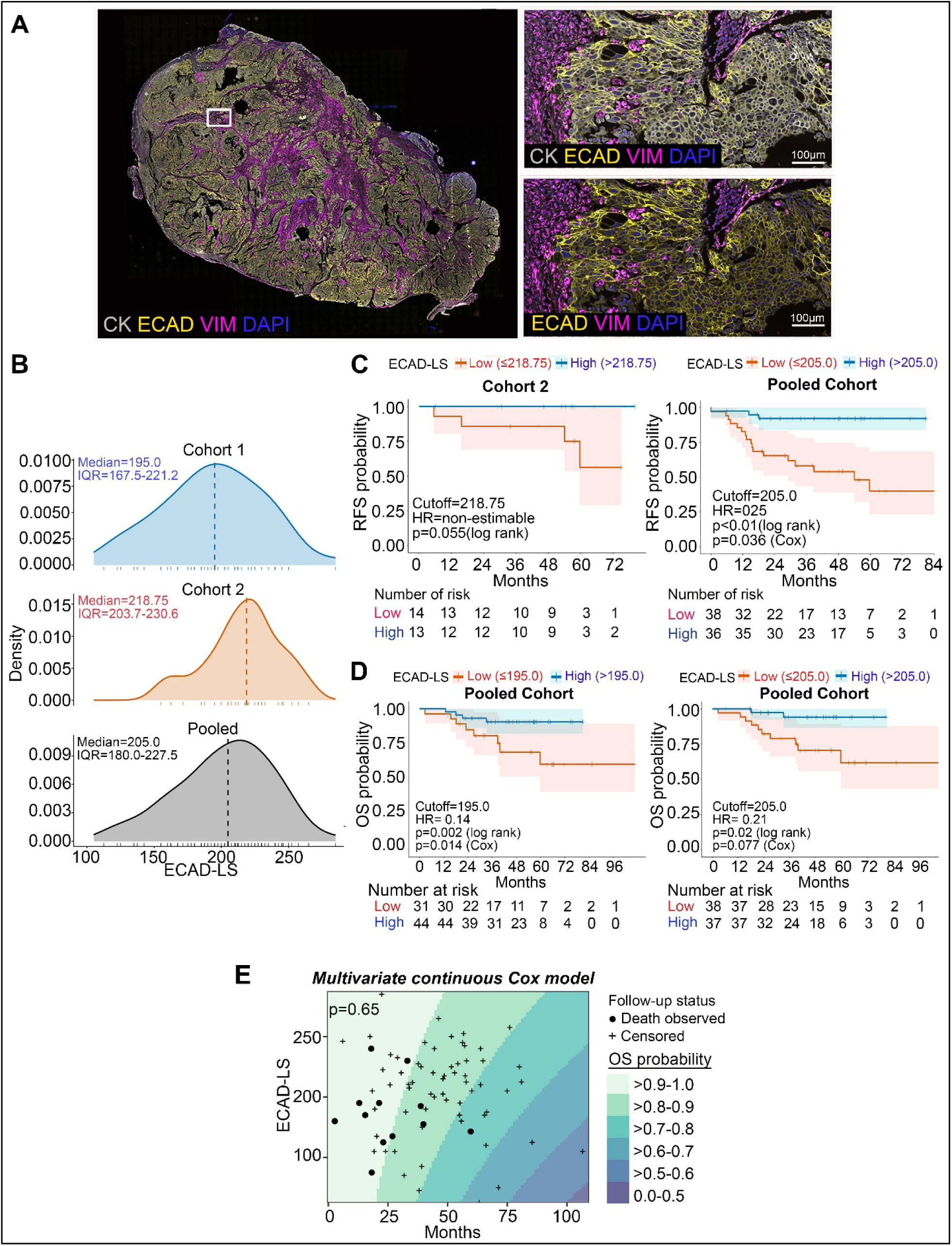
ECAD localization and clinical outcomes in HPV+ OPSCC. (A) Representative whole-slide (left) and enlarged (right) multiplex immunofluorescence images showing CK (grey), ECAD (yellow), VIM (magenta) and DAPI (blue). Panels on left are enlarged areas from boxed area from right. (B) Distributions of ECAD localization scores (ECAD-LS) in cohorts 1 and 2 and the pooled cohort; dashed lines indicate medians and tick marks in x axis of plots indicate individual tumors. (C) Recurrence-free survival (RFS) in cohort 2 and the pooled cohort according to low versus high ECAD-LS, using the cohort 1 median (195.0) as the cutoff. (D) Overall survival (OS) in the pooled cohort using the indicated ECAD-LS cutoffs. Left panel is stratified by cohort 1 median as a locked cutoff (195.0) and Right panel is stratified by pooled median (205.0). In D and E, shaded bands represent 95% confidence intervals, tick marks indicate censored observations, and numbers at risk are shown below the plots. (E) Multivariable continuous Cox model of ECAD-LS and OS; background colors indicate predicted OS-probability intervals, filled circles indicate deaths, and plus signs indicate censored observations.

**Figure S9.**
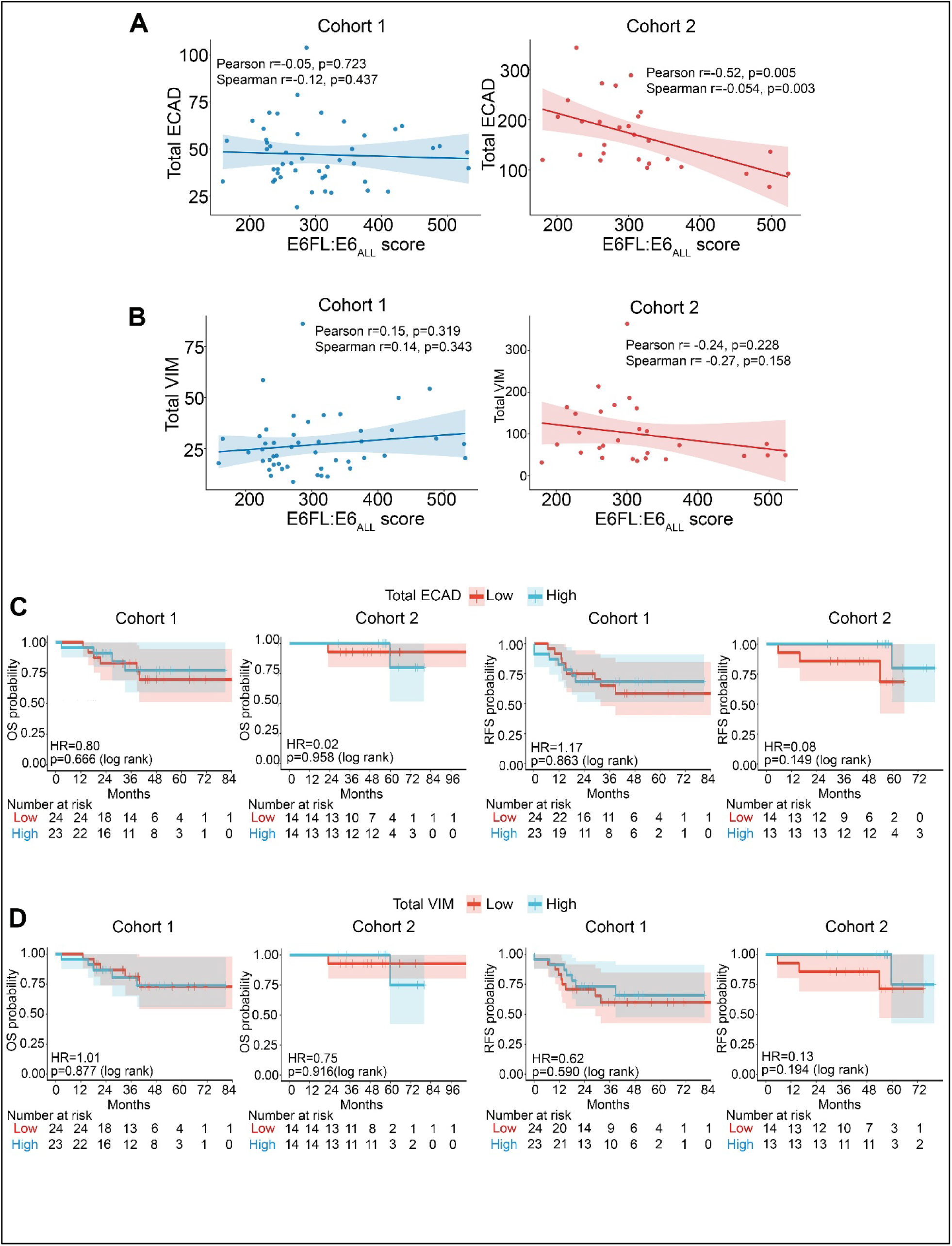
Associations of total ECAD and VIM expression with E6FL:E6ALL scores and clinical outcomes. (A&B) Associations of the E6FL:E6ALL score with total ECAD (A) and total vimentin (B) in cohorts 1 and 2. Lines indicate linear regression fits, and shaded bands indicate 95% confidence intervals. (C&D) Kaplan–Meier estimates of overall survival (C) and recurrence-free survival (D) according to low versus high total ECAD or VIM expression, dichotomized at the median within each cohort. Shaded bands indicate 95% confidence intervals, tick marks indicate censored observations, and numbers at risk are shown below each plot.

**Fig. S10:**
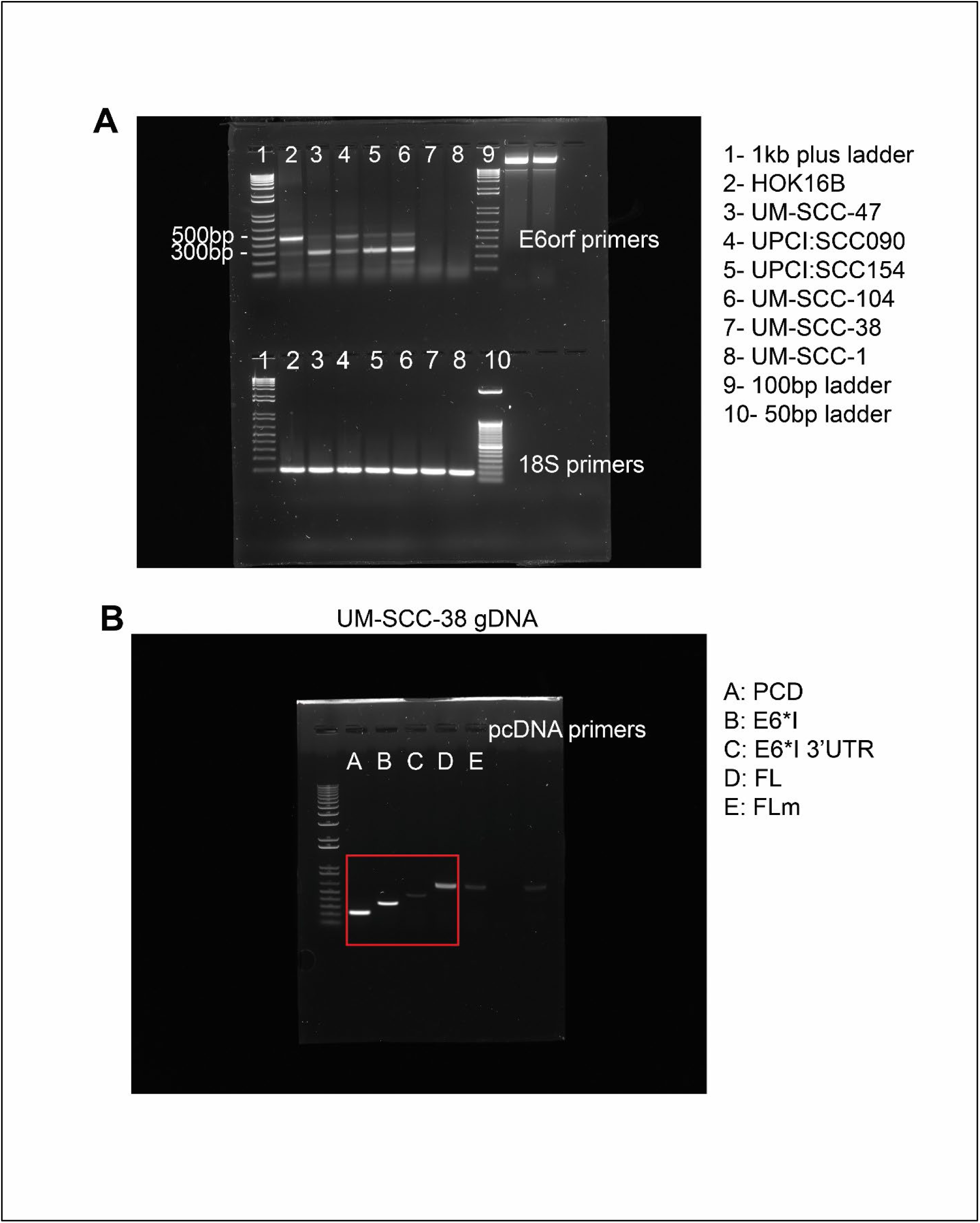
Uncropped images of agarose gel for RT-PCR experiments. (A) Uncropped raw image for Fig. 1B. (B) Uncropped raw image for Fig. S3A. Red box annotates the bands used in this study.

**Fig. S11:**
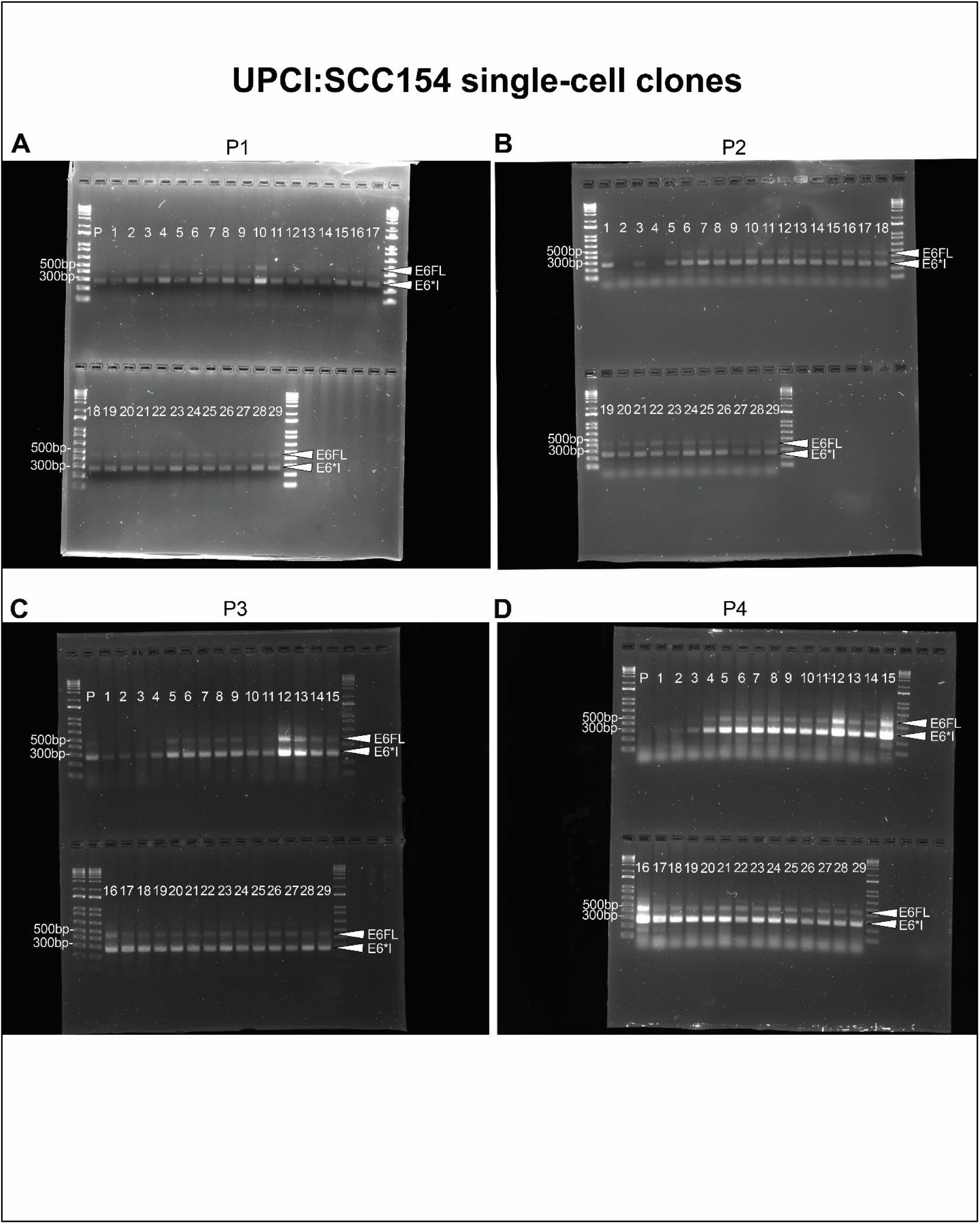
Raw data for screening of UPCI:SCC154SC clones in Fig. 2B. 29 clones were isolated from single cells from UPCI:SCC154 parent cell line. RT-PCR was performed using E6orf primers and PCR products were electrophoresed on an agarose gel. E6FL:(E6FL+E6*I) ratio was expressed as densitometric units of E6FL divided by the sum of densitometric units of E6FL and E6*I. Four consecutive passages (P1 - P4) were screened. P= UPCI:SCC154 parent cell line.

**Fig. S12:**
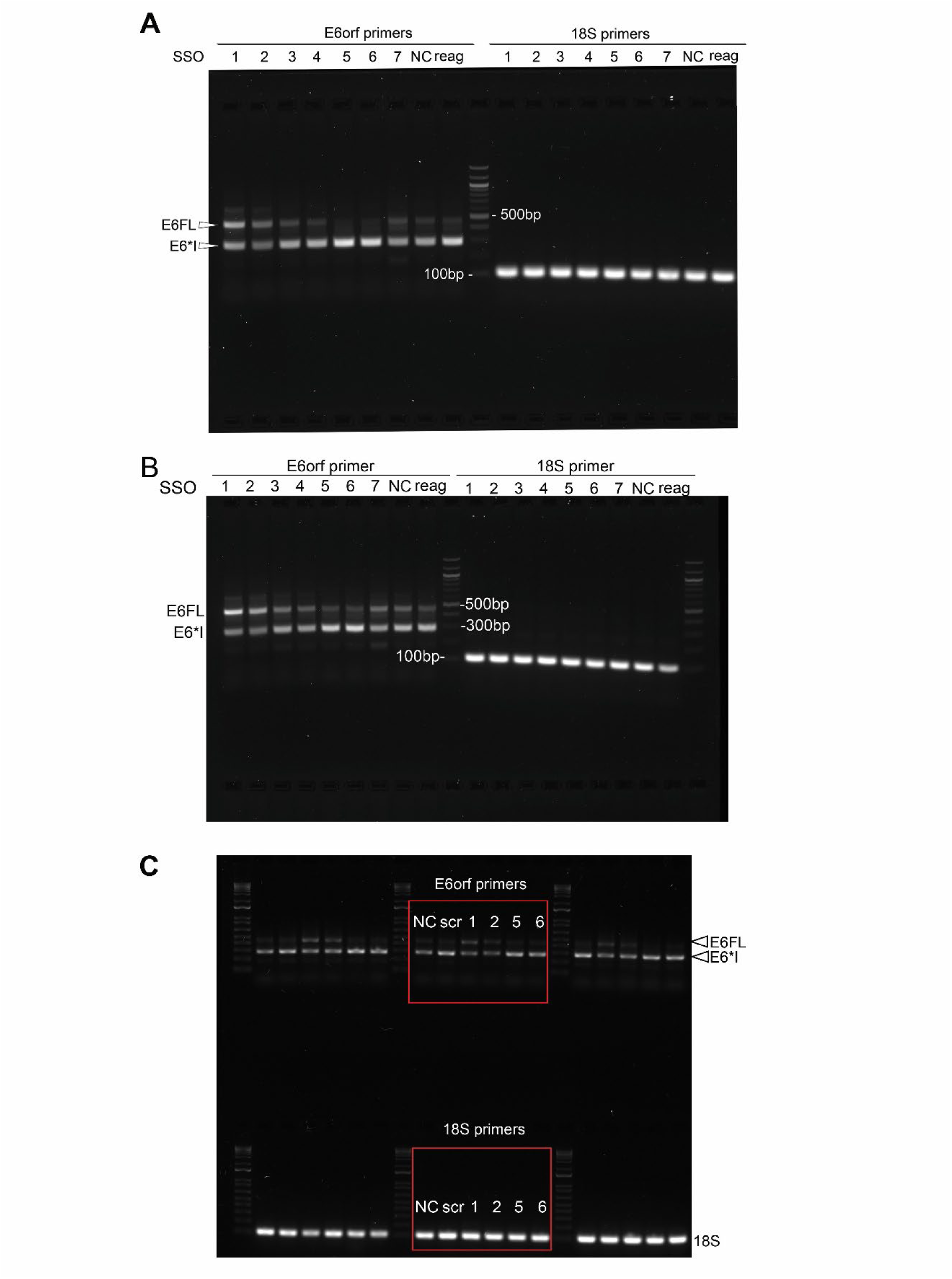
Raw data for screening of SSO experiments. (A) Raw data for Fig. 3B. (B) Fig. S2A (C) Fig. 3D

**Fig. S13:**
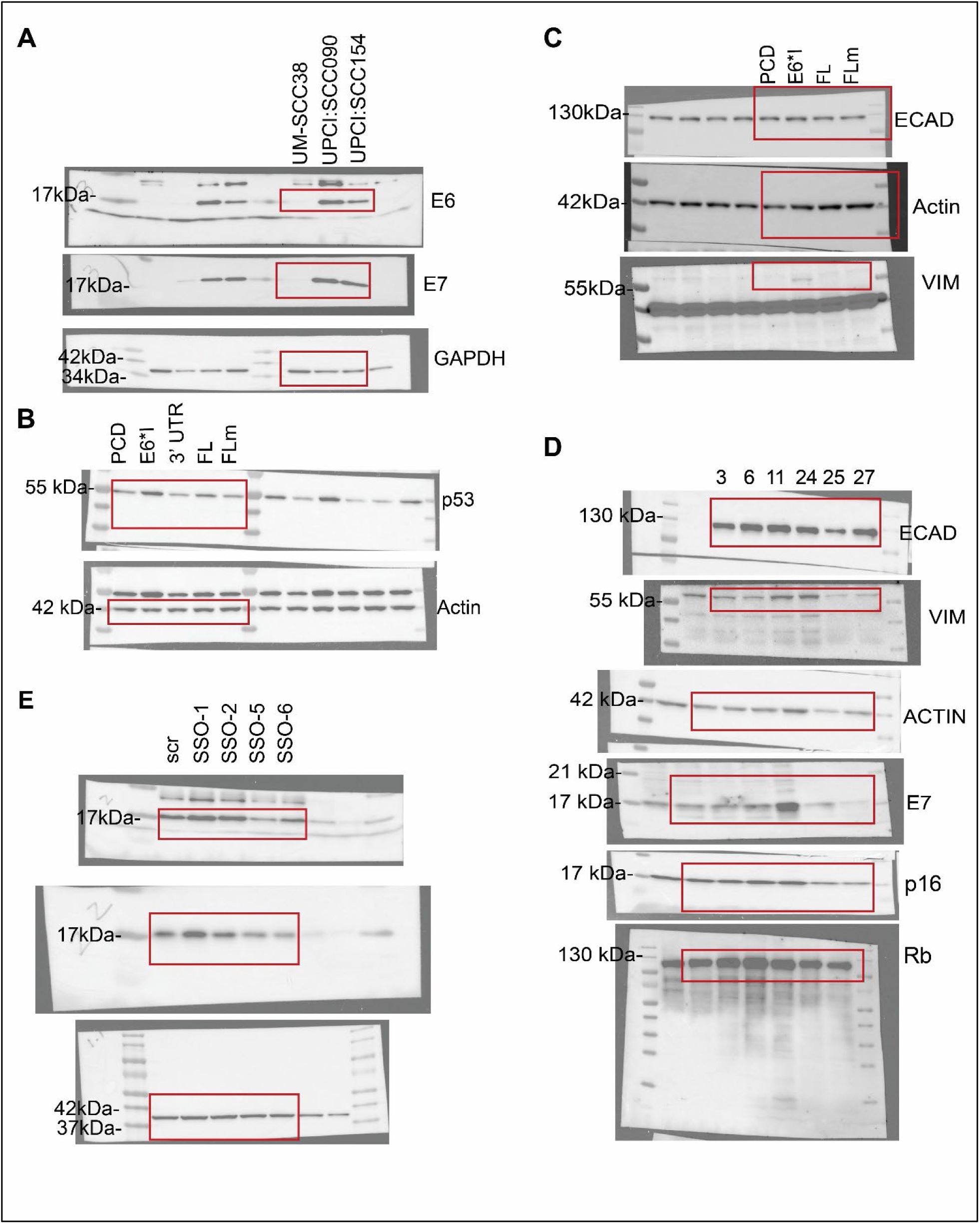
Raw immunoblots used in study. Raw data for (A) Fig. 1J; (B) Fig. S2E; (C) Fig. S6C; (D) Fig. S1E; (E) Fig. S2B

**Table S1.**
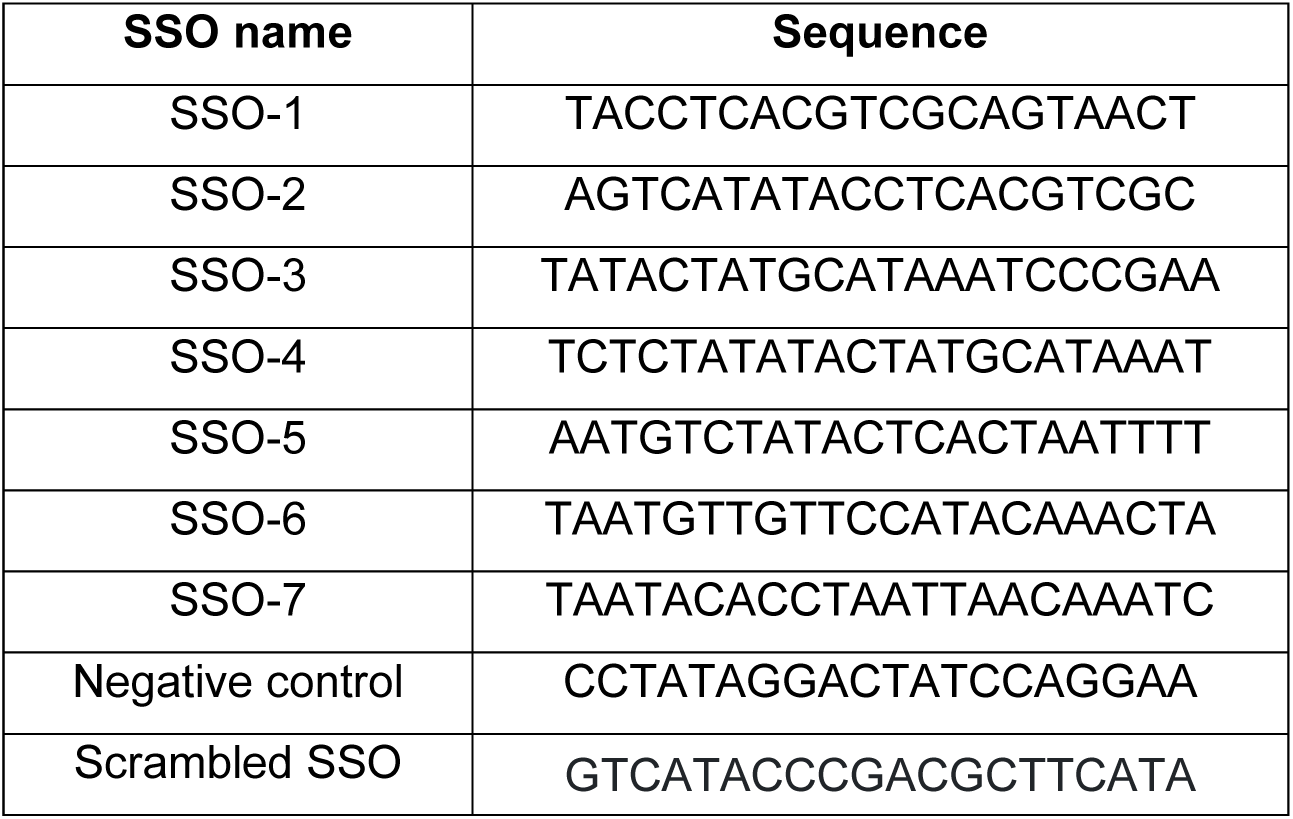
SSOs and sequences.

**Table S2.**
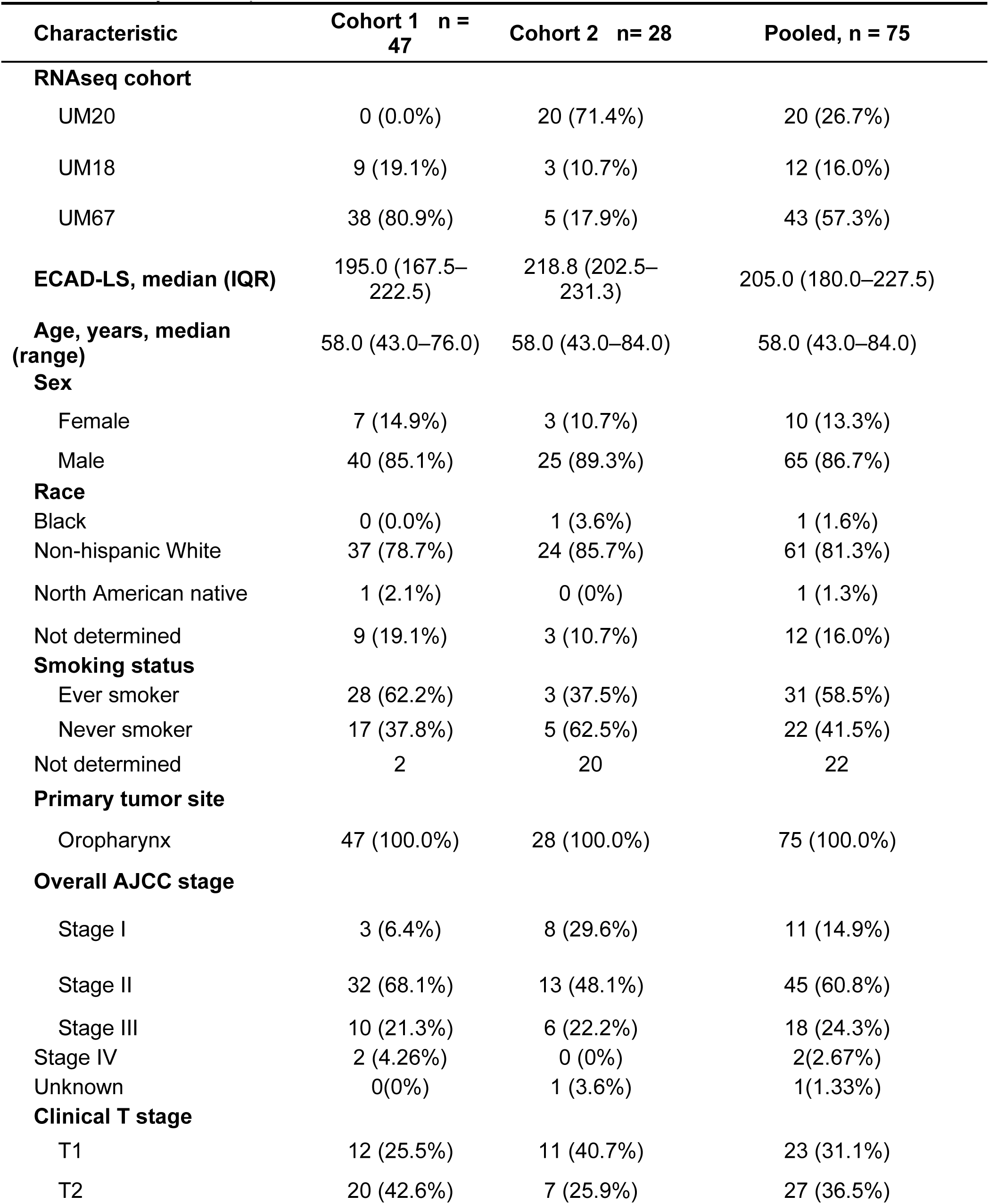
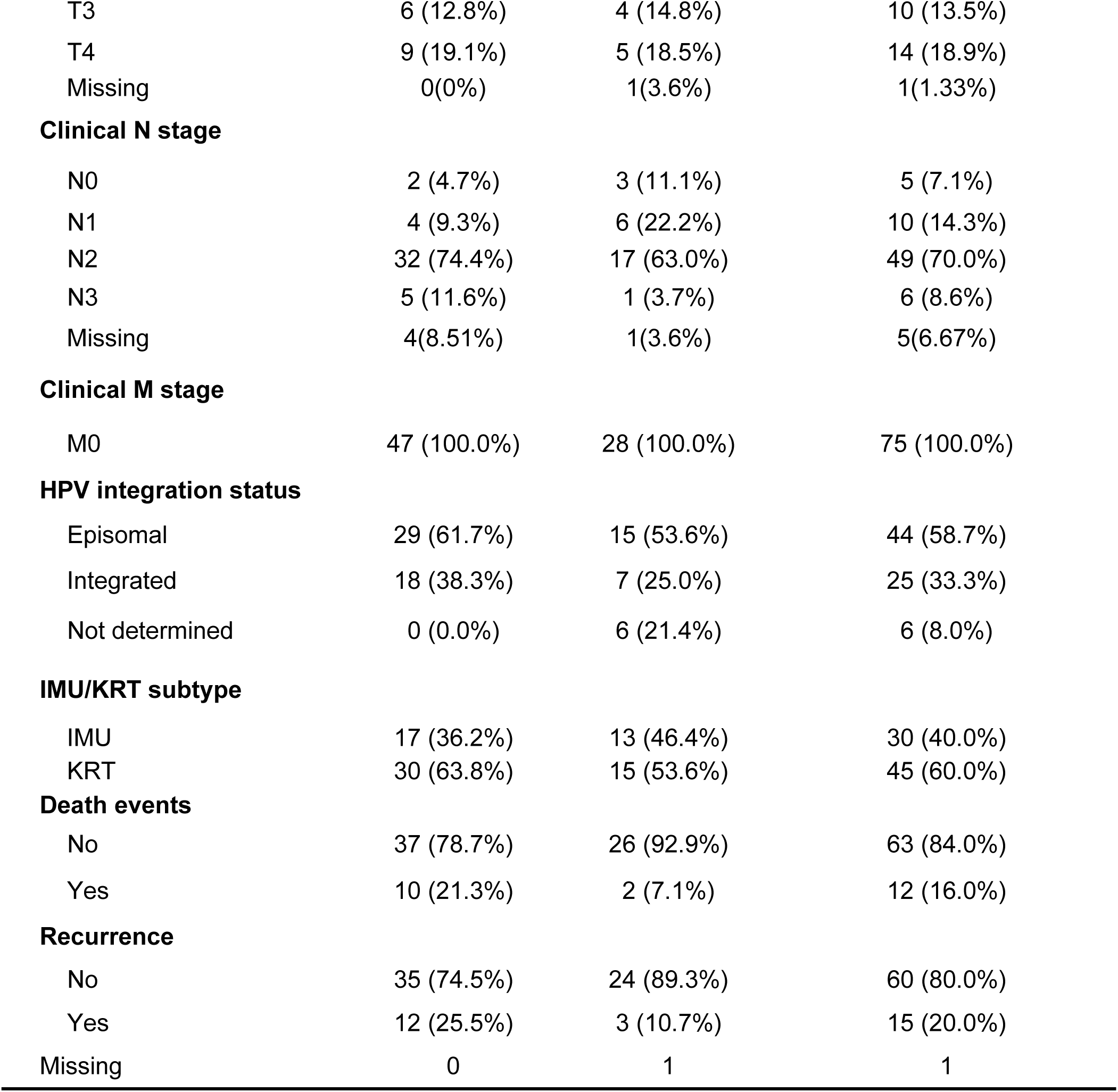
Clinical summary for multiplexed immunofluorescence cohort.

**Table S3.**
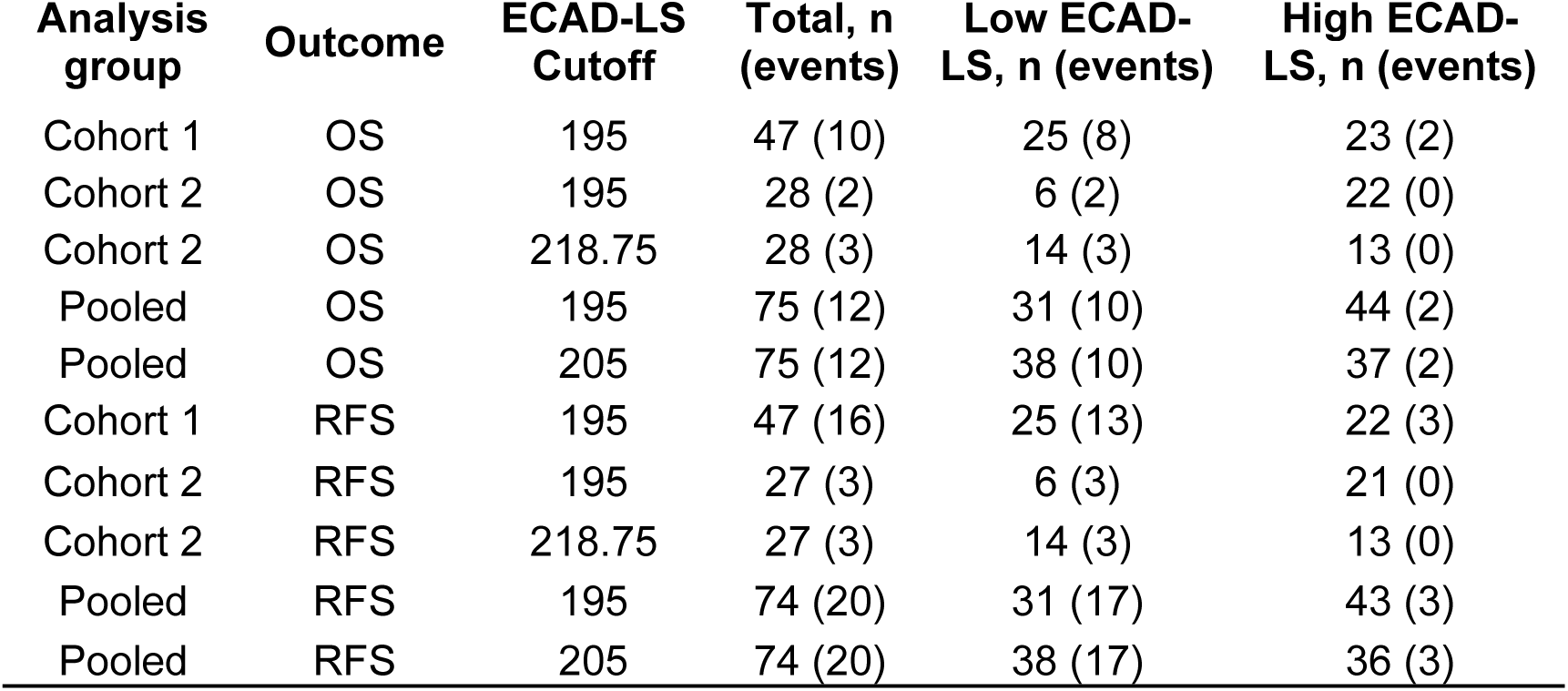
Cutoff and number of events for overall survival (OS) and recurrence-free survival (RFS) analysis for multiplexed immunofluorescence.

**Table S4.**
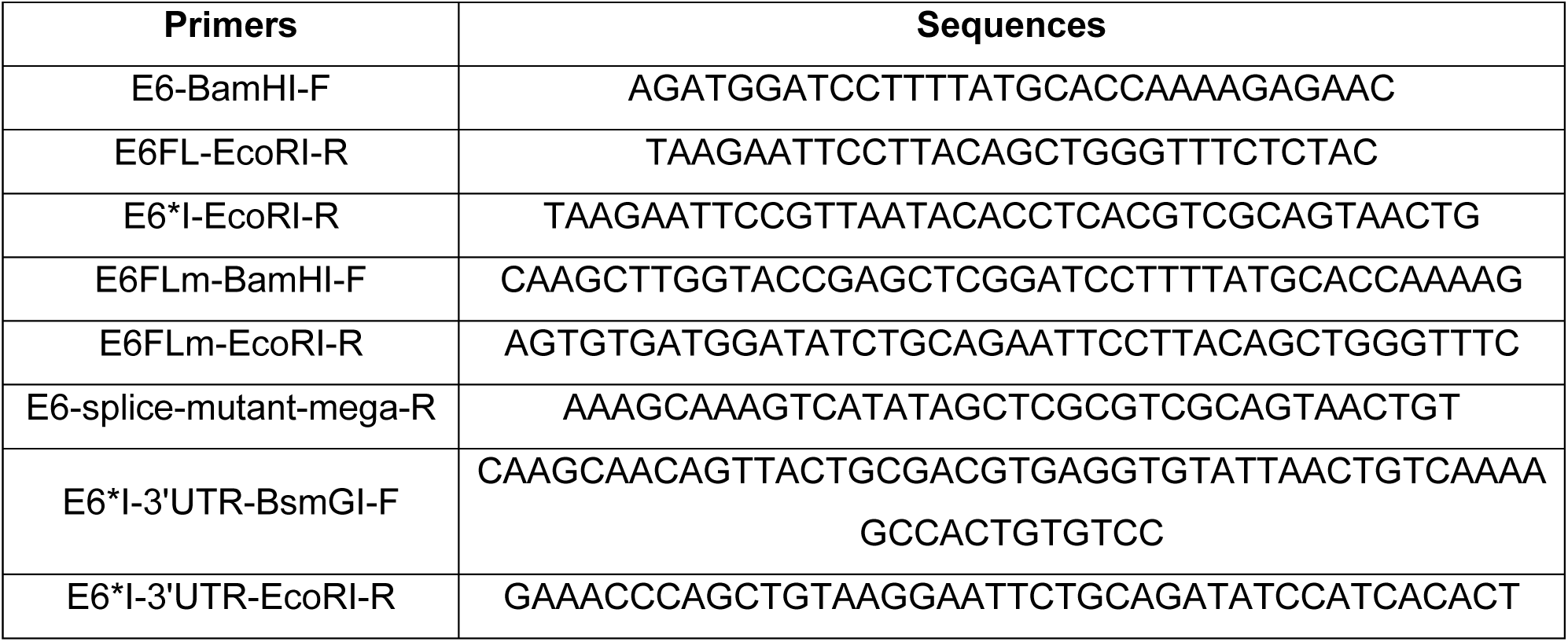
Primers for generation of plasmids.

**Table S5.**
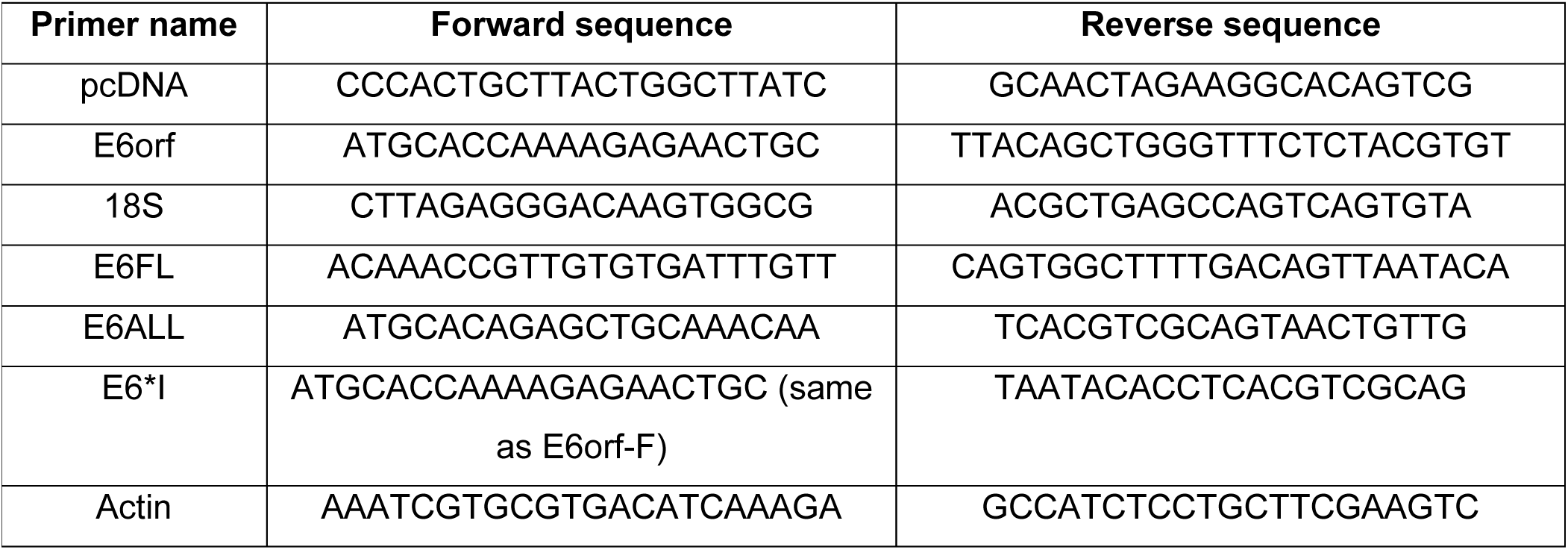
Primers for PCR and qPCR.

## Notes

### Competing Interest Statement

Patent applied based on ASO and biomarker

### Summary of Updates

New patient cohort for immunofluorescence staining New patient cohort for RNAsequencing Limit to only oropharyngeal cancer patients in survival analysis

