## Supplementary Figures and Tables for "Splicing of HPV16 E6 promotes aggressive invasion in oropharyngeal cancer via redistribution of E-cadherin"

### Supplementary Fig. S1

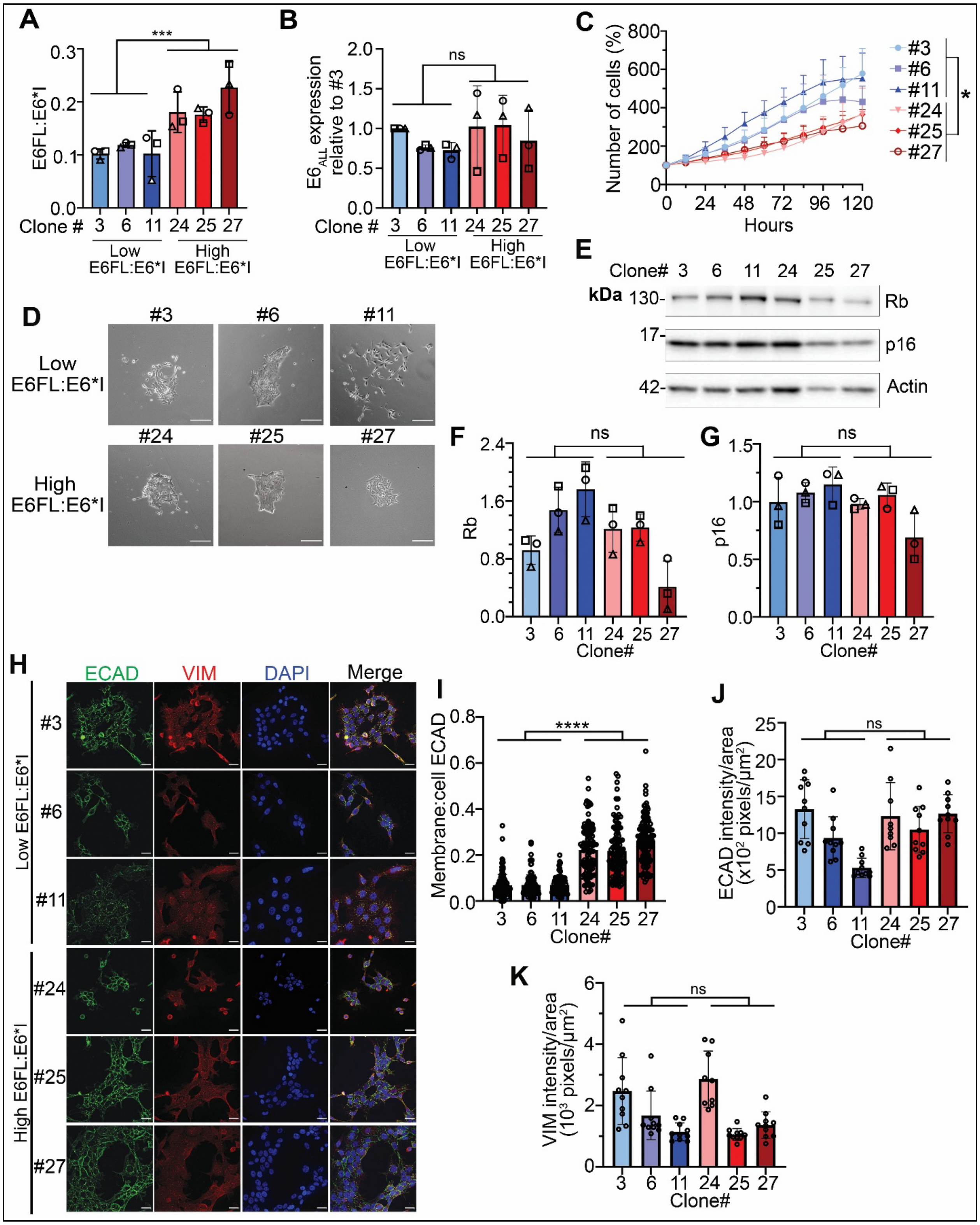

**Fig. S1: Validation and functional phenotypes of UPCI:SCC154 single-cell clones**

(A-B) Validation of E6FL:E6*I (A) and $E6_{ALL}$ (B) transcript expression in selected clones. $E6_{ALL}$ was normalized with 18S mRNA expression and expressed as a fold change relative to clone #3. (C) Proliferation assay performed on UPCI:SCC154 single-cell clones. Viable cells were quantified and expressed as percent (%) of 0h. Three independent experiments were quantified, with 4 replicates per experiment. (D) Phase contrast images of colonies after 15 days of growth. Scale bar = 500µm (E) Representative immunoblot showing Rb and p16 expression from three independent experiments. Actin was used as the loading control . (F&G) Densitometric quantification of Rb (F) and p16 (G), normalized to actin. Each shape shows one experiment (three). ns= not significant (unpaired t test). (H) Individual channels for immunofluorescence staining of DAPI (blue), E cadherin (ECAD, green) and vimentin (VIM, red) in UPCI:SCC154 single-cell clones. (I) Membrane:cell ECAD was quantified from (H). Each dot represents one cell. At least 120 cells were quantified. (J&K) ECAD (J) and VIM (K) was quantified per field and normalized by total cell area in the field. Each dot represents one field.

For all graphs, data are represented as mean ± SD. * $p<0.05$; ns= not significant (One-way ANOVA post-hoc Tukey)

Supplementary Fig. S2

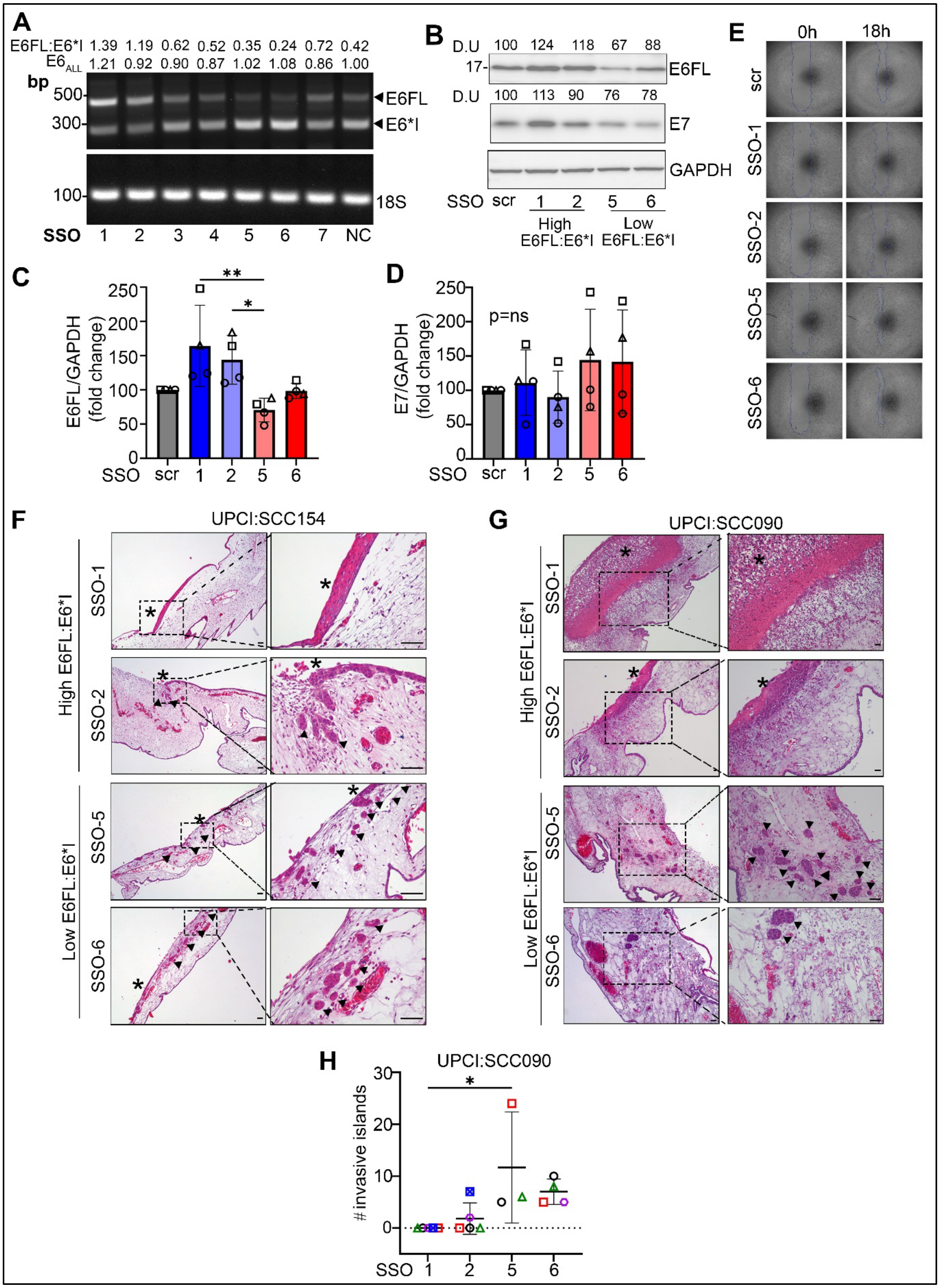

**Fig.S2: Functional studies performed in HPV+ OPSCC cell lines transfected with SSOs.**

(A) Representative agarose gel from 3 independent experiments validating expression of E6 isoforms in cDNA of UPCI:SCC090 transiently transfected with SSOs. NC represents non-targeting control, scr represents scrambled SSO control. E6FL:E6*I ratios were quantified by densitometry. $E6_{ALL}$ transcript expression was quantified by summing E6FL and E6*I and normalizing by 18S of the same sample, then expressed as a fold change relative to NC

(B) Representative immunoblot of E6FL and E7 in UPCI:SCC154 transfected with SSOs. GAPDH is used as a loading control

(C&D). Densitometry of E6FL (C) and E7 (D) of 4 independent experiments including (B)

(E) Wound healing images of UPCI:SCC154 transfected with SSOs. Quantification is in Fig.4E

(F-G) Representative H&E images from CAM assays performed in UPCI:SCC154 (F) and UPCI:SCC090 (G) transfected with SSOsArrows represent example of invasive islands that detach from the main tumor bulk. Asterisk represents the main tumor. In the images showing UPCI:SCC090 transfected with SSO-1 and SSO-2, no invasive island was identified. In images showing CAM from UPCI:SCC090 transfected with SSO-5 and SSO-6, the main tumor bulk could not be identified.

(H) Number of invasive islands detached from main tumor in CAM with UPCI:SCC090 injected with SSOs.

For all graphs, data are represented as mean ± SD. * $p<0.05$; ** $p<0.01$, ns= not significant (One-way ANOVA post-hoc Tukey)

**Supplementary Fig. S3**

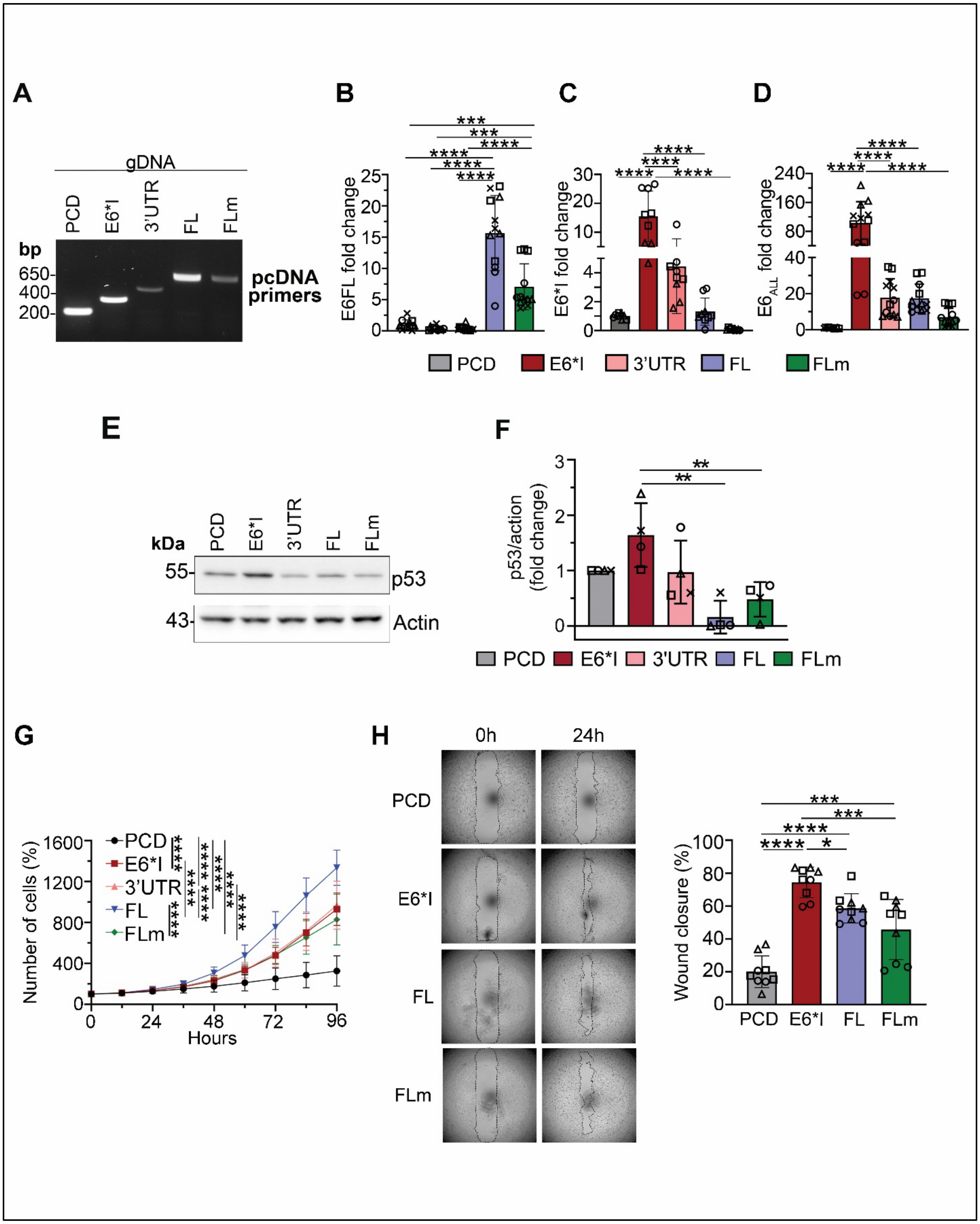

**Fig. S3: Overexpression of HPV16 E6 isoforms in HPV-negative OPSCC cell line, UM-SCC-38.**

(A) Agarose gel showing overexpression of E6 isoforms in genomic DNA with PCR using primers against pcDNA vector. Expected size: PCD= 198 bp; E6*I=356 bp; E6*I 3'UTR (3'UTR)=497 bp; FL/FLm=679 bp)

(B-D) RT-qPCR of cell lines using primers against full-length E6 (E6FL) (B), E6*I (C) and all E6 isoforms ($E6_{ALL}$) (D). Each shape represents an independent experiment, with three replicates per experiment. Transcript expression was represented as fold change relative to PCD control.

(E) Representative immunoblot for p53, the downstream target of E6, in lysates. Actin was used as a loading control.

(F) Densitometry for p53 was normalized to actin and expressed relative to PCD control from (C) and three other experiments (total four experiments). Each shape represents an independent experiment (four). Data is represented as mean ± SD.

(G) Proliferation assay performed on stable cells. Number of cells were quantified at indicated time points and normalized as a percent (%) to 0h.

(H) Representative images of wound healing assay of stable cells. Wound closure is represented as the percent of wound closure relative to 0h. Each shape is an independent experiment; same shapes represent replicates of the same experiment.

One-way ANOVA with post-hoc Tukey test was performed for all assays. **p<0.01, ***p<0.001, ****p<0.0001

Supplementary Fig. S4

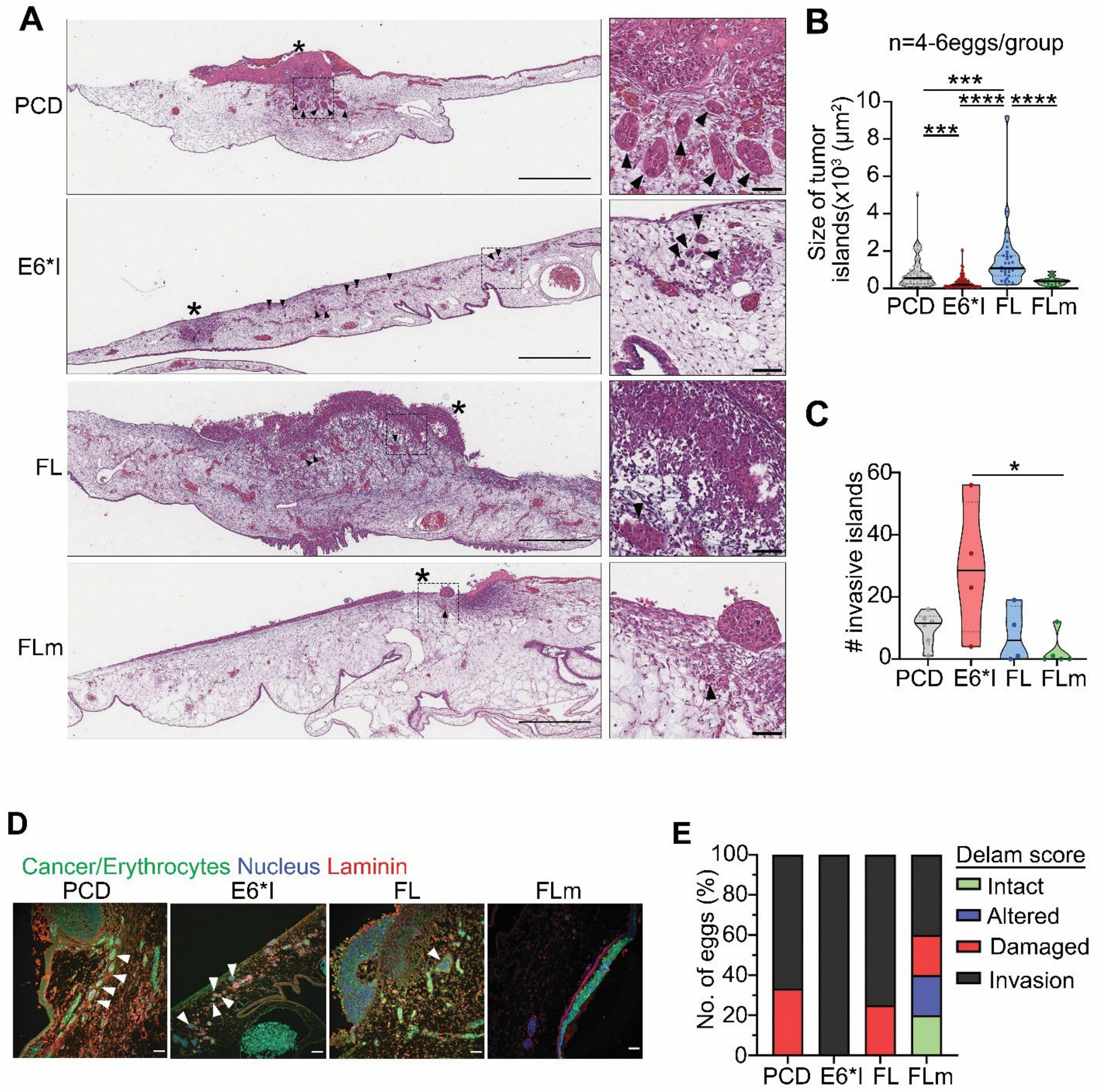

**Figure S4. CAM assay performed in UM-SCC-38 overexpressing E6 isoforms.**

(A) UM-SCC-38 labeled with CellTracker™ Green CMFDA Dye were seeded on the upper CAM that was harvested 3 days later, sectioned and stained with hematoxylin and eosin (H&E). Whole-slide imaging of CAM sections was performed. Invasive tumor islands are indicated by black arrowheads. Tumor bulk is indicated by an asterisk (*). Right panel shows enlarged images from boxed areas in left panels. Scale bar= 500μm (left), 50μm (right).

(B&C) Size of invasive islands (B) and number of invasive islands (C) were quantified from the entire CAM section shown in (A). N=4-6 eggs/group. Each point in (B) and (C) represents an invasive tumor island; each point in (D) represents a different egg. Data are represented as mean ± SD. *$p<0.05$, ***$p<0.001$, ****$p<0.0001$ (One-way ANOVA post-hoc Tukey test).

(D) Representative images of upper CAM stained for laminin (red; basement membrane) and DAPI (blue; nucleus). Green signals highlights fluorescently-labelled tumor cells with some non-specific staining observed in erythrocytes. Arrowheads indicate invasive tumor islands. Scale bar =50μm.

(E) Basement membrane degradation of immunofluorescence images including (D) was scored according to the CAM-Delam scoring system reported by Palaniappan et al. (2020)  (Fisher's Exact Score; p=ns).

**Supplementary Fig. S5**

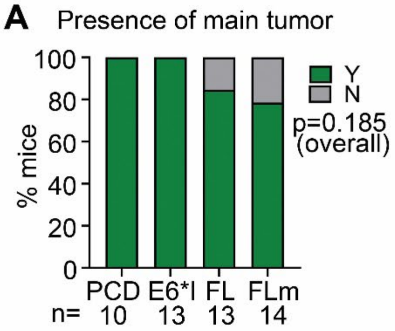


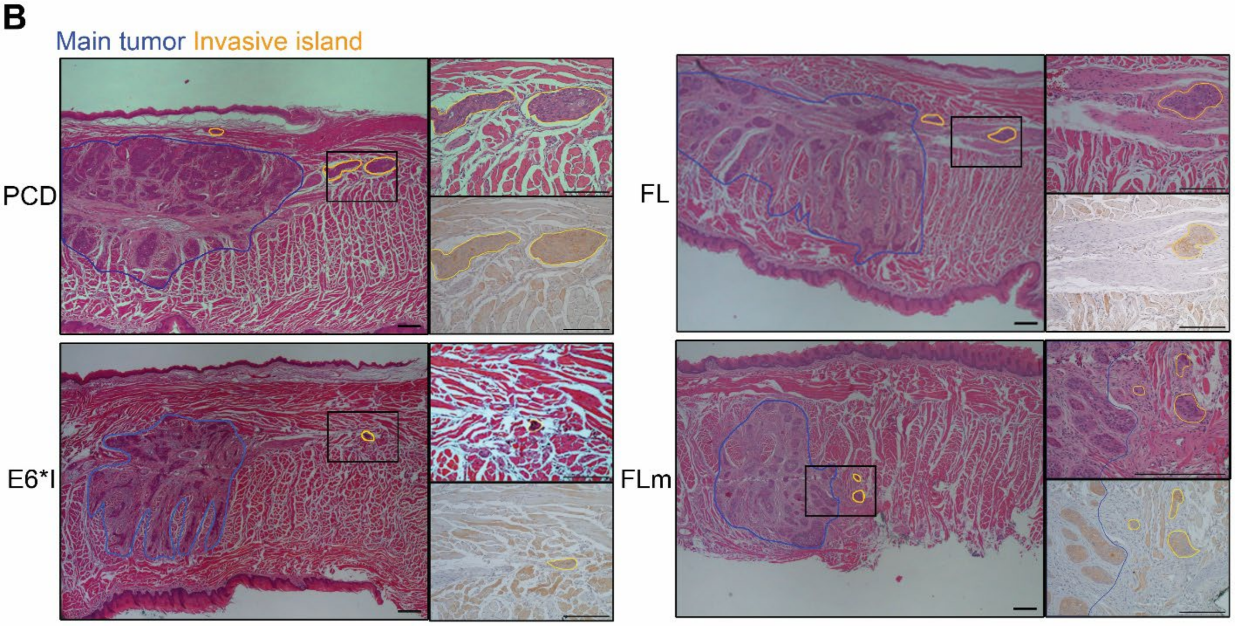

**Fig. S5: Mouse tumors.**

(A) Percentage (%) of mice showing presence of main tumor (Y=yes, N=no).

(B) H&E and CK-stained images of main tumor bulk (blue outline) and invasive islands (yellow outline) in mouse tongue. Right panels are high magnification H&E and CK-stained images of boxed area in left panels. Scale bar =200μm in both panels

**Supplementary Fig. S6**

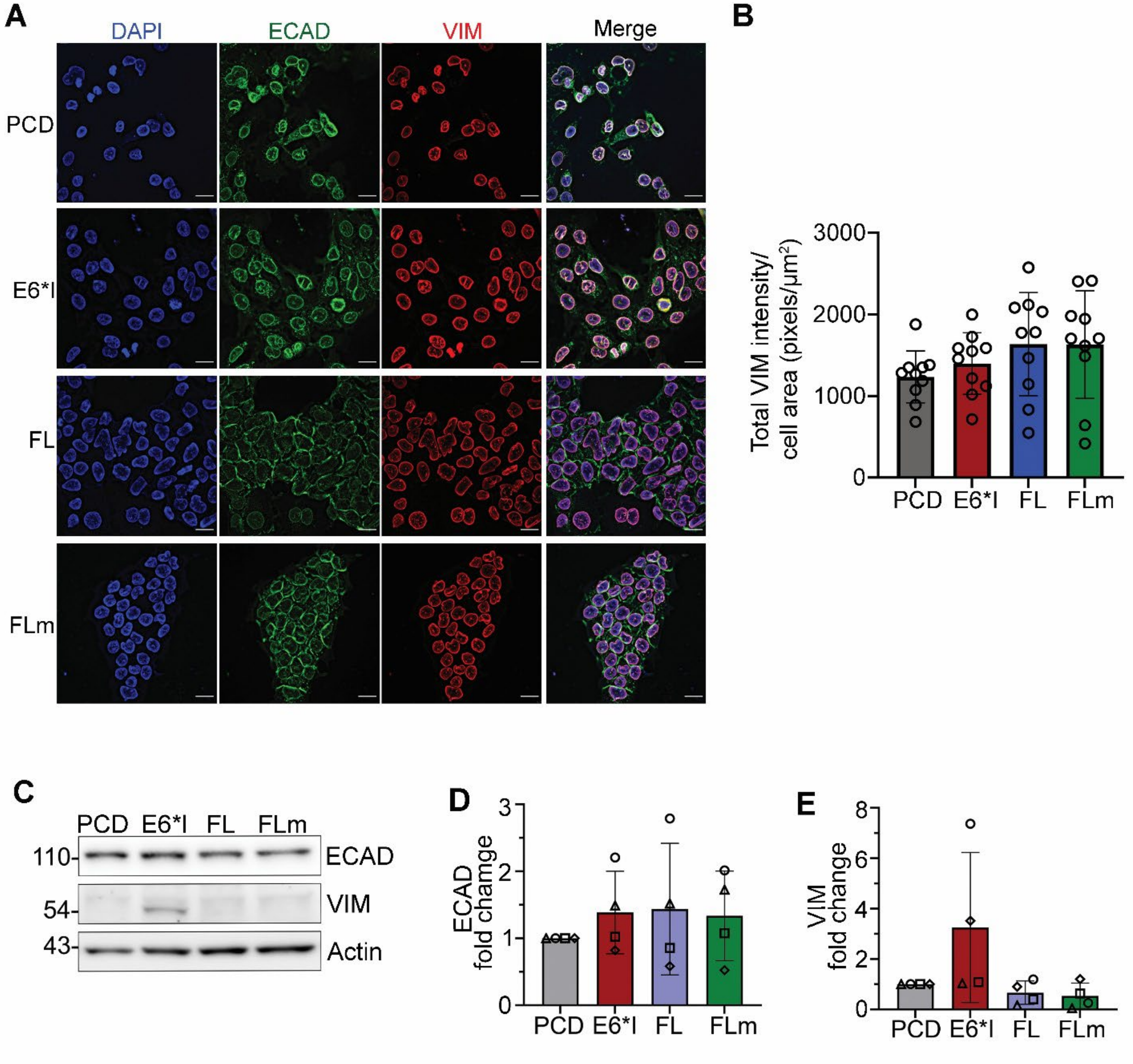

**Fig. S6: EMT markers in UM-SCC-38 overexpressing E6 isoforms.**

Individual channels of Immunofluorescence of DAPI (blue), E cadherin (ECAD, green) and vimentin (VIM, red) in UM-SCC-38 stably overexpressing E6 isoforms. Scale bar =100 μM.

Quantification of total VIM intensity normalized to cell area over 10 independent fields including (A) (one data point represents one field). One-way ANOVA post-hoc Tukey comparison; no statistical significance.

Representative Immunoblot of ECAD and VIM. Actin was a loading control.

(D&E) Densitometry of immunoblots for ECAD (D) and VIM (E) normalized to actin and expressed relative to PCD. Each data point is an independent experiment. For all graphs in this figure, data are represented as mean ± SD (one-way ANOVA post-hoc Tukey test; no asterisk=not significant)

**Supplementary Fig. S7**

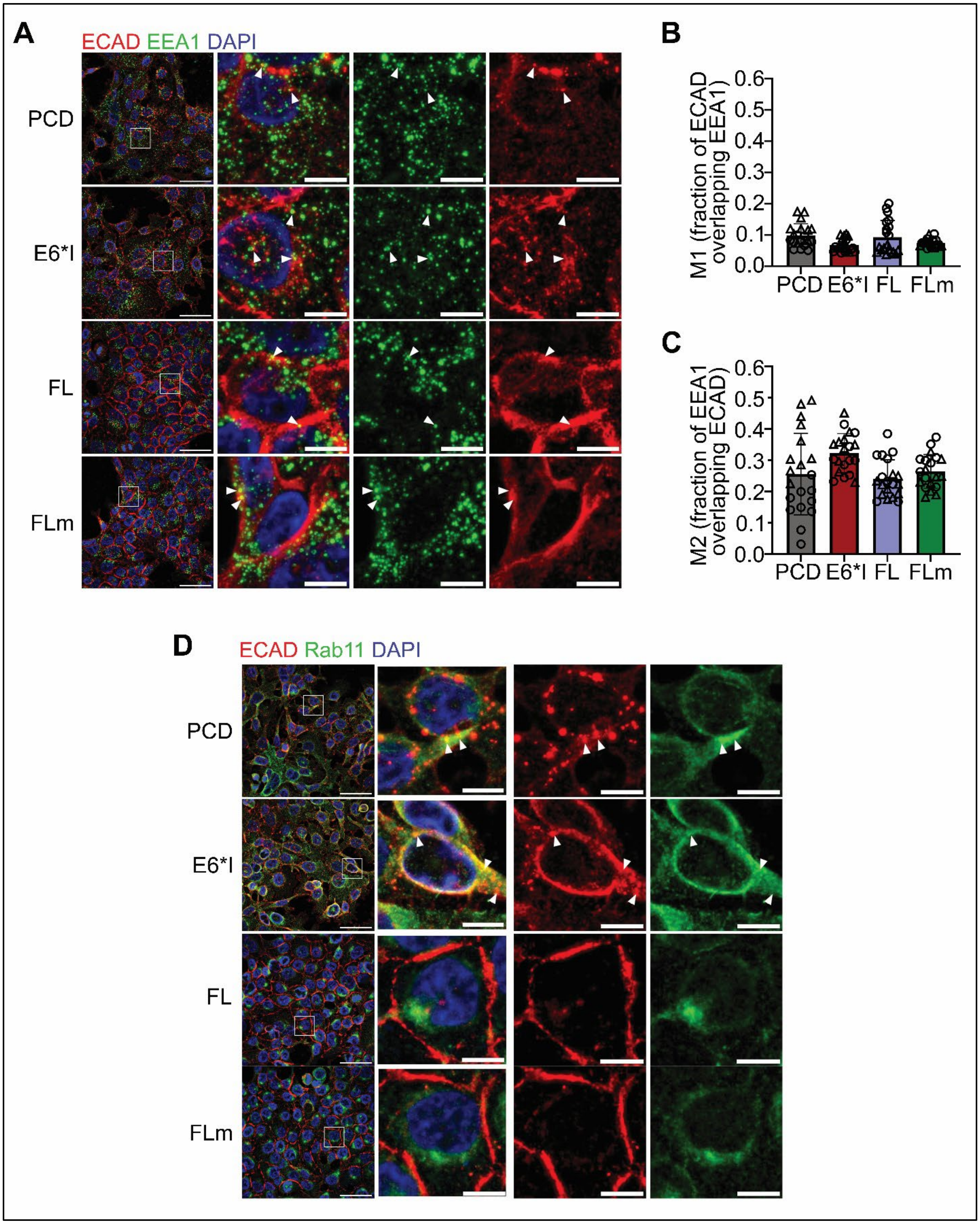

**Fig. S7: Co-localization of ECAD with markers for endocytosis vesicles, EEA1 and Rab11.**

(A) Maximum projection images of ECAD (red) and EEA1 (green) in UM-SCC-38 expressing E6 isoforms or vector control (PCD). Second panel shows merged image of boxed area in first panel. Third and fourth panels show individual channels for ECAD (red) and EEA1 (green), respectively. Arrowheads show co-localization of ECAD and EEA1. Scale bar= 50μm

(B&C) Co-localization of ECAD and EEA1 was quantified from 12 z slices, 5 fields/duplicate. Manders' coefficients, i.e. fraction of ECAD overlapping EEA1 (B) and fraction of EEA1 overlapping ECAD (C), were quantified using ImageJ JaCoP plugin. Each point represents a different field. Data are represented as mean ± SD. *$p<0.05$, one-way ANOVA post-hoc Tukey)

(D) Maximum projection images of ECAD (red) and Rab11 (green) in UM-SCC-38 with PCD or E6 isoforms. Second panel shows enlarged image of boxed area in first panel. Third and fourth panels are ECAD (red) and Rab11 (green), respectively. DAPI (blue) = nucleus. Scale bar = 50μm (left) and 10μm (right).

**Supplementary Fig. S8**

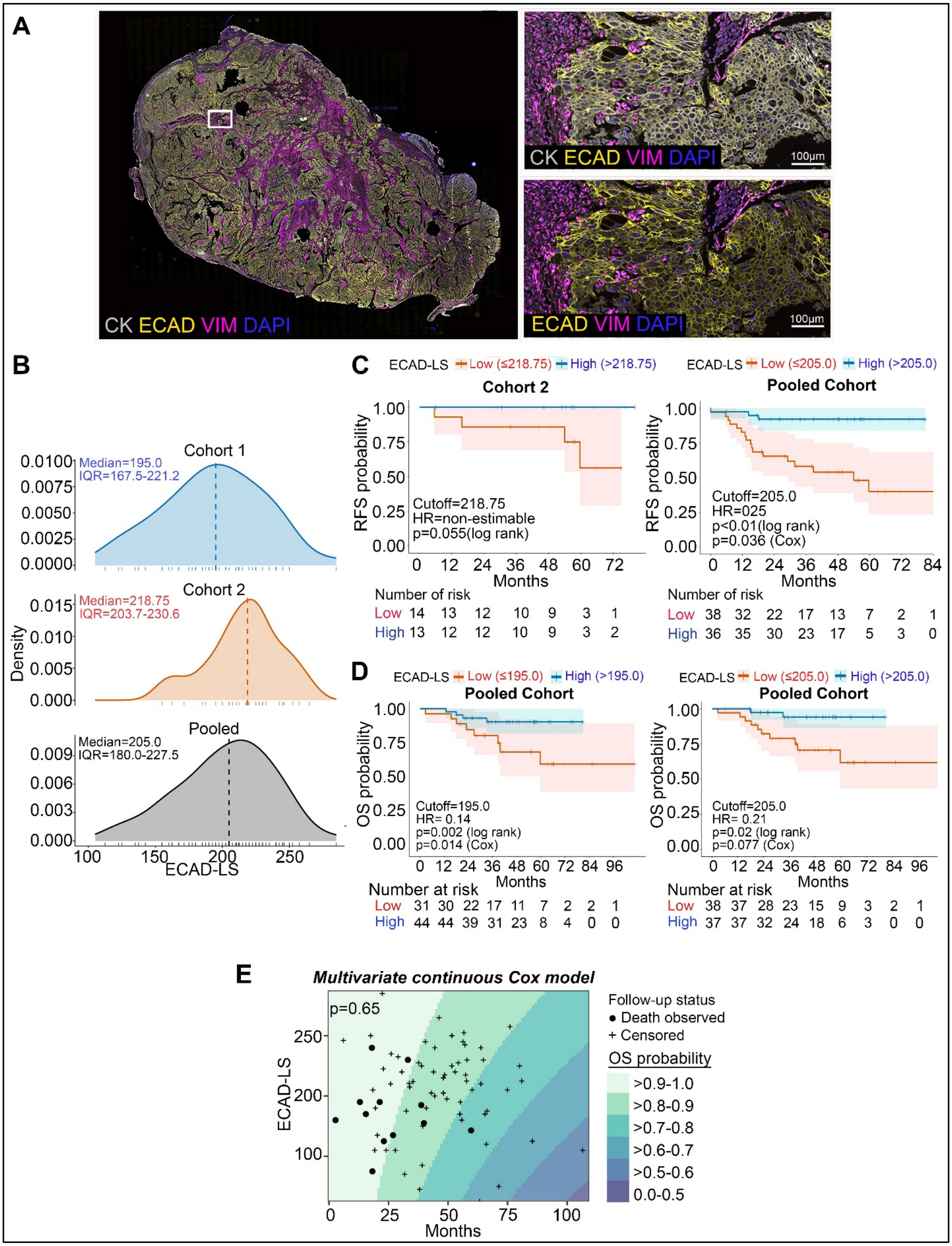

**Figure S8. ECAD localization and clinical outcomes in HPV+ OPSCC.**

(A) Representative whole-slide (left) and enlarged (right) multiplex immunofluorescence images showing CK (grey), ECAD (yellow), VIM (magenta) and DAPI (blue). Panels on left are enlarged areas from boxed area from right.

(B) Distributions of ECAD localization scores (ECAD-LS) in cohorts 1 and 2 and the pooled cohort; dashed lines indicate medians and tick marks in x axis of plots indicate individual tumors.

(C) Recurrence-free survival (RFS) in cohort 2 and the pooled cohort according to low versus high ECAD-LS, using the cohort 1 median (195.0) as the cutoff.

(D) Overall survival (OS) in the pooled cohort using the indicated ECAD-LS cutoffs. Left panel is stratified by cohort 1 median as a locked cutoff (195.0) and Right panel is stratified by pooled median (205.0). In D and E, shaded bands represent 95% confidence intervals, tick marks indicate censored observations, and numbers at risk are shown below the plots.

(E) Multivariable continuous Cox model of ECAD-LS and OS; background colors indicate predicted OS-probability intervals, filled circles indicate deaths, and plus signs indicate censored observations.

**Supplementary Fig. S9**

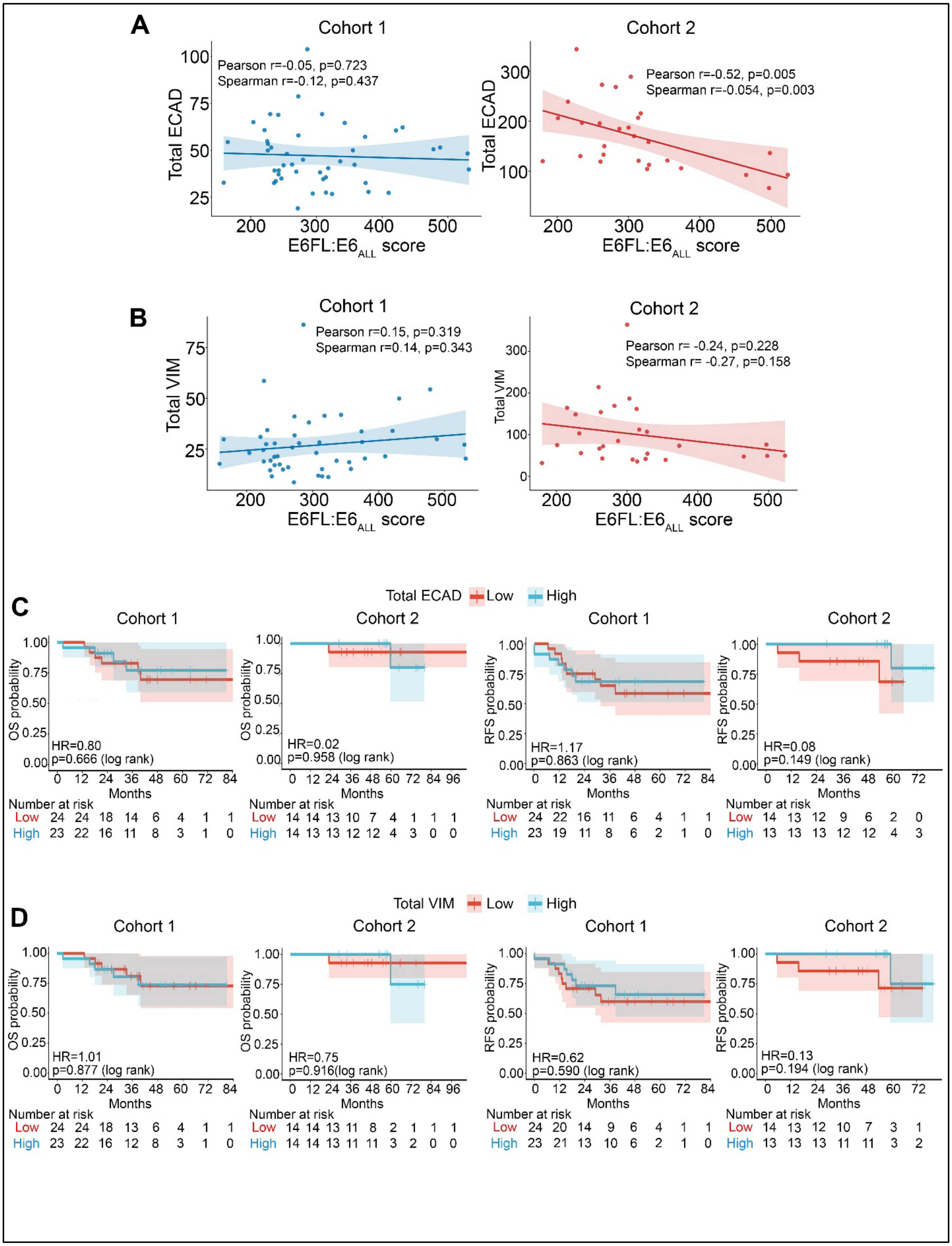

**Figure S9. Associations of total ECAD and VIM expression with E6FL:E6ALL scores and clinical outcomes.**

(A&B) Associations of the E6FL:E6ALL score with total ECAD (A) and total vimentin (B) in cohorts 1 and 2. Lines indicate linear regression fits, and shaded bands indicate 95% confidence intervals.

(C&D) Kaplan–Meier estimates of overall survival (C) and recurrence-free survival (D) according to low versus high total ECAD or VIM expression, dichotomized at the median within each cohort. Shaded bands indicate 95% confidence intervals, tick marks indicate censored observations, and numbers at risk are shown below each plot.

**Supplementary Fig. S10**

A

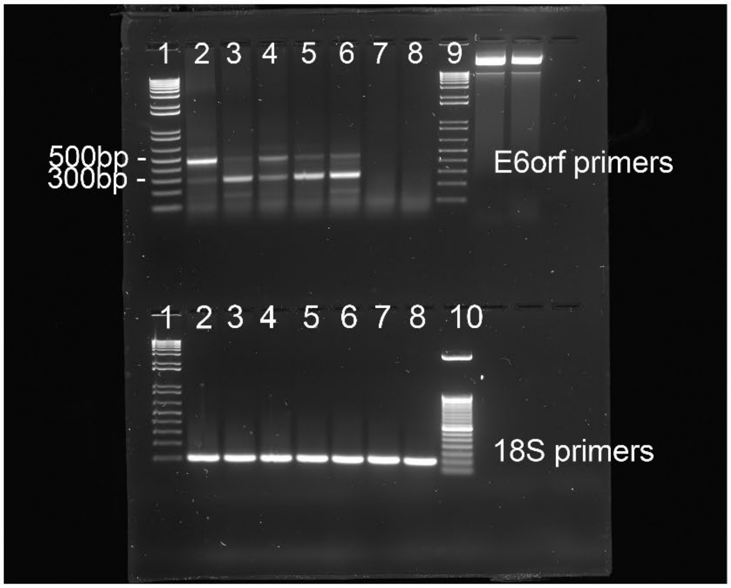


1- 1kb plus ladder
2- HOK16B
3- UM-SCC-47
4- UPCI:SCC090
5- UPCI:SCC154
6- UM-SCC-104
7- UM-SCC-38
8- UM-SCC-1
9- 100bp ladder
10- 50bp ladder

B

UM-SCC-38 gDNA

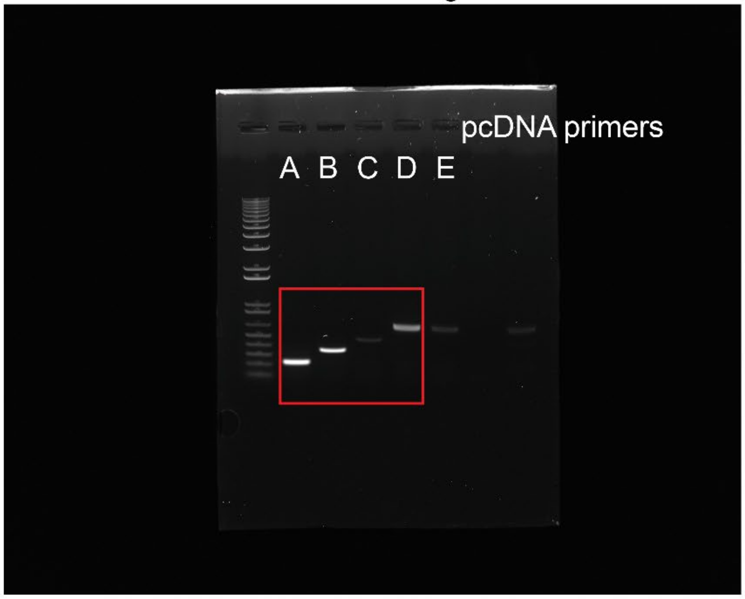


A: PCD
B: E6*I
C: E6*I 3'UTR
D: FL
E: FLm

**Fig. S10: Uncropped images of agarose gel for RT-PCR experiments.**

(A) Uncropped raw image for Fig. 1B. (B) Uncropped raw image for Fig. S3A. Red box annotates the bands used in this study.

**Supplementary Fig. S11**

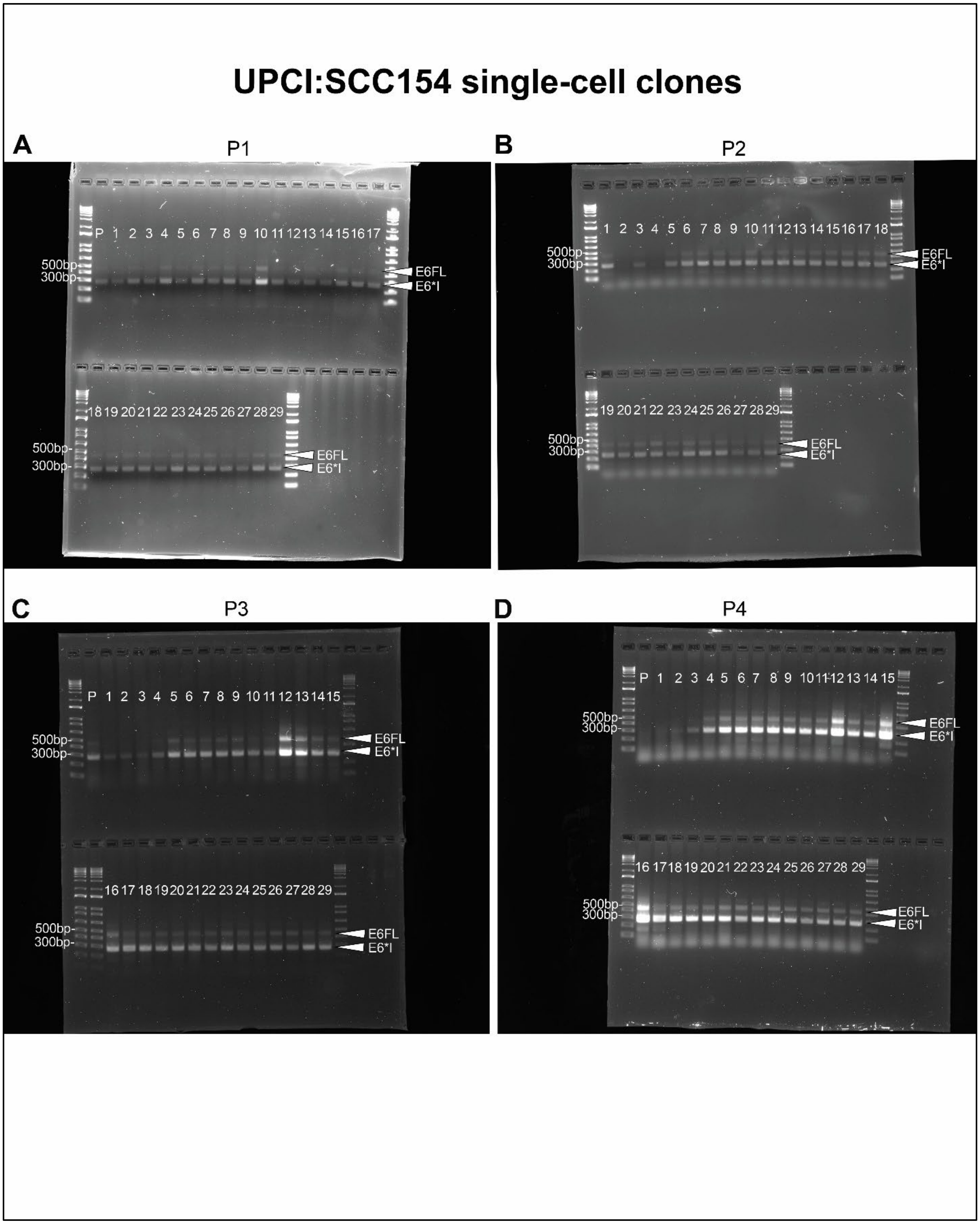

**Fig. S11: Raw data for screening of UPCI:SCC154SC clones in Fig. 2B.**

29 clones were isolated from single cells from UPCI:SCC154 parent cell line. RT-PCR was performed using E6orf primers and PCR products were electrophoresed on an agarose gel. E6FL:(E6FL+E6*I) ratio was expressed as densitometric units of E6FL divided by the sum of densitometric units of E6FL and E6*I. Four consecutive passages (P1 - P4) were screened.

P= UPCI:SCC154 parent cell line.

**Supplementary Fig. S12**

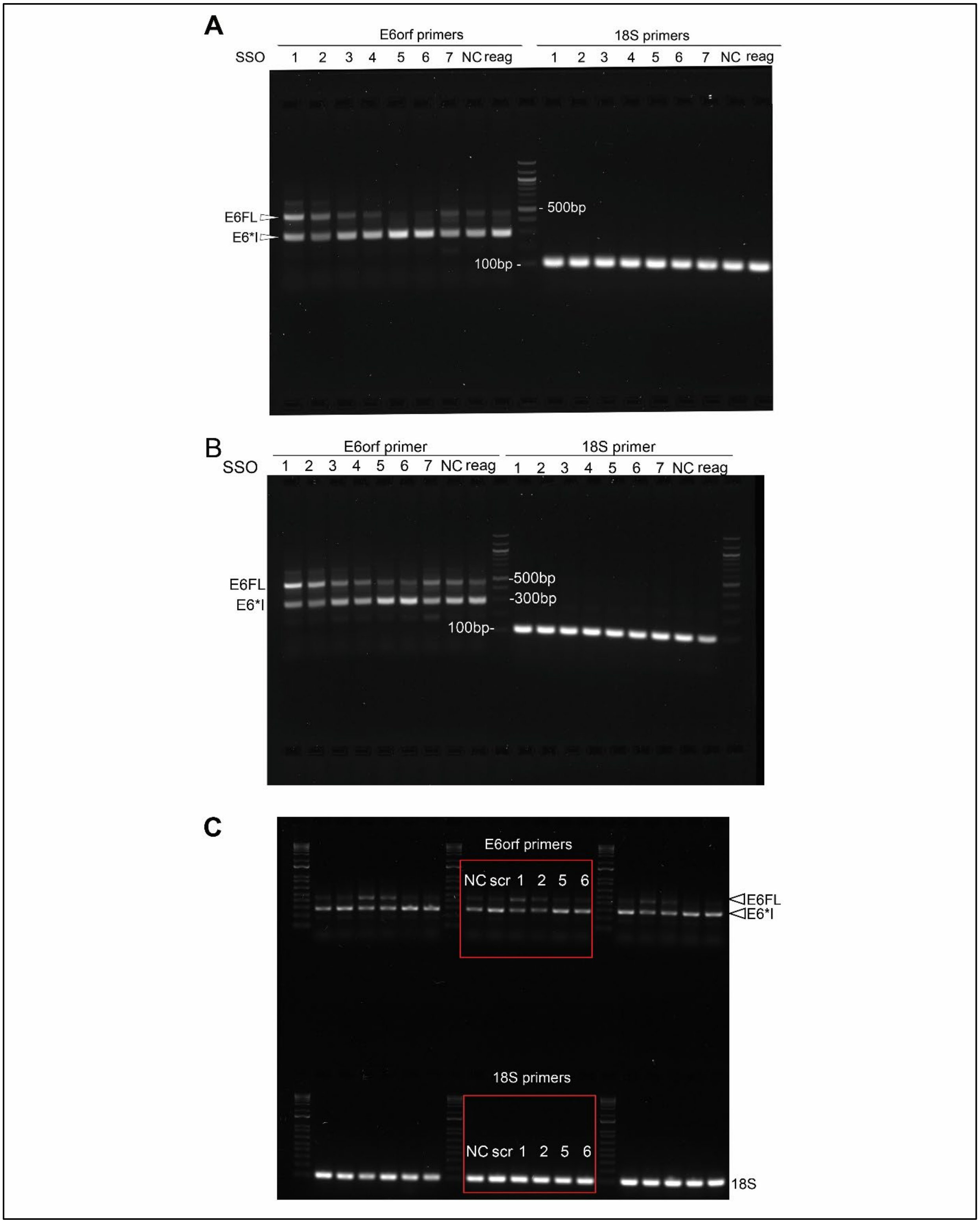

**Fig. S12: Raw data for screening of SSO experiments.**

(A) Raw data for Fig. 3B. (B) Fig. S2A (C) Fig. 3D

Supplementary Fig. S13

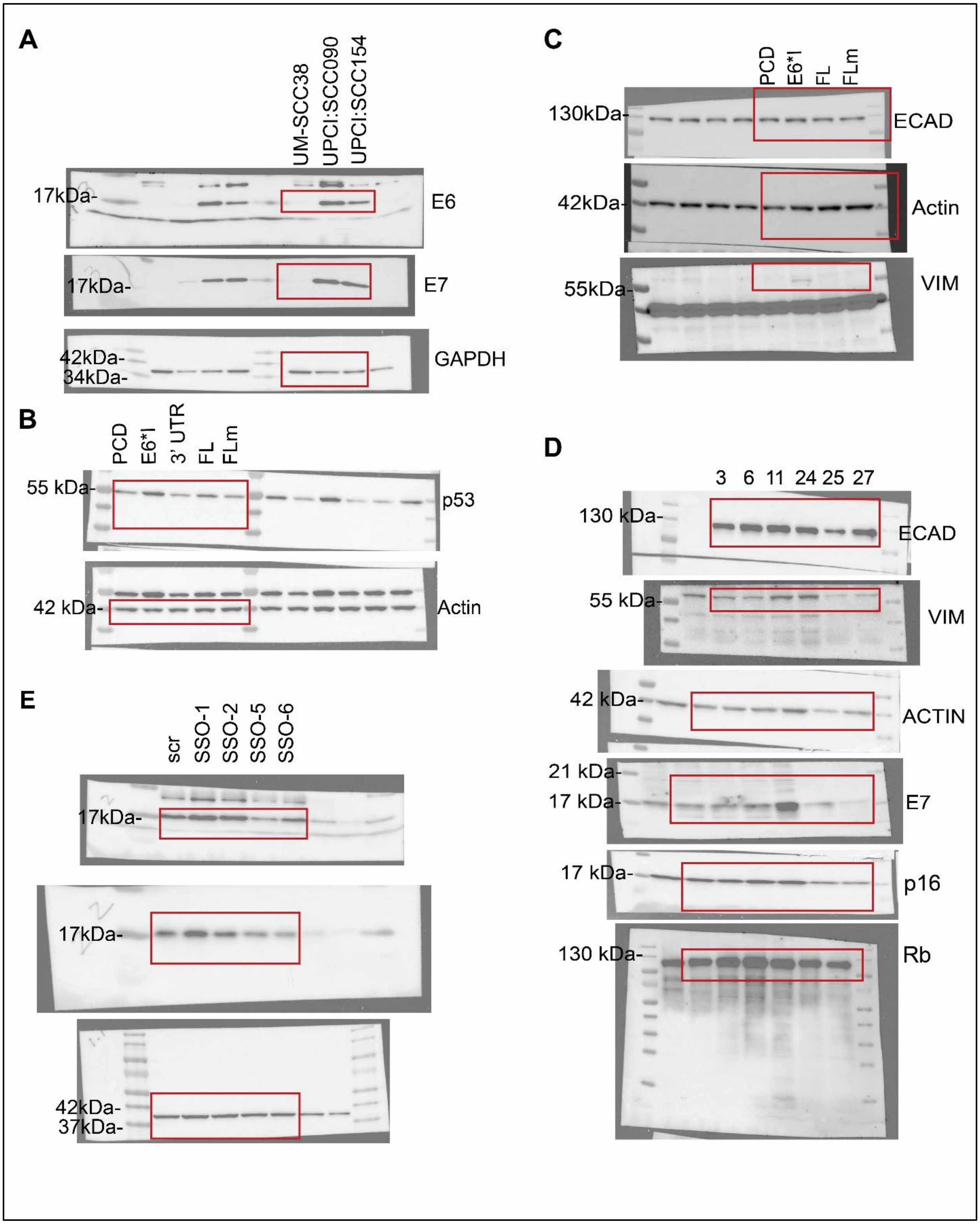

**Fig. S13: Raw immunoblots used in study.**

Raw data for (A) Fig. 1J; (B) Fig. S2E; (C) Fig. S6C; (D) Fig. S1E; (E) Fig. S2B

**Table S1.**
SSOs and sequences

| SSO name | Sequence |
|---|---|
| SSO-1 | TACCTCACGTCGCAGTAACT |
| SSO-2 | AGTCATATACCTCACGTCGC |
| SSO-3 | TATACTATGCATAAATCCCGAA |
| SSO-4 | TCTCTATATACTATGCATAAAT |
| SSO-5 | AATGTCTATACTCACTAATTTT |
| SSO-6 | TAATGTTGTTCCATACAAACTA |
| SSO-7 | TAATACACCTAATTAACAAATC |
| Negative control | CCTATAGGACTATCCAGGAA |
| Scrambled SSO | GTCATACCCGACGCTTCATA |

**Table S2**

Clinical summary for multiplexed immunofluorescence cohort

| Characteristic | Cohort 1 n = 47 | Cohort 2 n= 28 | Pooled, n = 75 |
|---|---|---|---|
| **RNAseq cohort** | | | |
| UM20 | 0 (0.0%) | 20 (71.4%) | 20 (26.7%) |
| UM18 | 9 (19.1%) | 3 (10.7%) | 12 (16.0%) |
| UM67 | 38 (80.9%) | 5 (17.9%) | 43 (57.3%) |
| **ECAD-LS, median (IQR)** | 195.0 (167.5–222.5) | 218.8 (202.5–231.3) | 205.0 (180.0–227.5) |
| **Age, years, median (range)** | 58.0 (43.0–76.0) | 58.0 (43.0–84.0) | 58.0 (43.0–84.0) |
| **Sex** | | | |
| Female | 7 (14.9%) | 3 (10.7%) | 10 (13.3%) |
| Male | 40 (85.1%) | 25 (89.3%) | 65 (86.7%) |
| **Race** | | | |
| Black | 0 (0.0%) | 1 (3.6%) | 1 (1.6%) |
| Non-hispanic White | 37 (78.7%) | 24 (85.7%) | 61 (81.3%) |
| North American native | 1 (2.1%) | 0 (0%) | 1 (1.3%) |
| Not determined | 9 (19.1%) | 3 (10.7%) | 12 (16.0%) |
| **Smoking status** | | | |
| Ever smoker | 28 (62.2%) | 3 (37.5%) | 31 (58.5%) |
| Never smoker | 17 (37.8%) | 5 (62.5%) | 22 (41.5%) |
| Not determined | 2 | 20 | 22 |
| **Primary tumor site** | | | |
| Oropharynx | 47 (100.0%) | 28 (100.0%) | 75 (100.0%) |
| **Overall AJCC stage** | | | |
| Stage I | 3 (6.4%) | 8 (29.6%) | 11 (14.9%) |
| Stage II | 32 (68.1%) | 13 (48.1%) | 45 (60.8%) |
| Stage III | 10 (21.3%) | 6 (22.2%) | 18 (24.3%) |
| Stage IV | 2 (4.26%) | 0 (0%) | 2(2.67%) |
| Unknown | 0(0%) | 1 (3.6%) | 1(1.33%) |
| **Clinical T stage** | | | |
| T1 | 12 (25.5%) | 11 (40.7%) | 23 (31.1%) |

| | | | |
|---|---|---|---|
| T2 | 20 (42.6%) | 7 (25.9%) | 27 (36.5%) |
| T3 | 6 (12.8%) | 4 (14.8%) | 10 (13.5%) |
| T4 | 9 (19.1%) | 5 (18.5%) | 14 (18.9%) |
| Missing | 0(0%) | 1(3.6%) | 1(1.33%) |
| **Clinical N stage** | | | |
| N0 | 2 (4.7%) | 3 (11.1%) | 5 (7.1%) |
| N1 | 4 (9.3%) | 6 (22.2%) | 10 (14.3%) |
| N2 | 32 (74.4%) | 17 (63.0%) | 49 (70.0%) |
| N3 | 5 (11.6%) | 1 (3.7%) | 6 (8.6%) |
| Missing | 4(8.51%) | 1(3.6%) | 5(6.67%) |
| **Clinical M stage** | | | |
| M0 | 47 (100.0%) | 28 (100.0%) | 75 (100.0%) |
| **HPV integration status** | | | |
| Episomal | 29 (61.7%) | 15 (53.6%) | 44 (58.7%) |
| Integrated | 18 (38.3%) | 7 (25.0%) | 25 (33.3%) |
| Not determined | 0 (0.0%) | 6 (21.4%) | 6 (8.0%) |
| **IMU/KRT subtype** | | | |
| IMU | 17 (36.2%) | 13 (46.4%) | 30 (40.0%) |
| KRT | 30 (63.8%) | 15 (53.6%) | 45 (60.0%) |
| **Death events** | | | |
| No | 37 (78.7%) | 26 (92.9%) | 63 (84.0%) |
| Yes | 10 (21.3%) | 2 (7.1%) | 12 (16.0%) |
| **Recurrence** | | | |
| No | 35 (74.5%) | 24 (89.3%) | 60 (80.0%) |
| Yes | 12 (25.5%) | 3 (10.7%) | 15 (20.0%) |
| Missing | 0 | 1 | 1 |

**Table S3**

Cutoff and number of events for overall survival (OS) and recurrence-free survival (RFS) analysis for multiplexed immunofluorescence.

| Analysis group | Outcome | ECAD-LS Cutoff | Total, n (events) | Low ECAD-LS, n (events) | High ECAD-LS, n (events) |
|---|---|---|---|---|---|
| Cohort 1 | OS | 195 | 47 (10) | 25 (8) | 23 (2) |
| Cohort 2 | OS | 195 | 28 (2) | 6 (2) | 22 (0) |
| Cohort 2 | OS | 218.75 | 28 (3) | 14 (3) | 13 (0) |
| Pooled | OS | 195 | 75 (12) | 31 (10) | 44 (2) |
| Pooled | OS | 205 | 75 (12) | 38 (10) | 37 (2) |
| Cohort 1 | RFS | 195 | 47 (16) | 25 (13) | 22 (3) |
| Cohort 2 | RFS | 195 | 27 (3) | 6 (3) | 21 (0) |
| Cohort 2 | RFS | 218.75 | 27 (3) | 14 (3) | 13 (0) |
| Pooled | RFS | 195 | 74 (20) | 31 (17) | 43 (3) |
| Pooled | RFS | 205 | 74 (20) | 38 (17) | 36 (3) |

**Table S4**

Primers for generation of plasmids

| Primers | Sequences |
|---|---|
| E6-BamHI-F | AGATGGATCCTTTTATGCACCAAAAGAGAAC |
| E6FL-EcoRI-R | TAAGAATTCCTTACAGCTGGGTTTCTCTAC |
| E6*I-EcoRI-R | TAAGAATTCCGTTAATACACCTCACGTCGCAGTAACTG |
| E6FLm-BamHI-F | CAAGCTTGGTACCGAGCTCGGATCCTTTTATGCACCAAAAG |
| E6FLm-EcoRI-R | AGTGTGATGGATATCTGCAGAATTCCTTACAGCTGGGTTTC |
| E6-splice-mutant-mega-R | AAAGCAAAGTCATATAGCTCGCGTCGCAGTAACTGT |
| E6*I-3'UTR-BsmGI-F | CAAGCAACAGTTACTGCGACGTGAGGTGTATTAACTGTCAAAA GCCACTGTGTCC |
| E6*I-3'UTR-EcoRI-R | GAAACCCAGCTGTAAGGAATTCTGCAGATATCCATCACACT |

**Table S5**

Primers for PCR and qPCR

| Primer name | Forward sequence | Reverse sequence |
|---|---|---|
| pcDNA | CCCACTGCTTACTGGCTTATC | GCAACTAGAAGGCACAGTCG |
| E6orf | ATGCACCAAAAGAGAACTGC | TTACAGCTGGGTTTCTCTACGTGT |
| 18S | CTTAGAGGGACAAGTGGCG | ACGCTGAGCCAGTCAGTGTA |
| E6FL | ACAAACCGTTGTGTGATTTGTT | CAGTGGCTTTTGACAGTTAATACA |
| E6ALL | ATGCACAGAGCTGCAAACAA | TCACGTCGCAGTAACTGTTG |
| E6*I | ATGCACCAAAAGAGAACTGC (same as E6orf-F) | TAATACACCTCACGTCGCAG |
| Actin | AAATCGTGCGTGACATCAAAGA | GCCATCTCCTGCTTCGAAGTC |
